# Integrative reappraisal of the Amazonian nurse frog *Allobates gasconi* (Morales 2002) based on topotypical data, with implications for the systematics and taxonomy of a large species complex

**DOI:** 10.1101/2023.07.10.548442

**Authors:** Anthony S. Ferreira, Miquéias Ferrão, Antonio S. Cunha-Machado, William E. Magnusson, James Hanken, Albertina P. Lima

**Affiliations:** Coordenação de Pesquisas em Biodiversidade, Instituto Nacional de Pesquisas da Amazônia, Manaus, Amazonas, Brazil; Pós-Doutorado Júnior – PDJ, Instituto Nacional de Pesquisas da Amazônia, Manaus, Amazonas, Brazil; Museum of Comparative Zoology, Harvard University, Cambridge, Massachusetts, United States of America; Laboratório de Herpetologia e Comportamento Animal, Departamento de Ecologia, Instituto de Ciências Biológicas, Universidade Federal de Goiás, Goiânia, Goiás, Brazil

**Keywords:** Bioacoustics, Integrative Taxonomy, Juruá River, Morphology, Tadpoles, *Várzea*

## Abstract

Taxonomic uncertainty at the species level compromises our knowledge of biodiversity, conservation and systematics. The impact of such uncertainty is heightened in megadiverse regions such as Amazonia due to high levels of cryptic diversity. We used integrative taxonomy based on newly collected topotypical specimens to redescribe the Amazonian nurse frog *Allobates gasconi* and infer its phylogenetic relationships. This species was described in 2002 based solely on morphology, but several characters crucial for the reliable diagnosis of species in *Allobates* were not considered. Our results show that *A. gasconi* sensu stricto is not a member of the *A. caeruleodactylus* clade as previously claimed, but is a member of the *A. trilineatus* clade. *Allobates gasconi* is readily distinguished from congeners by a combination of morphological and bioacoustic characters; a revised diagnosis is provided. The type series of *A. gasconi* comprises more than one species, and we exclude a paratype from lower Juruá River. The species is restricted geographically to flooded environments in the middle and upper Juruá River in Brazil and in the Ucayali River in Peru. The initial misidentification, subsequent absence of topotypic molecular and acoustic data, and the poor preservation condition of the type series have contributed to taxonomic confusion since *A. gasconi* was first described. The descriptions of other species of *Allobates* published more than two decades ago were based mainly on gross morphology and we recommend integrative taxonomic revisions to elucidate their systematics.

## Introduction

The taxonomy and systematics of many tropical anuran species described before the widespread utilization of DNA sequencing represent an impediment to describe of new species, especially for taxa belong to species complexes in highly biodiverse regions, such as Amazonia. Genetic data obtained from historic fluid-preserved type specimens has resolved taxonomic and systematic issues of some species that are rare or known only from their type series (e.g. Lyra et al., 2020; Rancilhac et al., 2020). However, DNA extraction and sequencing of formalin-fixed tissues is still challenging. On the other hand, revisiting topotypic localities to collect fresh material is a feasible alternative for those species with known and accessible type localities (e.g. Lima et al., 2009; Ferrão et al., 2020; Köhler et al., 2022). This approach also presents opportunities to collect additional data to integrate with molecular evidence, such as advertisement calls and larval morphology; two valuable sources of diagnostic characters used in contemporary anuran taxonomy.

Despite the large number of recently described species and advances in the systematics of nurse frogs of the genus *Allobates* Zimmermann and Zimmermann, 1988, several taxonomic uncertainties persist due to gaps left by insufficiently detailed or otherwise deficient descriptions (Jaramillo et al., 2021). One such problem concerns the identity of *A. gasconi* (Morales, 2002), a cryptically colored nurse frog originally described as *Colostethus gasconi* based on specimens collected in 1991 by Claude Gascon during an expedition along the Juruá River, western Brazilian Amazonia. The original description includes a diagnosis with 13 characters, a description of the holotype based on seven morphometric measurements, a brief account of its coloration in preservative, and a stylized drawing (Morales, 2002, pp. 30–32); it does not consider several characters that are now known to be very important to distinguish species of *Allobates*. Currently, the holotype is poorly preserved (dehydrated and with a muted coloration), 74% of the type series are juveniles and only some of the latter are in good condition. Moreover, the advertisement calls and tadpoles of *A. gasconi* are undocumented.

Since its original use, the name *Allobates gasconi* has been applied to specimens from many localities in Brazilian Amazonia (Grant et al., 2006; Lima et al., 2014; Melo-Sampaio et al., 2018; Réjaud et al., 2020; Vacher et al., 2020). Grant et al. (2006) inferred the phylogenetic relationships of *A. gasconi* for the first time, based on individuals from the Ituxí River (Amazonas, Brazil). Subsequently, Lima et al. (2014) sequenced specimens attributed to *A. gasconi* from a locality in the middle Juruá River ∼160 km from the type locality. Melo-Sampaio et al. (2018) compiled these data, sequenced additional specimens from southwestern Amazonia and reexamined the morphology of the type series. They noted differences in coloration and finger morphology between the type series and specimens recently collected by them, attributing this to population-level variation (Melo-Sampaio et al., 2018). Later, Vacher et al. (2020) sequenced specimens of *A. gasconi* from the Madeira River (Rondônia, Brazil) for DNA barcoding and concluded that populations from Juruá, Ituxí and Madeira rivers represent three distinct taxa. Finally, *A. gasconi* was inferred as sister to the *Allobates tapajos* species complex and a member of the *Allobates caeruleodactylus* clade by Réjaud et al. (2020).

At present, it is unclear whether any of the above-mentioned lineages represents *Allobates gasconi* sensu stricto. In view of the likelihood that the *A. gasconi* species complex represents an instance of cryptic diversity, robust diagnostic characters are needed to bolster further studies of its taxonomy and systematics. To that end, we collected specimens from the type locality and three paratype localities of *A. gasconi*. Based on this material, we present a reappraisal of *A. gasconi* sensu stricto through integrative taxonomy, infer its phylogenetic relationships, and assess if any of the previously reported lineages includes the species from the topotypic locality.

## Material and methods

### Sampling

Fieldwork was carried out in January 2021 and March 2022 along the middle and lower courses of the Juruá River, western Brazilian Amazonia (Fig. 1). Holotype and paratype localities were found by using approximate georeferenced coordinates available in Gascon et al. (2000), original ledger entry notes of the INPA-H collection and with assistance of local field guides. Adults were collected at the type locality (Jainú along the west bank of Juruá River, hereafter *Jainú West* [06°27’41” S, 68°44’18” W]) and three paratype localities along the east bank of the Juruá River (*Jainú East* [06°27’57” S, 68°44’35” W]; *Altamira* [06°34’35” S, 68°52’31” W] and *Vai-Quem-Quer* [03°18’38” S, 65°59’47” W]). Three lots of tadpoles were collected near adult males in *Jainú West* (*n* = 2) and *Altamira* (*n* = 1) and raised for morphological description in the laboratory until they reached Gosner stages 34–41. Tadpole identifications were subsequently confirmed through molecular analysis. The paratype localities Condor and Nova Vida were not sampled.

**Fig. 1.**
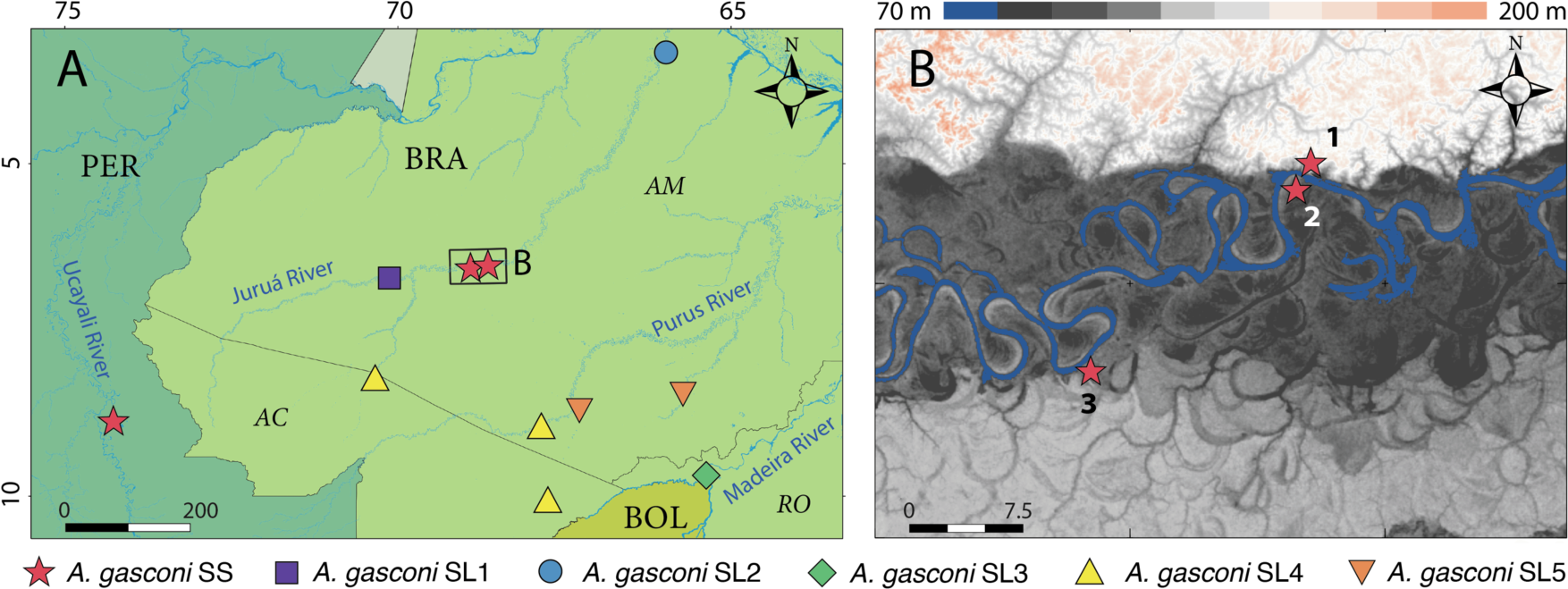
Geographic distribution of *Allobates gasconi* sensu lato (A) and sensu stricto (B). In the panel (B) numbers correspond to the type (1, *Jainú West*) and paratype (2, *Jainú East*; 3, *Altamira*) localities of *Allobates gasconi*. Dark grey areas in panel (B) are subject to flooding regimes (*várzea*), while other areas are not (*terra firme*). Abbreviations: SL, sensu lato; SS, sensu stricto; N, north. Countries: BOL, Bolivia, PER, Peru. Brazilian states: AC, Acre; AM, Amazonas; RO, Rondônia.

Adults and tadpoles were killed with a topical solution of 2% aqueous benzocaine. Tissue samples were taken from every adult and one tadpole from each clutch and preserved in 96% ethanol. Adults were fixed in 10% formalin and permanently stored in 70% ethanol; tadpoles were preserved and stored in 5% formalin. Protocols for collection and animal care follow the Brazilian Federal Council for Biology resolution number 148/2012 (Conselho Federal de Biologia – CFBio, 2012). Specimens were collected under a permit to A. P. Lima (number 1337) issued by the Instituto Brasileiro do Meio Ambiente e dos Recursos Naturais Renováveis – IBAMA (Ministry of Environment, Government of Brazil). Specimens are deposited in the herpetology collection of the Instituto Nacional de Pesquisas da Amazônia – **INPA-H** (Manaus, Brazil) and the Museum Paraense Emílio Goeldi (**MPEG** 44618–44629), Belém, Brazil.

Advertisement calls of 14 males were recorded in *Jainú West* (n = 5), *Jainú East* (n = 6) and *Altamira* (n = 3). Active calling males were recorded for 3 min with an AKG 568 EB directional microphone connected to a Marantz PMD660 or SONY PCM-D50 Digital Recorder with built-in microphones. Recordings were stored in uncompressed WAV files at a sampling frequency of 44.1 kHz and resolution of 16 bits. Microphones were positioned approximately 1 m from each calling male. Snout-vent length of recorded males ranged from 15.4 to 17.1 mm, and air temperature at the moment of recording ranged from 24.4 to 26.8°C. Recordings are deposited in the acoustic repository Fonoteca Neotropical Jacques Vielliard (**FNJV** 58775–58788), University of Campinas, Campinas, Brazil.

### DNA sequencing

Total genomic DNA was extracted from tissue samples of 19 specimens of *Allobates gasconi* (*Jainú West* = 4; *Jainú East* = 5, *Altamira* = 5 and *Vai-Quem-Quer* = 5). DNA extractions were obtained using the commercial PureLink Genomic DNA Kit (Invitrogen Corp., Walthan, MA, USA) following the manufacturer’s instructions. Fragments of the 16S rRNA mitochondrial gene were amplified by polymerase chain reaction (PCR) using the universal primers 16sar and 16sbr (Palumbi, 1996). The PCR and DNA sequencing protocols followed Maia et al. (2017). We used Geneious 2021.2.2 (Kearse et al., 2012) to assemble contigs and edit consensus sequences by direct inspection of the original chromatographs. Sequences ranging in length from 453 to 608 base pairs (bp) are deposited in GenBank (ON997545–ON997564 and OQ297604–OQ297608).

### Phylogenetic analysis

We downloaded from GenBank homologous 16S sequences (*n* = 85) of nominal and candidate species of *Allobates* as well as outgroups. These included representative sequences of all lineages previously assigned to *Allobates gasconi* (Grant et al., 2006; Lima et al., 2014; Melo-Sampaio et al., 2018; Vacher et al., 2020). To improve phylogenetic inference, sequences of the following four mitochondrial genes were also retrieved: 12S ribosomal RNA (12S, *n* = 58), cytochrome c oxidase I (COI, *n* = 50), cytochrome b (CYTB, *n* = 65) and NADH dehydrogenase 1 (ND1, *n* = 42). We include as outgroups sequences from species of the superfamily Dendrobatoidea, *Ameerega* Bauer (1986), *Anomaloglossus* Grant et al. (2006), *Aromobates* Myers, Paolillo-O and Daly (1991), *Colostethus* Cope (1866), *Dendrobates* Wagler (1830), *Leucostethus* Grant et al. (2017), *Mannophryne* La Marca (1992), *Phyllobates* Bibron (1840), *Rheobates* Grant et al. (2006) and *Silverstoneia* Grant et al. (2006). Sequences were aligned through the MAFFT algorithm server (https://mafft.cbrc.jp/) using the E-INS-i strategy with default parameters (Katoh & Standley, 2013) for noncoding genes (12S and 16S) and the G-INS-I strategy for protein-coding genes (COI, CYTB and ND1). The final concatenated alignment was composed 6,251 bp and 110 terminals. Supplemental Table 1 lists vouchers and GenBank accession numbers.

The concatenated alignment was divided into one partition for 12S+16S and one for each codon of COI, CYTB and ND1. ModelFinder (Kalyaanamoorthy et al., 2017) was used to infer the following best partition schemes and respective best-fit evolutionary models: 12S+16S = GTR+F+R6; Cytb/1+ND1/1= SYM+I+G4; Cytb/2+ND1/2+COI/2= TN+F+I+G4; Cyt/3+ND1/3+COI/3= TN+F+R5; and COI/1 = SYM+I+G4. Phylogenetic relationships were inferred though Maximum Likelihood using IQ-TREE (Trifinopoulos et al., 2016). Clade support was calculated through 10,000 ultrafast bootstrap approximation replicates (Hoang et al., 2018) with 10,000 maximum iterations, a minimum correlation coefficient of 0.99 and 10,000 replicates of the Shimodaira-Hasegawa approximate likelihood ratio. Uncorrected and Kimura two-parameter distances (Kimura, 1980) between recently collected specimens of *Allobates gasconi* and other lineages previously assigned to it were calculated with MEGA 11 (Tamura et al., 2021) using pairwise deletion.

In addition to phylogenetic relationships and genetic distances (*p* and K2P), we use the Bayesian implementation of the Tree Poisson Process model (bPTP; Zhang et al., 2013) to infer whether topotypic individuals collected in *Jainú West*, *Jainú East*, *Altamira*, *Vai-Quem-Quer* and those previously reported elsewhere in the literature as *A. gasconi* represent the same Operational Taxonomic Unit (OTU). A 16S phylogenetic tree inferred through ML in IQ-TREE with the same terminals (110) and parameters used in the multilocus tree was used as input in bPTP. The bPTP was run using default parameters in the webserver https://species.h-its.org/ptp/, except for amount of MCMC generations (400,000 instead 100,000).

### Adult morphology

Sex was determined using the presence (males) or absence (females) of vocal sac, vocal slits and swollen fingers. Sexual maturity of specimens from the type series was evaluated through dissection (presence of mature testes or oviducts); all recently collected specimens were considered adults because they were actively courting or mating when collected.

Adult morphology is described from recently collected specimens (*n* = 31) from populations molecularly assigned to *A. gasconi* sensu stricto and from well-preserved adults (*n* = 9) of the type series. With the aid of a micrometer coupled to a stereoscopic microscope, we took the following measurements: snout-to-vent length (SVL); head length (HL); head width (HW); snout length (SL); eye–nostril distance (EN); internarial distance (IN); eye length (EL); interorbital distance (IO); tympanum diameter (TYM); forearm length (FAL); arm length (AL); hand length from proximal edge of palmar tubercle to tip of fingers I–IV (HANDI–IV); width of disc on finger III (WFD); width of distal phalanx of finger III (WPF); diameter of palmar tubercle (DPT); width of thenar tubercle (WTT); tibia length (TL); thigh length (THL); foot length (FL); width of disc on toe IV (WTD). HANDI–IV was used to compare length of fingers (Kaplan, 1997) because it is difficult to determine the intersection between palm and fingers in some instances (Grant et al., 2006). Morphological terminology follows Grant et al. (2006), except for finger counting. Description of adult morphology follows Lima et al. (2014). Description of adult coloration in life is based on field notes and on digital photographs by A. P. Lima, A. S. Ferreira and W. E. Magnusson. Raw morphometric measurements are provided in Table S2.

Sexual dimorphism is common in anuran species (Shine, 1979), being expressed sometimes in morphology, chromatism, or both. In nurse frogs, it is commonly reported for color pattern (e.g. Morales, 2002; Moraes et al., 2019; Melo-Sampaio et al., 2020; Lima et al., 2020), and rarely in morphometric traits (Ferrão et al., 2022). We describe sexual dimorphism in color pattern qualitatively but we tested sexual dimorphism in morphometric measurements using the Wilcoxon rank sum test with continuity correction. Wilcoxon tests were used only on recently collected individuals to avoid bias related to dehydration of preserved specimens. Analyses were run in the R environment (R Core Team, 2022) using the function *wilcox.test* of the package *stats*.

### Larval morphology

Larval developmental stages were determined according to Gosner (1960). Terminology and diagnostic characters follow Altig & McDiarmid (1999) and Schulze et al. (2015). Twenty-eight morphometric measurements were taken from 25 tadpoles with a micrometer coupled to a stereoscopic microscope and follow Moraes & Lima (2021). Measurements included body width at level of spiracle (BW); head width at level of eyes (HWLE); interorbital distance (IOD); internarial distance (IND); body length (BL); tail length (TAL); total length (TL); spiracle-snout distance (SS); body height at level of spiracle (BH); tail muscle maximum width (TMW); tail muscle maximum height (TMH); maximum tail height (MTH); eye diameter (ED); eye–nostril distance (END); nostril-snout distance (NSD); spiracle-tube length (STL); vent tube length (VTL); oral disc width (ODW); posterior labium width (PL); anterior labium width (AL); length of median gap in tooth row A2 (GAP); length of tooth rows A1 (LA1) and A2 (LA2); length of tooth rows P1 (LP1), P2 (LP2) and P3 (LP3); upper jaw sheath width (UJW); upper jaw-sheath length (UJL). Color in life was described from photographs and field notes. Raw morphometric measurements are provided in a Table S3.

### Bioacoustics

The advertisement call of *Allobates gasconi* is emitted in three distinct structural arrangements. Whenever possible, five calls of each arrangement were analyzed for each recorded male, totaling 142 analyzed calls. Using Raven 1.6 (Bioacoustics Research Program, 2015), we measured the following acoustic parameters: call duration, inter-call interval, call rate (number of calls per minute), number of notes per call, note duration, inter-note interval and dominant, lower and upper frequencies. Raven 1.6 was set as follows: Blackman window, DFT size 1,024 samples, 3 dB filter bandwidth 82 Hz, hop size 3.99 ms. Call duration was quantified considering the three structural arrangements that *A. gasconi* presents; note duration was quantified for the first and last notes of each call; inter-note interval was quantified between the first and second notes, and also before the last note in calls with 3–4 notes. Frequencies were measured in the first and last notes of each call; dominant frequency was measured trough the function *Peak Frequency*; lower and upper frequencies were measured 20 dB below the peak frequency to avoid background-noise interference.

Call terminology follows the call-centered approach as suggested by Köhler et al. (2017) who illustrated them in Figures 7B and 9. Graphic representation of advertisement calls was produced in R using the package *seewave* (Sueur et al., 2008). *Seewave* was set up as follows: window = Hanning, FFT size = 256 samples, and FFT overlap = 85%. Bioacoustics raw data are provided in Table S4 and S5.

### Species comparisons

We compare *Allobates gasconi* sensu stricto with nurse frogs distributed in southwestern Amazonia and closely related species belonging to the *A. trilineatus* clade (sensu Réjaud et al., 2020): *A. algorei* Barrio-Amorós and Santos, 2009, *A. amissibilis* Kok, Hölting, and Ernst, 2013, *A. bacurau* Simões, 2016, *A. caldwellae* Lima, Ferrão, and Silva, 2020, *A. chalcopis* (Kaiser, Coloma, and Gray, 1994), *A. conspicuus* (Morales, 2002), *A. femoralis* (Boulenger, 1884), *A. flaviventris* Melo-Sampaio, Souza, and Peloso, 2013, *A. fuscellus* (Morales, 2002), *A. granti* (Kok, MacCulloch, Gaucher, Poelman, Bourne, Lathrop, and Lenglet, 2006), *A. hodli* Simões, Lima, and Farias, 2010, *A. humilis* (Rivero, 1980), *A. insperatus* (Morales, 2002), *A. juami* Simões, Gagliardi-Urrutia, Rojas-Runjaic, and Castroviejo-Fisher, 2018, *A. kamilae* Ferrão, Hanken, and Lima, 2022; *A. marchesianus* (Melin, 1941), *A. melanolaemus* (Grant and Rodriguez, 2001), *A. nidicola* (Caldwell and Lima, 2003), *A. ornatus* (Morales, 2002), *A. pacaas* Melo-Sampaio, Prates, Peloso, Recoder, Vechio, Marques-Souza, and Rodrigues, 2020, *A. paleovarzensis* Lima, Caldwell, Biavati, and Montanarin, 2010, *A. pittieri* (La Marca, Manzanilla, and Mijares-Urrutia, 2004), *A. sieggreenae* Gagliardi-Urrutia, Castroviejo-Fisher, Rojas-Runjaic, Jaramillo-Martinez, Solís, and Simões, 2021, *A. subfolionidificans* (Lima, Sanchez, and Souza, 2007), *A. sumtuosus* (Morales, 2002), *A. tinae* Melo-Sampaio, Oliveira, and Prates, 2018, *A. trilineatus* (Boulenger, 1884), *A. vanzolinius* (Morales, 2002) and *A. velocicantus* Souza, Ferrão, Hanken, and Lima, 2020.

## Results

### Phylogenetic relationships and genetic distances

*Allobates* was recovered as monophyletic in the multilocus phylogeny (Fig. 2A). The monophyly of all currently recognized species clades is strongly supported. The recovered topology of major clades is identical to that of Réjaud et al. (2020) and similar to other recent phylogenies (e.g. Melo-Sampaio et al., 2018). The *A. caeruleodactylus* and *A. trilineatus* clades are sister taxa. Specimens of *A. gasconi* from the type locality (*Jainú West*) and two paratype localities (*Jainú East* and *Altamira*) in the middle Juruá River nest together in a well-supported lineage within the *A. trilineatus* clade (Fig. 2B). A Peruvian specimen from Ucayali recently reported as *Allobates* sp. “Ucuyali” by Réjaud et al. (2020) nests as sister to the *A. gasconi Jainú West+East+Altamira* lineage, with a very low uncorrected *p*-distance between them (0.7%; Table 1). The bPTP algorithm delimited them as a single Operational Taxonomic Unit (OUT) (Fig. 2B). We accordingly regard Peruvian and Brazilian specimens as conspecific and hereafter refer to them as *A. gasconi* sensu stricto (SS).

**Fig. 2.**
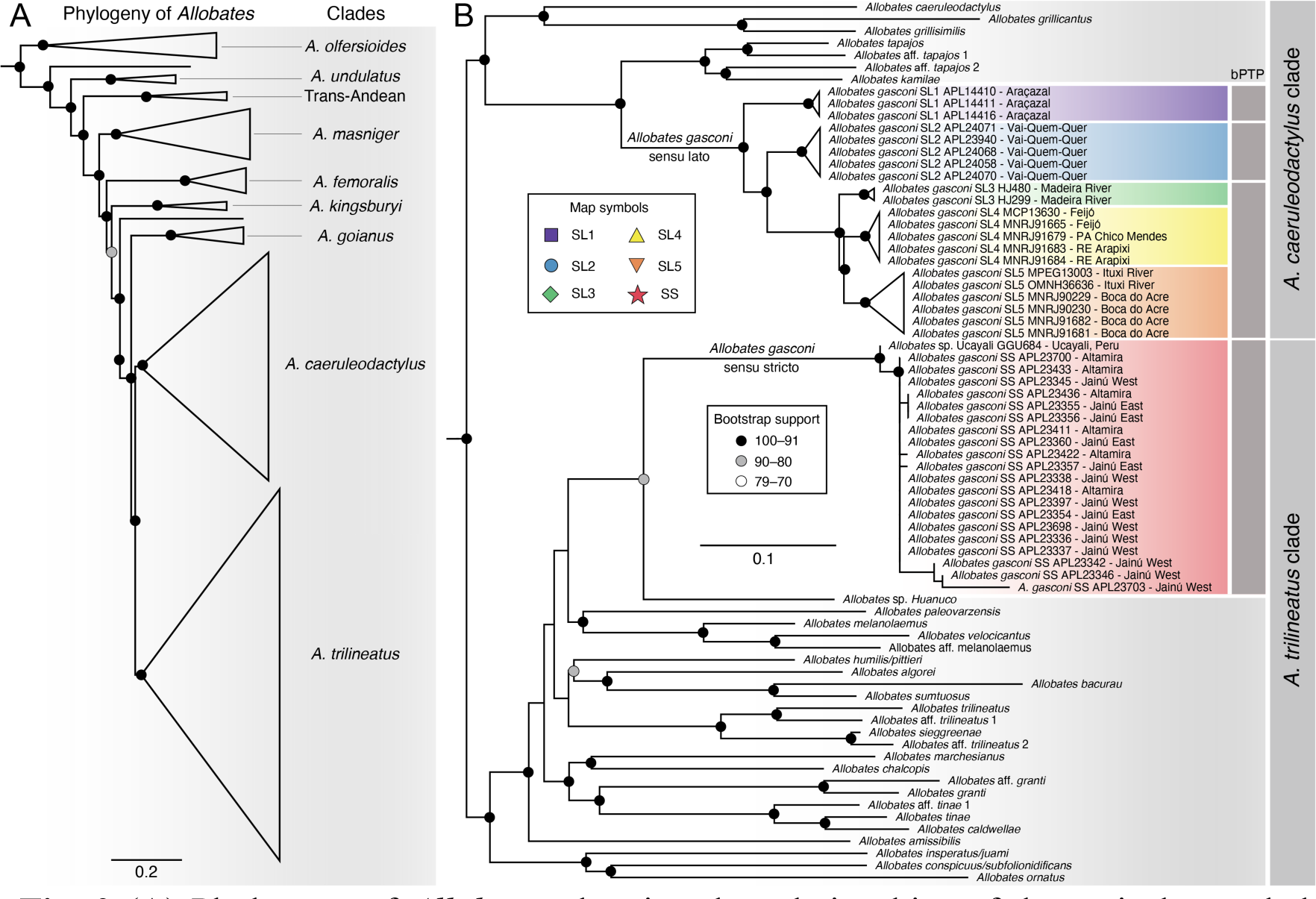
(A) Phylogeny of *Allobates* showing the relationships of the main large clades inferred through Maximum Likelihood based on five mitochondrial genes (12S, 16S, COI, Cytb, ND1) and 6251 bp. (B) A detailed view of the *A. caeruleodactylus* and *A. trilineatus* clades. Dark grey vertical bars denote species delimitation from the Bayesian Poisson Tree Processes model (bPTP). Abbreviations: SL, sensu lato; SS, sensu stricto. Map symbols are the same as those used in Figure 1. Scale bars represent substitutions per site. The unreduced phylogeny is presented in Figure S1. Illustration: M. Ferrão.

**Table 1.** Pairwise interspecific and intraspecific genetic distances between *Allobates gasconi* sensu stricto and closely related taxa. Uncorrected p-distances (below the diagonal) and kimura two-parameter distances (above the diagonal) are based on a fragment of the 16s rRNA mitochondrial gene and expressed as percentages. Numbers in bold represent intraspecific *p*-distance values. Abbreviation: SL, sensu lato; SS, sensu stricto; WE, west + east.

| Taxa | 1 | 2 | 3 | 4 | 5 | 6 | 7 | 8 |
| --- | --- | --- | --- | --- | --- | --- | --- | --- |
| 1 *Allobates* sp. Ucayali | – | 6.6 | 8.7 | 8.3 | 8.8 | 9.4 | 9.2 | 0.7 |
| 2 *Allobates* sp. Huanuco | 6.3 | – | 8.0 | 8.9 | 12.3 | 9.4 | 11.8 | 7.0 |
| 3 *Allobates gasconi* SL1 | 8.2 | 7.6 | **0.1** | 3.1 | 4.6 | 3.9 | 4.3 | 9.9 |
| 4 *Allobates gasconi* SL2 *Vai-Quem-Quer* | 7.8 | 8.4 | 3.0 | **0.2** | 3.7 | 3.1 | 3.8 | 9.4 |
| 5 *Allobates gasconi* SL3 | 8.3 | 11.2 | 4.7 | 3.8 | **0.4** | 1.8 | 2.2 | 9.8 |
| 6 *Allobates gasconi* SL4 | 8.7 | 8.8 | 4.1 | 3.2 | 1.9 | **0.2** | 1.9 | 10.5 |
| 7 *Allobates gasconi* SL5 | 8.6 | 10.8 | 4.5 | 3.9 | 2.3 | 1.9 | **0.6** | 10.3 |
| 8 *Allobates gasconi* SS *Jainú* WE+ *Altamira* | 0.7 | 6.7 | 9.2 | 8.7 | 9.2 | 9.7 | 9.6 | **0.4** |

All specimens reported as *Allobates gasconi* in previously published phylogenies (e.g. Grant et al., 2006; Lima et al., 2014; Melo-Sampaio et al., 2018; Réjaud et al., 2020; Vacher et al., 2020) group together in a large clade within the *A. caeruleodactylus* clade (Fig. 2B). Surprisingly, specimens of *A. gasconi* collected by us at the paratype locality in the lower Juruá River (*Vai-Quem-Quer*) also nest in the *A. caeruleodactylus* clade (Fig. 2B), a distant phylogenetic position from topotypic specimens of *A. gasconi SS* from the middle Juruá River. The bPTP algorithm applied to the 16S tree delimited the populations of *Vai-Quem-Quer* and *Jainú West+East+Altamira* as distinct OTUs. Moreover, the uncorrected *p*-distance between these populations is highly divergent (8.7%; Table 1). We accordingly regard them as heterospecific. All the *A. gasconi* lineages within the *A. caeruleodactylus* clade are hereafter referred to as *A. gasconi* sensu lato (SL1–5).

The clade that groups specimens of *A. gasconi* SL2 (*Vai-Quem-Quer*) is sister to a major clade formed by *A. gasconi* SL3 (Madeira River) + (SL4 [Tarauacá and Purus basins] + SL5 [Ituxí and Purus rivers]). The *A. gasconi* SL1 reported by Lima et al. (2014) from the middle Juruá River (Araçazal Village) is recovered as sister to SL2–5 (Fig. 2B). Uncorrected *p*-distances between *A. gasconi* SL2 (*Vai-Quem-Quer*) and its sister taxa, which range from 3.0% (SL2 *vs.* SL1) to 3.9% (SL2 *vs.* SL5), are much lower than its distances to *A. gasconi* SS (Table 1). The bPTP algorithm delimited *A. gasconi* SL1, *A. gasconi* SL2 and *A. gasconi* SL3–5 as three distinct OTUs (Fig. 2B).

### Taxonomic remarks

The type series of *Colosthetus gasconi* comprises adults and juveniles from the type locality (*Jainú West*) and five paratype localities (*Jainú East*, *Condor*, *Altamira*, *Nova Vida* and *Vai-Quem-Quer*) (Morales, 2002). The bPTP delimitation, phylogenetic relationships and genetic distances support specimens from the type locality, *Jainú East* and *Altamira* as conspecifics. However, this is not true for specimens from *Vai-Quem-Quer*.

Only three species of *Allobates* (*A. femoralis*, *A. vanzolinius* and *A. gasconi* SL2) were found at *Vai-Quem-Quer* and adjacent regions after exhaustive sampling in different environments (flooded forests, unflooded forests and dense- and open-canopy forests, both little and moderately anthropized). *Allobates gasconi* SL2 was the most widespread species, both locally and regionally, occurring in all surveyed environments. In contrast, *A. gasconi* SS from *Jainú West*, *Jainú East* and *Altamira* was found exclusively in environments with a flooding regime (e.g. *várzea* forests).

In addition to habitat preference, *A. gasconi* SL2 is readily distinguished from typical *A. gasconi* SS based on external coloration. For example, the throat and vocal sac in breeding males of *A. gasconi* SS are grey or dark grey, while *A. gasconi* SL2 has a bright yellow throat and vocal sac with a few melanophores in the central and lower regions (Fig. S1). Similarly, female *A. gasconi* SS have light to bright yellow throats, while the throat is white in female *A. gasconi* SL2 (Fig. S1). Based on molecular, morphological and ecological evidence, we exclude the specimen INPA-H 4889 from the type series of *Allobates gasconi* and the following redescription, as this specimen clearly belongs to another species.

### Taxonomic account

***Allobates gasconi* (Morales, 2002)**

*Colostethus gasconi* Morales (2002)

*Allobates* sp. “Ucuyali” Réjaud et al. (2020)

#### Holotype

INPA-H 3082 (Fig. 3), sub-adult male from *Jainú West*, west bank of the middle Juruá River, municipality of Itamarati, state of Amazonas, Brazil, collected by Claude Gascon on 17 October 1991.

**Fig. 3.**
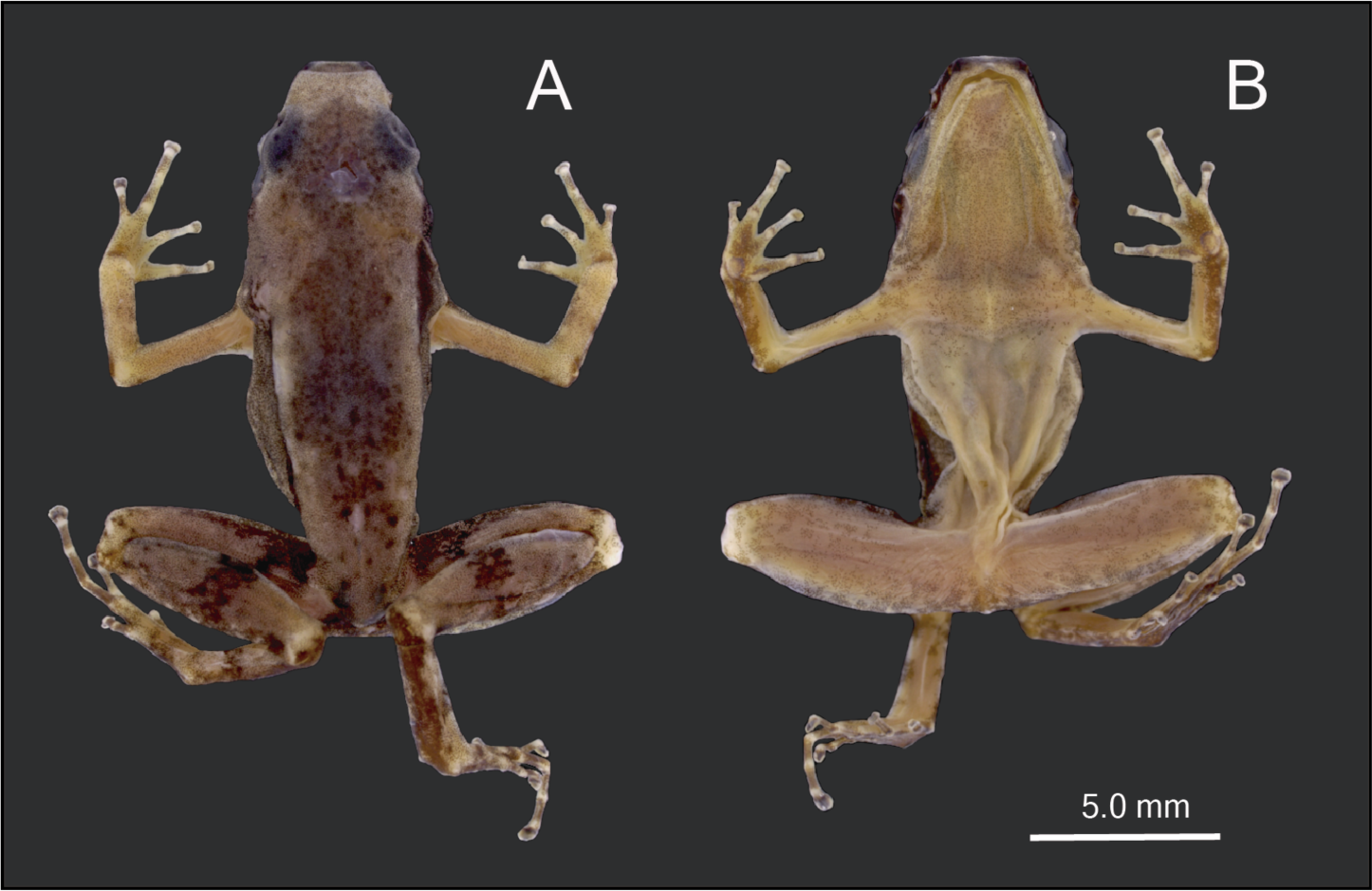
(A) Dorsal and (B) ventral views of the sub-adult male holotype (INPA-H 3082) of *Allobates gasconi* from *Jainú West*, Juruá River, Amazonas, Brazil. Photographs by A. S. Ferreira.

#### Paratypes

Thirty–three individuals, all collected by Claude Gascon in the Juruá River, Brazil. West bank: *Jainú West*: six juveniles (INPA-H 3150–51, 3172, 3406, 3412 and 3415), 19–29 October 1991; two adult males (INPA-H 3512–13), 03 November 1991. East bank: *Condor*: one juvenile (INPA-H 2649), 19 September 1991. *Jainú East*: fifteen juveniles (INPA-H 3066, 3068, 3075, 3078, 3084–85, 3087, 3090, 3094, 3220, 3401, 3483–84, 3494 and 3496), 17–29 October 1991 and 2 November 1991; three adult females (INPA-H 3055, 3079 and 3491), 16–17 October 1991 and 2 November 1991; three adult males (INPA-H 3083, 3093 and 3249), 17–23 October 1991. *Altamira*: one adult male (INPA-H 3541), 9 November 1991. *Nova Vida*: one juvenile (INPA-H 4726), 21 March 1992.

#### Referred specimens

Thirty-one adults, all from the middle Juruá River, municipality of Itamarati, state of Amazonas, Brazil, collected by A.S. Ferreira, A.P. Lima and M. Ferrão. West bank: *Jainú West*: five females (field numbers APL 23345–48 and 23350) and twelve males (field numbers APL 23336–39, 23341–44, 23349, 23351–52 and 23397), 22–24 January 2021. East bank: *Jainú East*: one female (field number APL 23358) and six males (field numbers APL 23354–57, 23359–60), 23 January 2021.

#### Altamira

one female (field number APL 23410) and six males (field numbers APL 23411, 23418–19, 23422, 23433, 23436), 25 and 26 January 2021.

#### Reappraisal diagnosis

*Allobates gasconi* is a small nurse frog diagnosed by the following combination of characters: SVL 14.8–17.1 mm in males (n = 30) and 16.0– 18.0 mm in females (n = 10); finger III strongly swollen in adult males; expanded disk on all fingers; one subarticular tubercle on finger IV; basal toe webbing between toes II–III and III–IV; lateral fringes present on pre- and postaxial edges of all fingers and toes; dark brown hourglass-shaped mark on dorsum; males with a gray to dark gray throat; advertisement call comprises 2–4 notes emitted repeatedly, with a dominant frequency of 5,081–6,761 Hz; larvae with labial keratodont row formula 2(2)/2(1).

*Allobates gasconi* is easily distinguished from *A. amissibilis*, *A. bacurau*, *A. caldwellae*, *A. chalcopis*, *A. conspicuus*, *A. femoralis*, *A. fuscellus*, *A. granti*, *A. hodli*, *A. insperatus*, *A. juami*, *A. marchesianus*, *A. nidicola*, *A. paleovarzensis*, *A*. *sieggreenae*, *A. subfolionidificans*, *A. sumtuosus*, *A. tinae*, *A. trilineatus*, *A. vanzolinius* and *A. velocicantus* by having a distinct dark brown hourglass-shaped mark on the dorsum (absent in the other species). Additionally, *A. gasconi* differs from *A*. *caldwellae*, *A. conspicuous*, *A. chalcopis*, *A. femoralis*, *A. granti*, *A. hodli*, *A. insperatus*, *A. juami*, *A. marchesianus*, *A. nidicola* and *A. tinae* by having males with finger III strongly swollen (weakly or not swollen in the other species); from *A. paleovarzensis* and *A. vanzolinius* by its maximum SVL 17.1 mm in adult males (minimum SVL 18.3 mm in *A. paleovarzensis*, 22.9 mm in *A. vanzolinius*); from *A*. *sieggreenae*, *A*. *subfolionidificans* and *A. velocicantus* by having males with a gray to dark gray throat in life (white to translucent in *A*. *sieggreenae*, opaque white in *A*. *subfolionidificans*, white centrally and yellow marginally in *A. velocicantus*); from *A. bacurau* and *A. sumtuosus* by having dark brown transverse bars on the dorsal surface of the thighs (absent in the other species); from *A. trilineatus* by the absence of ventrolateral stripe (present in *A. trilineatus*); from *A. amissibilis* by having a conspicuous pale dorsolateral stripe present (absent in *A. amissibilis*); and from *A. fuscellus* by having lateral fringes on toes (absent in *A. fuscellus*).

The dark brown hourglass-shaped mark on the dorsum of *A. gasconi* is similar to that of *A. algorei*, *A. flaviventris*, *A. humilis*, *A. kamilae*, *A*. *melanolaemus*, *A. ornatus*, *A. pacaas* and *A. pittieri*. However, *A. gasconi* differs from *A. algorei*, *A. flaviventris*, *A. humilis*, *A*. *melanolaemus* and *A. pacaas* by having finger III greatly swollen in adult males (weakly swollen in all the other species); from *A. flaviventris*, *A. humilis* and *A*. *melanolaemus* by the maximum SVL 18.0 mm in adults (minimum SVL 19.3 mm in *A. flaviventris*, 21.8 mm in *A. humilis* and 21.1 mm in *A*. *melanolaemus*); from *A. kamilae* and *A. ornatus* by having males with a gray to dark gray throat in life (bright yellow in *A. kamilae*, pale lemon yellow with scattered melanophores in *A. ornatus*); from *A. algorei* by having basal toe webbing (absent); from *A. ornatus* by having lateral fringes on toes and oblique lateral stripe (absent); from *A. pittieri* by having expanded disk on finger III (not expanded) and absence of cloacal sheath (present); and from *A. pacaas* by having one subarticular tubercle on finger IV (two tubercles).

The advertisement call of *Allobates gasconi* is easily distinguished from those of *A. algorei*, *A. amissibilis*, *A. bacurau*, *A. caldwellae*, *A. femoralis*, *A. humilis*, *A. hodli*, *A. juami*, *A. kamilae*, *A. marchesianus*, *A. melanolaemus*, *A. nidicola*, *A. paleovarzensis*, *A. sieggreenae*, *A. subfolionidificans*, *A. sumtuosus* and *A. tinae* by having 2–4 notes (calls composed of one note in the other species). Moreover, calls of *A. gasconi* are emitted with a dominant frequency of 5,081–6,761 Hz, which differs from *A. femoralis*, *A. humilis*, *A. hodli*, *A. nidicola* and *A. paleovarzensis* (3,100–3,375 Hz in *A. femoralis*; 4,200 Hz in *A. humilis*; 2,900–3,600 Hz in *A. hodli*; 3,759–4,689 Hz in *A. nidicola*; and 4,050–4,930 Hz in *A. paleovarzensis*); from *A. bacurau*, *A. caldwellae*, *A. juami*, *A. marchesianus*, *A. melanolaemus*, *A. sieggreenae*, *A. sumtuosus* and *A. tinae* by having calls emitted continuously (calls arranged in call series in the other species). The only species with multi-note calls among those compared here are *A. flaviventris*, *A. granti*, *A. trilineatus* and *A. velocicantus*. However, *A. gasconi* differs from *A. granti*, *A. trilineatus* and *A. velocicantus* by having calls with 2–4 notes, that are emitted continuously (call consisting of 2 notes, arranged in call series with 6–11 calls in *A. trilineatus*; call series with 1–17 calls and average 6.2 ± 2.6 calls per series in *A. granti*; and call composed of 66–138 pulsed notes with a note duration of 5–13 ms in *A. velocicantus*), and from *A. flaviventris* by having a dominant frequency of 5,081– 6,761 Hz (3,618–4,651 Hz in *A. flaviventris*). The advertisement calls of *A. chalcopis*, *A. conspicuus*, *A. fuscellus*, *A. insperatus*, *A. ornatus*, *A. pacaas*, *A. pittieri* and *A. vanzolinius* remain unknown.

Tadpoles of *Allobates gasconi* have only two tooth rows on the posterior labium, which differs from *A. caldwellae*, *A. femoralis*, *A. granti*, *A. hodli*, *A. kamilae*, *A. paleovarzensis*, *A. pittieri*, *A. subfolionidificans*, *A. sumtuosus* and *A. velocicantus* (three rows in the other species) and from *A. nidicola* (teeth absent). Like *A. gasconi*, tadpoles of *A. marchesianus* have two tooth rows on the posterior labium, but *A. gasconi* also has two tooth rows on the anterior labium (only one row in *A. marchesianus*). Tadpoles of *A. algorei*, *A. amissibilis*, *A. bacurau*, *A. chalcopis*, *A. conspicuus*, *A. flaviventris*, *A. fuscellus*, *A. humilis*, *A. insperatus*, *A. juami*, *A. melanolaemus*, *A. ornatus*, *A. pacaas*, *A. sieggreenae*, *A. tinae*, *A. trilineatus* and *A. vanzolinius* have not been described.

#### Adult morphology

The following description of external morphology is based on adult males of the type series (n = 6) and recently collected males (n = 24) from *Jainú West*, *Jainú East* and *Altamira*. Coloration of males and females, is described in the next section (Intraspecific variation).

A small-bodied species, SVL 14.8–17.1 mm in males and 16.0–18.0 mm in females. Head slightly wider than long (HW/HL = 106 ± 8%), head width 30 ± 2% of SVL and head length 28 ± 2% of SVL. Snout blunt, rounded to nearly truncate in dorsal view and acutely rounded in lateral view; snout length 55 ± 6% of HL; canthus rostralis predominantly concave in dorsal view; internarial distance 49 ± 3% of HW; nostril opening directed laterally, and only visible laterally and ventrally; eye length 90 ± 5% of EN; tympanic membrane inconspicuous, rounded and small, TYM 38 ± 4% of EL; tongue attached anteriorly, longer than wide, slightly rounded posteriorly and with scattered small papillae; median lingual process absent; vocal sac distinct, single and subgular; teeth present on the maxillary arch.

Forearm slightly thicker and smaller than the upper arm (FAL/UAL = 92 ± 8%); forearm and upper arm length represent 22 ± 2% and 24 ± 2% of SVL, respectively; black arm glands absent. Hand small, HANDIII 24 ± 2% of SVL; paired dorsal digital scutes present; fingers weakly expanded, but preaxial swelling well developed on finger III (Fig. 4); width of the third phalange of finger III 82 ± 13% of WFD; basal finger webbing absent; when finger IV is appressed, its tip reaches the base of distal subarticular tubercle of finger III in most individuals; relative length of fingers III > I > II > IV; carpal pad absent; excrescences on thumbs males absent; palmar tubercle rounded, diameter 14 ± 2% of HANDIII (Fig. 4); thenar tubercle width 56–133% of the DPT; two subarticular tubercles on finger III, a single proximal subarticular tubercle on fingers I, II and IV (Fig. 4); distal subarticular tubercle of finger IV absent; supernumerary tubercle present on the outer edge of the palm; disc expanded on all fingers; disc on finger III wider than the others; poorly defined basal and lateral keels on pre- and postaxial edges of all fingers.

**Fig. 4.**
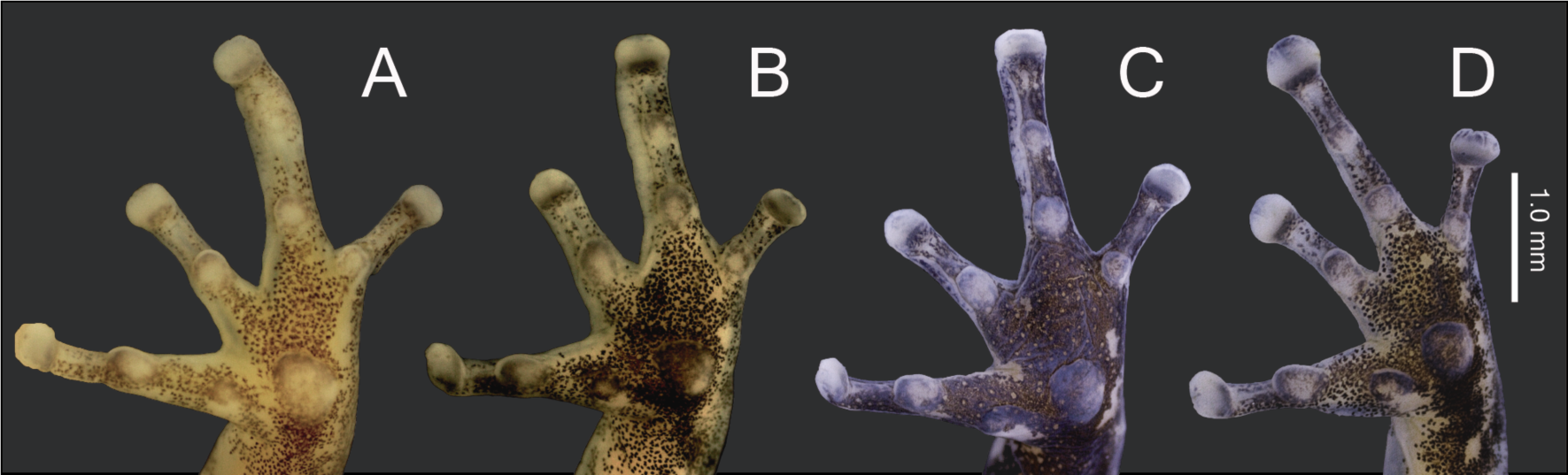
Ventral views of left hands of preserved male and female *Allobates gasconi*. (A) Adult male paratype (INPA-H 3249), (B, C) topotypic adult males (APL 23344 and APL 23356, respectively) and (D) topotypic adult female (APL 23345). Photographs by A. S. Ferreira.

Hind limbs robust; tibia slightly longer than thigh (TL/THL = 103 ± 6%); tibia and thigh length 47 ± 4% and 46 ± 3% of SVL, respectively; foot length 96 ± 5% of TL and 99 ± 7% of THL. Relative length of toes IV > III > V > II > I (Fig. 5); black band above elbow absent; basal webbing present between toes II–III and III–IV; tip of toe I reaches the subarticular tubercle on toe II; tip of toe III reaches the medial subarticular tubercle on toe IV; tip of toe V reaches the mid-level of the proximal phalange of toe IV (Fig. 5); discs of toes I and V moderately expanded, moderately to greatly expanded on toes II, III and IV (Fig. 5); width of disc on toe IV 69–150% of disc on finger III (Fig. 5). Inner metatarsal tubercle oval and outer metatarsal tubercle rounded; median metatarsal tubercle absent; tarsal keel short, tubercle-like and strongly curved at proximal end, extending from metatarsal tubercle; foot and tarsal fringe present; weak metatarsal fold present, small and swollen midway between the outer metatarsal tubercle and the first subarticular tubercle of toe V. A single subarticular tubercle on toes I and II, two subarticular tubercles on toes III and V, and three subarticular tubercles on toe IV; proximal tubercles small and poorly defined on toes III–V; disc on all toes wider than the penultimate phalanges; poorly developed lateral keels on preaxial and postaxial sides of all toes; basal keels between toes I–II and IV–V.

**Fig. 5.**
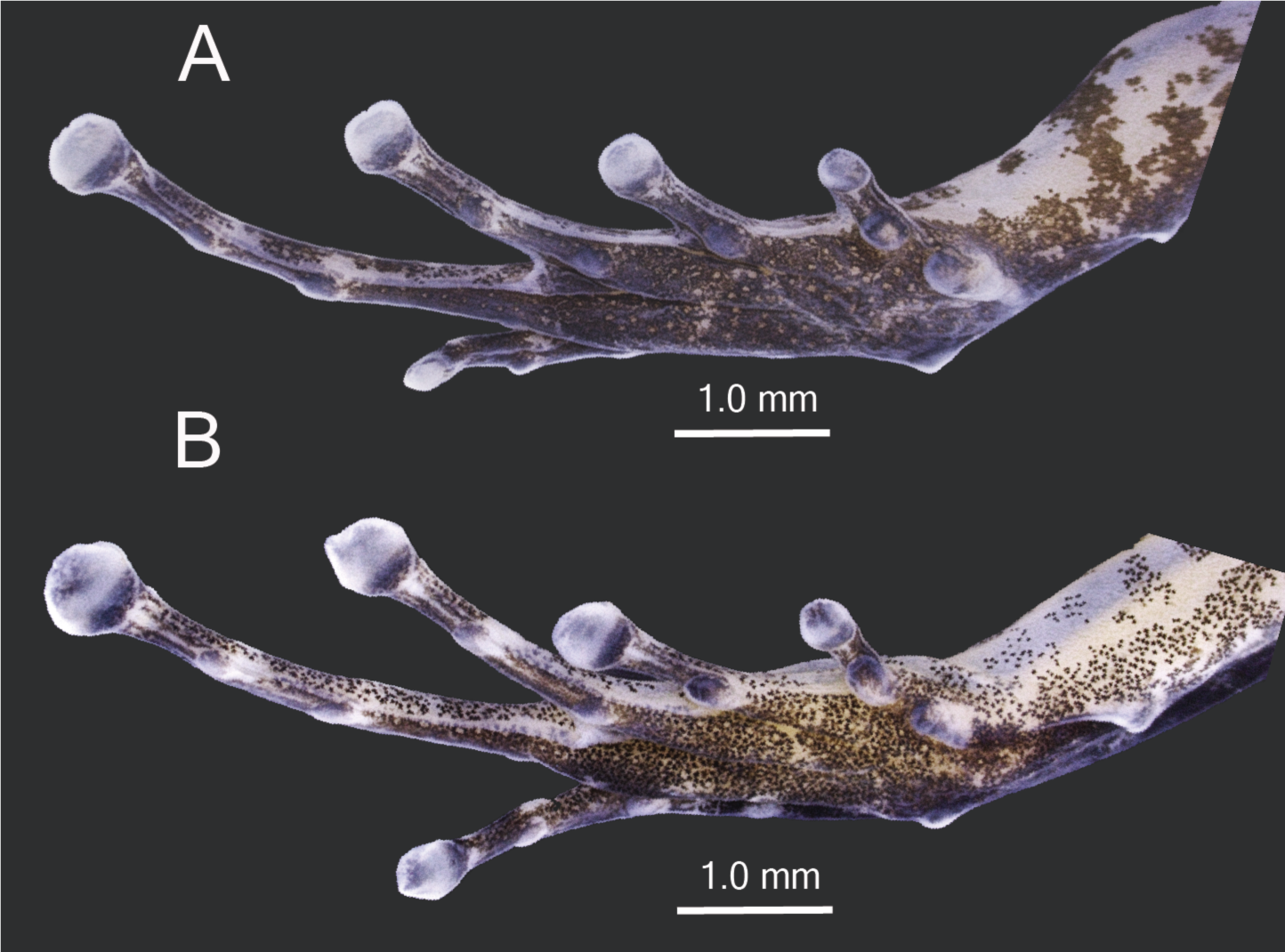
(A) Ventral surface of the right foot of a preserved topotypic adult male (APL 23356) and (B) right foot of topotypic adult female (APL 23345) of *Allobates gasconi*. Photographs by A. S. Ferreira.

Skin smooth on anterior portion of the dorsum, slightly tubercular from mid-body to the sacral region; dorsal surface of forelimbs smooth; dorsal surface of hind limbs with small and scattered irregular tubercles; flanks and ventral surface smooth.

#### Intraspecific variation

The adult morphology of the original type series and of additional recently collected specimens are similar. They differ only in the degree of swelling on finger III of males, which is less prominent in the type series, collected in 1991, than in the recent sample collected in 2021, likely due to reproductive state or a preservation artifact. Moreover, all specimens have similar dorsal color patterns, which differ mostly in color contrast. External coloration appears to have faded in the type series in comparison with the fresh material, probably due to the greater time in preservative and possibly due to different procedures of fixation and/or preservation.

Canthus rostralis is concave in 84% of specimens and straight in 16%; finger I longer than finger II in 95% of specimens, same size in 5%; finger II longer than finger IV in 70% of specimens, same size in 27.5% and shorter in 2.5%; finger III strongly swollen in adult males; when appressed, the tip of finger IV reaches the base of the distal subarticular tubercle of finger III in 74% of specimens but does not reach there in 26%; thenar tubercle elliptic in 52% of specimens and sub-circular in 48%; tarsal keel short, tubercle-like in all specimens; basal webbing present between toes II and III in 70% of specimens and between toes III and IV in all; basal keels present between toes of 78% of specimens; a small fold midway between the outer metatarsal tubercle and the first subarticular tubercle of toe V in 48% specimens.

*Allobates gasconi* is sexually dimorphic in several traits. The snout is longer in females (SL; *W* = 125.5, *p* = 0.045), as well as the width of disc on toe IV (WTD; *W* = 134.5, *p* = 0.009). On the other hand, females tend to be slightly larger than males (SVL; *W* = 123.5, *p* = 0.064). Morphometric measurements are summarized in Table 2.

**Table 2.** Morphometric measurements (in millimeters) of *Allobates gasconi* from the type series and from newly collected material. Values depict mean ± standard error and range (minimum–maximum). Trait acronyms are described in the text. Abbreviations: *n*, sample size.

| Traits | Type series |  |  | New material |  |
| --- | --- | --- | --- | --- | --- |
| | Males ($n = 6$) | Females ($n = 3$) | Juveniles ($n = 25$) | Males ($n = 24$) | Females ($n = 7$) |
| SVL | 15.4 ± 0.4 (14.8–15.9) | 16.6 ± 0.6 (16.0–17.1) | 14.5 ± 0.7 (13.1–15.7) | 16.2 ± 0.6 (14.8–17.1) | 16.8 ± 0.8 (16.0–18.0) |
| HL | 4.4 ± 0.3 (4.0–5.0) | 4.3 ± 0.2 (4.2–4.5) | 4.3 ± 0.3 (3.5–4.7) | 4.6 ± 0.4 (4.0–5.7) | 4.7 ± 0.3 (4.3–4.9) |
| HW | 4.6 ± 0.3 (4.2–5.0) | 4.4 ± 0.4 (4.0–4.7) | 4.3 ± 0.3 (3.5–5.0) | 4.9 ± 0.3 (4.3–6.0) | 4.9 ± 0.2 (4.5–5.1) |
| SL | 2.2 ± 0.2 (2.1–2.5) | 2.2 ± 0.0 (2.1–2.2) | 2.1 ± 0.2 (1.6–2.6) | 2.5 ± 0.2 (2.1–2.9) | 2.7 ± 0.1 (2.5–2.8) |
| EN | 1.5 ± 0.1 (1.4–1.6) | 1.6 ± 0.0 (1.6–1.6) | 1.5 ± 0.1 (1.3–1.8) | 1.9 ± 0.2 (1.4–2.1) | 1.9 ± 0.1 (1.8–2.0) |
| IN | 2.3 ± 0.1 (2.2–2.4) | 2.3 ± 0.2 (2.1–2.5) | 2.1 ± 0.1 (1.9–2.4) | 2.4 ± 0.1 (2.2–2.7) | 2.5 ± 0.1 (2.3–2.6) |
| EL | 2.1 ± 0.0 (2.0–2.1) | 2.1 ± 0.1 (2.0–2.2) | 1.9 ± 0.2 (1.6–2.2) | 2.1 ± 0.1 (1.9–2.4) | 2.3 ± 0.1 (2.1–2.5) |
| IO | 4.2 ± 0.4 (3.7–4.9) | 4.4 ± 0.3 (4.0–4.6) | 4.0 ± 0.3 (3.4–4.6) | 4.7 ± 0.3 (4.3–5.5) | 4.7 ± 0.2 (4.3–4.9) |
| TYM | 0.7 ± 0.1 (0.6–0.8) | 0.8 ± 0.0 (0.8–0.8) | 0.7 ± 0.1 (0.5–0.9) | 0.8 ± 0.1 (0.6–0.9) | 0.9 ± 0.1 (0.6–0.9) |
| FAL | 3.1 ± 0.5 (2.3–3.6) | 3.4 ± 0.2 (3.2–3.6) | 2.9 ± 0.3 (1.9–3.4) | 3.6 ± 0.3 (2.9–4.4) | 3.7 ± 0.3 (3.5–4.3) |
| UAL | 3.5 ± 0.2 (3.1–3.8) | 3.6 ± 0.1 (3.6–3.8) | 3.3 ± 0.4 (2.5–4.1) | 4.0 ± 0.2 (3.5–4.5) | 3.9 ± 0.2 (3.8–4.4) |
| HANDI | 2.6 ± 0.3 (2.3–3.0) | 2.5 ± 0.2 (2.4–2.8) | 2.4 ± 0.3 (1.9–3.0) | 2.9 ± 0.2 (2.4–3.3) | 3.0 ± 0.2 (2.8–3.2) |
| HANDII | 2.7 ± 0.1 (2.5–2.8) | 2.6 ± 0.1 (2.4–2.6) | 2.4 ± 0.3 (1.8–2.8) | 2.7 ± 0.2 (2.4–3.1) | 2.8 ± 0.1 (2.6–3.0) |
| HANDIII | 3.6 ± 0.2 (3.3–3.8) | 3.6 ± 0.1 (3.5–3.8) | 3.3 ± 0.4 (2.3–4.1) | 3.9 ± 0.3 (3.4–4.4) | 4.0 ± 0.2 (3.8–4.2) |
| HANDIV | 2.3 ± 0.2 (2.0–2.6) | 2.3 ± 0.2 (2.1–2.4) | 2.2 ± 0.2 (1.8–2.7) | 2.6 ± 0.3 (2.0–3.1) | 2.7 ± 0.1 (2.5–2.8) |
| WFD | 0.5 ± 0.1 (0.4–0.5) | 0.4 ± 0.0 (0.4–0.4) | 0.4 ± 0.0 (0.3–0.5) | 0.5 ± 0.1 (0.4–0.6) | 0.5 ± 0.0 (0.5–0.6) |
| TL | 7.0 ± 0.4 (6.5–7.5) | 7.0 ± 0.3 (7.0–7.5) | 6.6 ± 0.6 (4.8–7.8) | 7.7 ± 0.6 (6.6–9.2) | 7.7 ± 0.4 (7.2–8.1) |
| FL | 6.6 ± 0.4 (6.0–7.1) | 6.9 ± 0.4 (6.5–7.2) | 6.1 ± 0.5 (4.6–6.9) | 7.4 ± 0.5 (6.6–8.9) | 7.5 ± 0.4 (7.0–8.1) |
| THL | 7.1 ± 0.4 (6.5–7.7) | 7.1 ± 0.5 (6.6–7.5) | 6.8 ± 0.4 (6.0–7.5) | 7.4 ± 0.6 (6.5–8.8) | 7.4 ± 0.3 (7.0–7.8) |
| DPT | 0.5 ± 0.1 (0.3–0.6) | 0.5 ± 0.0 (0.4–0.5) | 0.4 ± 0.1 (0.3–0.5) | 0.6 ± 0.1 (0.4–0.8) | 0.5 ± 0.0 (0.5–0.6) |
| WTT | 0.3 ± 0.0 (0.3–0.4) | 0.4 ± 0.0 (0.3–0.4) | 0.4 ± 0.1 (0.3–0.6) | 0.5 ± 0.1 (0.3–0.7) | 0.5 ± 0.1 (0.3–0.7) |
| WTD | 0.4 ± 0.1 (0.3–0.6) | 0.5 ± 0.2 (0.3–0.6) | 0.5 ± 0.1 (0.3–0.6) | 0.6 ± 0.1 (0.5–0.7) | 0.7 ± 0.1 (0.6–0.8) |
| WPF | 0.4 ± 0.1 (0.3–0.6) | 0.3 ± 0.0 (0.3–0.3) | 0.3 ± 0.1 (0.2–0.5) | 0.4 ± 0.1 (0.3–0.5) | 0.3 ± 0.0 (0.3–0.4) |

There is no sexual dimorphism in dorsal coloration. In life, ground color varies from orange brown to light brown (Figs. 6, 7). A dark hourglass-shaped mark extends from the interorbital region to the posterior dorsum, ending before the cloacal opening (Fig. 8); it can be conspicuous (Fig. 8; APL 23341 and 23345) or inconspicuous (Fig. 8; APL 23356; INPA-H 3491). Scattered small brown, non-keratinized tubercles on posterior dorsum; medial line absent; pale cream dorsolateral stripe with melanophores has a regular border laterally but not medially; dark lateral stripe from snout tip to groin covers the upper half of the tympanum and is fainter but wider posteriorly; oblique lateral stripe present as a whitish but incomplete diffuse line or series of small diffuse spots that extend from the inguinal region to midbody; ventrolateral stripe absent, but irregular white spots present along the inferior edge of the lateral stripe; arm and forearm pale orange to brown, with dark brown longitudinal band along postaxial sides; anterior portion of arm light yellow or orange-white; dark disks with paired iridescent-white scutes. Dorsal surfaces of hind limbs pale orange or yellowish brown, with a transverse dark brown bar on thigh, shank and tarsus in 90%, 97.5% and 98% of specimens, respectively. Paracloacal mark whitish or cream, short, half-moon-shaped in 97% of specimens, always surrounded by a dark brown frame. Fingers brown to pale orange; toes light to dark brown with transverse white bars; paired dorsal digital scutes present on digits I, II and III, with coloration varying from white to black in 71% of specimens, and only white in 29%; digit IV is always black. Iris matte black in 45% of specimens and matte black and silver in 55%, with a thin, dark-brown reticulation (Fig. 7).

**Fig. 6.**
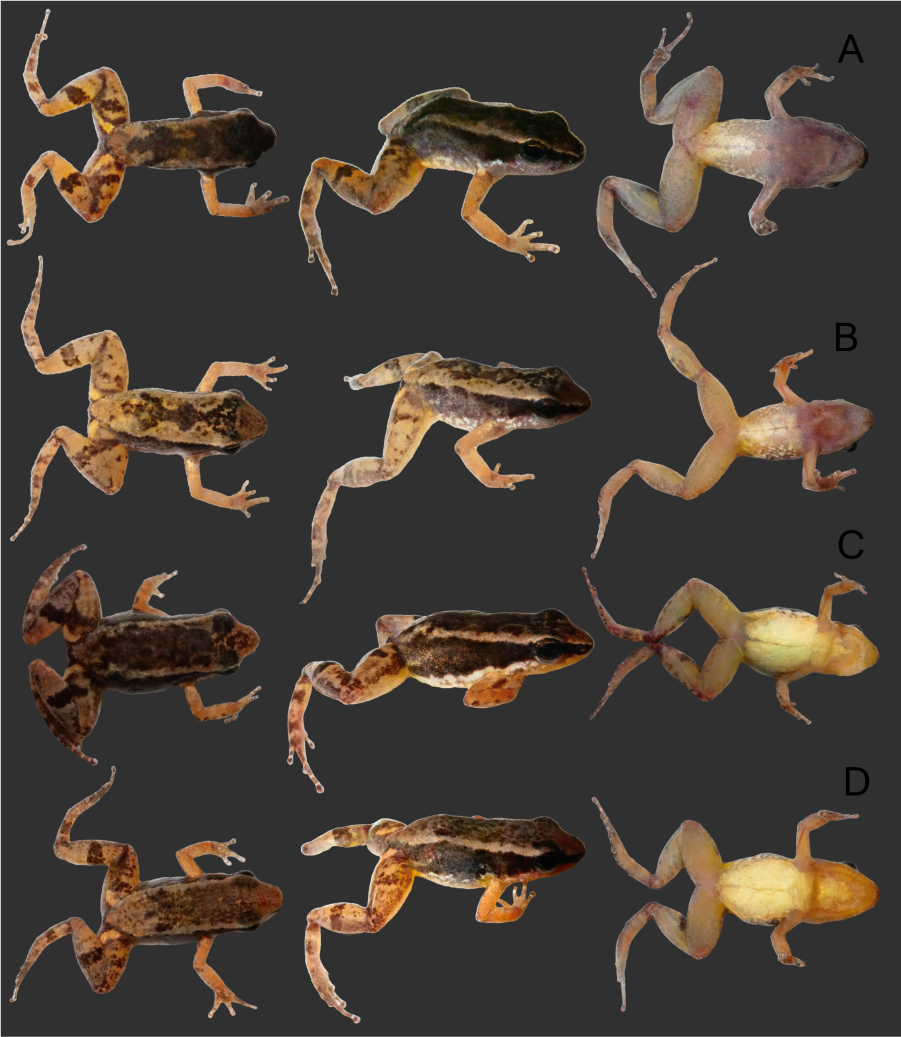
Color in life of adult males and females of *Allobates gasconi*. (A, B) Topotypic adult males, APL 23336 and APL 23339, respectively. (C, D) Topotypic adult females, APL 23346 and 23348, respectively. Photographs by A. P. Lima.

**Fig. 7.**
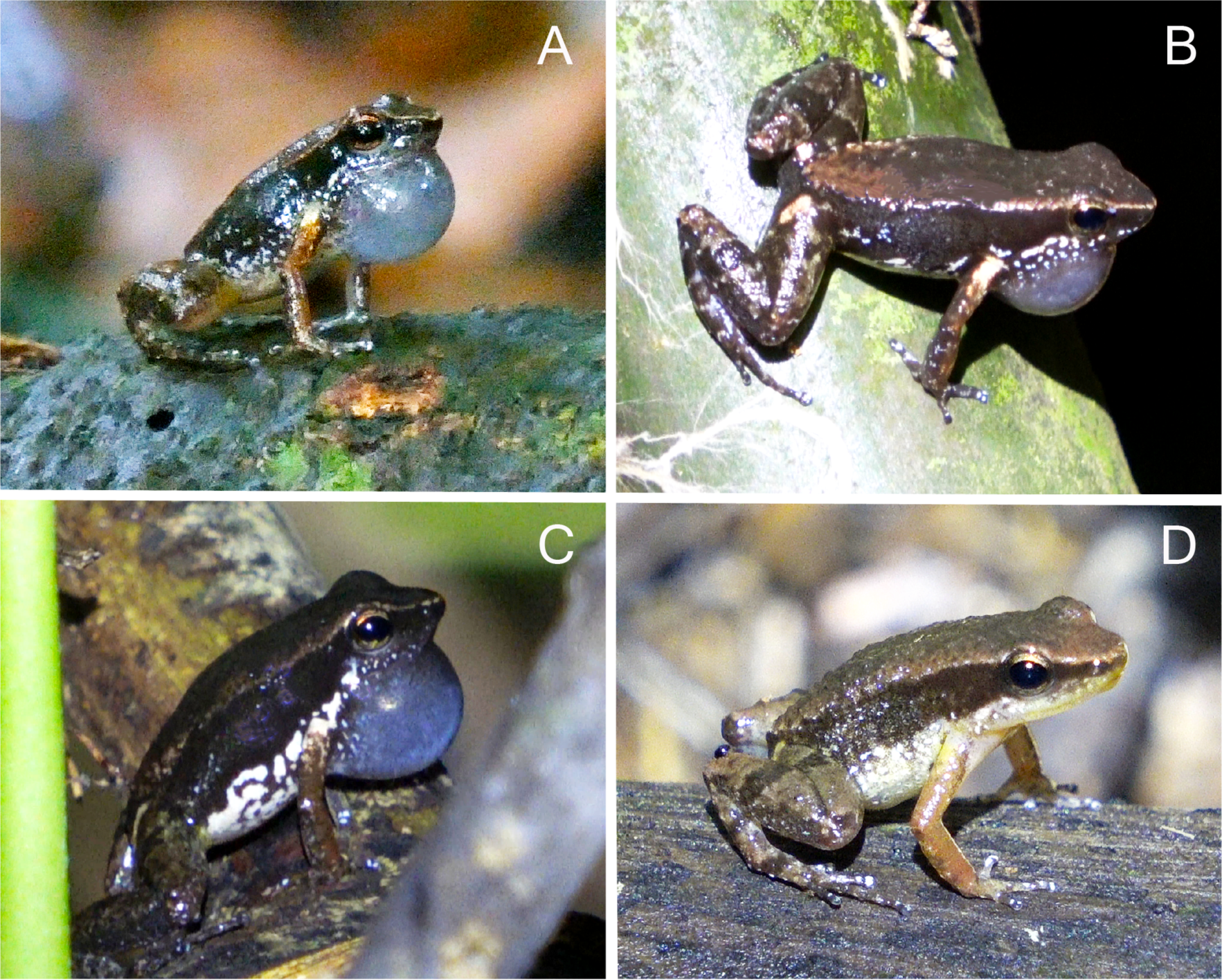
Coloration in life of topotypic adult males (A, C and D) and females (B) of *Allobates gasconi*. Note the densely pigmented throat and vocal sac of actively calling males. Photographs by A.P. Lima (B, C) and W. E. Magnusson (A, D).

**Fig. 8.**
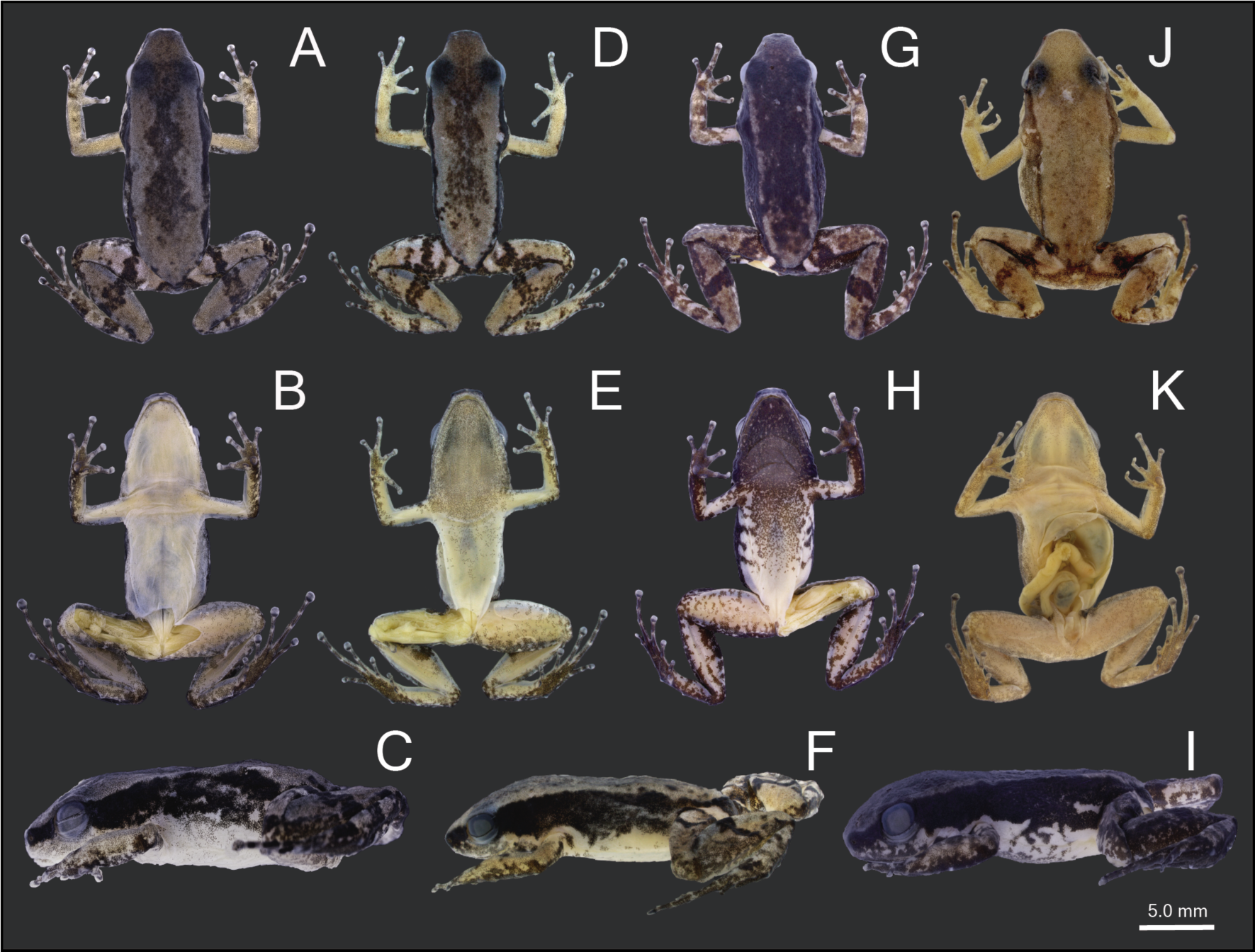
Dorsal, ventral and lateral color-pattern variation among preserved *Allobates gasconi*. (A–C) Adult female, APL 23345, from *Jainú West*. (D–F) adult male, APL 23341, from *Jainú West*. (G–I) adult male, APL 23356, from *Jainú East*. (J, K) adult female paratype, INPA-H 3491, from *Jainú East*. Photographs by A. S. Ferreira.

Ventral coloration is sexually dimorphic. In life, males have gray to dark gray throats and vocal sacs with a profusion of melanophores (Figs. 6, 7); females have bright- to whitish-yellow throats with few scattered dark melanophores on jaw edges. The chest is translucent to pinkish grey in males, and bright- to whitish-yellow without dark spots in females. In males, the belly is pinkish-grey anteriorly, whitish-grey centrally, light- or translucent-grey with white and brown small blotches laterally, and yellowish or translucent-grey posteriorly. Some males have melanophores over the entire belly, but most have densely concentrated melanophores until the midbody; one male recently collected has a darkly pigmented belly (Fig. 8; APL 23356). In females, the belly is uniformly yellowish-cream to whitish-yellow with few scattered melanophores on the outer margins. Both sexes have pale- to translucent-grey thighs, and shanks and tarsus greyish with a few scattered melanophores. Palmar and plantar surfaces light-brown, dark-grey or dark-brown, respectively.

In preservative (Fig. 8), dark dorsal marks are similar to when observed in life, but orange-brown shades have faded to pale-brown or pale-gray and dark-brown coloration has become blackish-gray. Paracloacal marks are paler than in life. White spots above lateral stripe have disappeared in preserved specimens, and ventral melanophores in males are less intense (Fig. 8; APL 23341 and 23356). Bright- and whitish-yellow parts, as well as light-grey and translucent-white ones, have become cream or light-cream in preserved specimens.

#### Bioacoustics

The advertisement calls of 14 males *Allobates gasconi* were analyzed, with eight presenting calls with 2–3 notes, three with calls containing 2–4 notes, two with calls consisting of 3–4 notes, and one individual with a call consisting of only 2 notes (each call emitted with one exhalation). Notes were repeated at a rate of 116.2 ± 33.5 notes/min (43.4 notes/min for two-note calls, 67.2 notes/min for three-note calls, and 5.5 notes/min for four-note calls). The duration of two-, three-, and four-note calls was 59 ± 4 ms, 98 ± 27 ms, and 150 ± 11 ms, respectively. The inter-call interval had a duration of 387 ± 154 ms (161–1,048 ms). The first and last notes had similar durations, with a note duration of 15 ± 4 ms (6–28 ms) and an inter-note interval of 30 ± 12 ms (19–42 ms).

Notes were frequency-modulated (Fig. 9). The first note was usually emitted with a lower amplitude than the last note; the average linear frequency modulation was 130 ± 192 Hz (-904–560 Hz). The first note had a low frequency of 5,373 ± 264 Hz (4,795–6,487 Hz), a high frequency of 5,810 ± 267 Hz (5,261–6,878 Hz) and a dominant frequency of 5,613 ± 270 Hz (5,082–6,761 Hz). The last note had a low frequency of 5,309 ± 168 Hz (5,007–5,643 Hz), a high frequency of 5,790 ± 155 Hz (5,461–6,059 Hz) and a dominant frequency of 5,573 ± 165 Hz (5,297–6,059 Hz). However, differences in spectral parameters between the first and last notes were not statistically significant, and frequencies are given as overall values in Table 3.

**Fig. 9.**
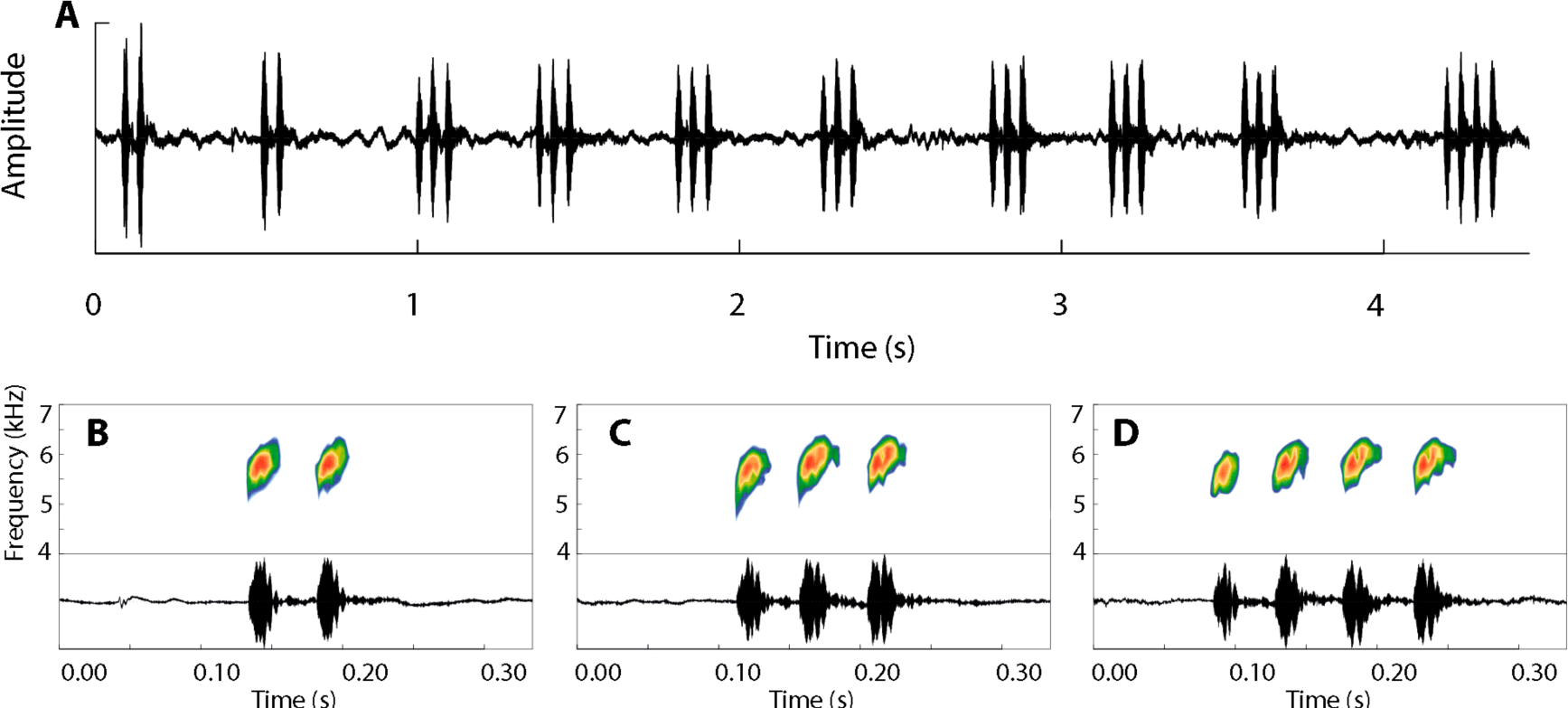
Advertisement call of *Allobates gasconi* from its type locality. (A) Oscillogram showing the continuous emission of 10 calls with distinct note arrangements (2–4 notes per call). Spectrograms and oscillograms of the three notes arrangements: (B) two-, (C), three- and (D) four notes (each call arrangements corresponds to an expiration). Illustration: A. S. Ferreira.

**Table 3.**
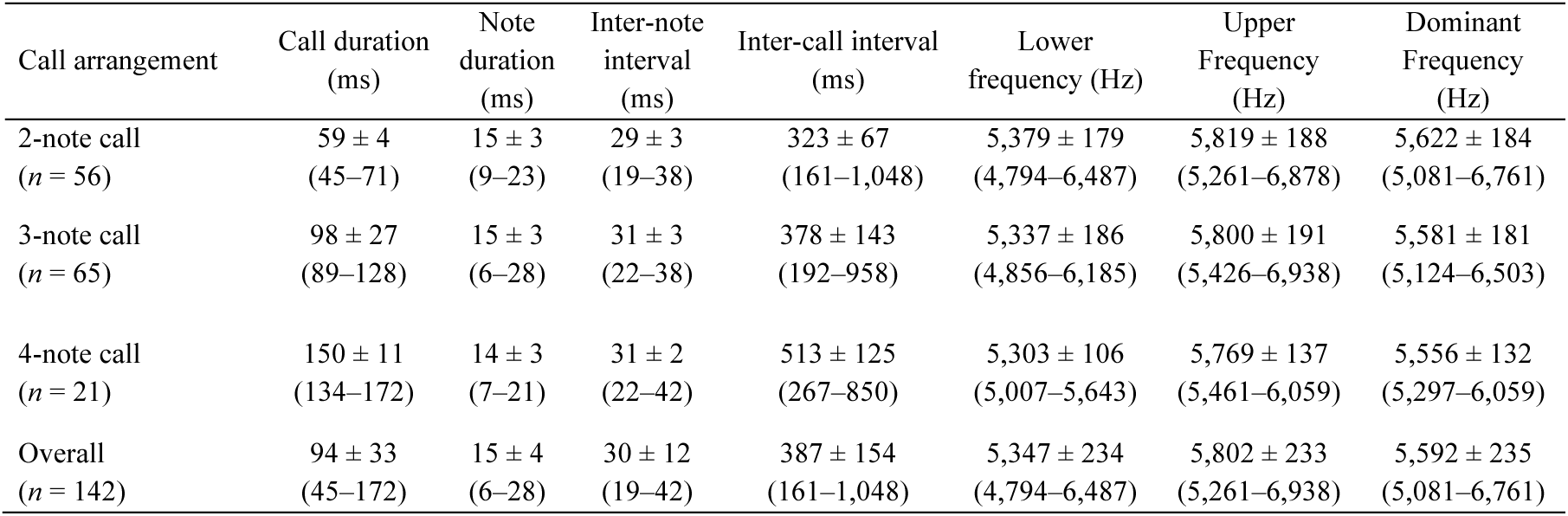
Advertisement call parameters of *Allobates gasconi* from topotypic localities in the Jurua River, Amazonas, Brazil. Values depict mean ± standard error and range. Abbreviations: *n*, sample size; ms, milliseconds; Hz, Hertz.

| Call arrangement | Call duration<br>(ms) | Note<br>duration<br>(ms) | Inter-note<br>interval<br>(ms) | Inter-call interval<br>(ms) | Lower<br>frequency (Hz) | Upper<br>Frequency<br>(Hz) | Dominant<br>Frequency<br>(Hz) |
| --- | --- | --- | --- | --- | --- | --- | --- |
| 2-note call<br>(*n* = 56) | 59 ± 4<br>(45–71) | 15 ± 3<br>(9–23) | 29 ± 3<br>(19–38) | 323 ± 67<br>(161–1,048) | 5,379 ± 179<br>(4,794–6,487) | 5,819 ± 188<br>(5,261–6,878) | 5,622 ± 184<br>(5,081–6,761) |
| 3-note call<br>(*n* = 65) | 98 ± 27<br>(89–128) | 15 ± 3<br>(6–28) | 31 ± 3<br>(22–38) | 378 ± 143<br>(192–958) | 5,337 ± 186<br>(4,856–6,185) | 5,800 ± 191<br>(5,426–6,938) | 5,581 ± 181<br>(5,124–6,503) |
| 4-note call<br>(*n* = 21) | 150 ± 11<br>(134–172) | 14 ± 3<br>(7–21) | 31 ± 2<br>(22–42) | 513 ± 125<br>(267–850) | 5,303 ± 106<br>(5,007–5,643) | 5,769 ± 137<br>(5,461–6,059) | 5,556 ± 132<br>(5,297–6,059) |
| Overall<br>(*n* = 142) | 94 ± 33<br>(45–172) | 15 ± 4<br>(6–28) | 30 ± 12<br>(19–42) | 387 ± 154<br>(161–1,048) | 5,347 ± 234<br>(4,794–6,487) | 5,802 ± 233<br>(5,261–6,938) | 5,592 ± 235<br>(5,081–6,761) |

#### Tadpole morphology

The following description is based on 14 tadpoles at Gosner stage 36 (lots APL 23698 and 23700) from *Jainú West* and *Altamira*. Table 4 summarizes 27 morphometric traits of 25 additional tadpoles at stages 34–41.

**Table 4.** Morphometric measurements (in mm) of tadpoles of *Allobates gasconi* from *Jainú West* (lots APL 23698 and 23703) and *Altamira* (lot APL 23700), middle Juruá River, Amazonas, Brazil. Values depict mean ± standard error and range. Acronyms are defined in the text. Abbreviation: *n*, sample size.

| Traits | Stages 34–35 (*n* = 7) | Stage 36 (*n* = 14) | Stages 40–41 (*n* = 4) |
| --- | --- | --- | --- |
| TMW | 1.8 ± 0.1 (1.6–1.9) | 1.9 ± 0.1 (1.5–2.0) | 2.2 ± 0.1 (2.1–2.2) |
| BW | 3.6 ± 0.3 (3.3–4.0) | 3.6 ± 0.3 (3.1–4.0) | 4.5 ± 0.2 (4.3–4.6) |
| HWLE | 3.3 ± 0.2 (3.1–3.6) | 3.3 ± 0.2 (3.1–3.9) | 3.6 ± 0.1 (3.4–3.6) |
| IOD | 2.1 ± 0.1 (2.0–2.2) | 2.1 ± 0.1 (2.0–2.2) | 2.3 ± 0.1 (2.2–2.4) |
| IND | 1.4 ± 0.1 (1.3–1.5) | 1.4 ± 0.1 (1.1–1.6) | 1.5 ± 0.0 (1.5–1.6) |
| BL | 5.4 ± 0.2 (5.1–5.7) | 5.7 ± 0.3 (5.3–6.0) | 6.1 ± 0.1 (6.0–6.1) |
| TAL | 12.4 ± 0.6 (11.5–13.3) | 13.0 ± 0.6 (11.5–14.0) | 16.0 ± 0.5 (15.3–16.5) |
| TL | 17.9 ± 0.8 (17.1–19.0) | 18.7 ± 0.7 (17.5–19.5) | 22.0 ± 0.5 (21.4–22.5) |
| SS | 4.1 ± 0.1 (4.0–4.2) | 4.1 ± 0.2 (3.9–4.4) | 4.4 ± 0.1 (4.3–4.5) |
| BH | 2.8 ± 0.1 (2.7–2.9) | 2.9 ± 0.1 (2.7–3.1) | 3.3 ± 0.1 (3.1–3.4) |
| TMH | 1.7 ± 0.0 (1.9–1.6) | 1.7 ± 0.1 (1.5–1.9) | 1.8 ± 0.1 (1.7–1.9) |
| MTH | 2.6 ± 0.1 (2.6–2.7) | 2.6 ± 0.1 (2.4–2.8) | 3.0 ± 0.1 (3.0–2.9) |
| ED | 0.9 ± 0.0 (0.9–1.0) | 0.9 ± 0.1 (0.8–1.0) | 1.0 ± 0.1 (1.0–1.1) |
| END | 0.9 ± 0.0 (0.8–0.9) | 0.9 ± 0.0 (0.8–1.0) | 1.0 ± 0.0 (1.0–1.0) |
| NSD | 0.7 ± 0.1 (0.5–0.9) | 0.8 ± 0.1 (0.6–0.9) | 0.5 ± 0.1 (0.5–0.6) |
| STL | 0.5 ± 0.0 (0.5–0.5) | 0.5 ± 0.0 (0.5–0.7) | 0.5 ± 0.1 (0.5–0.7) |
| VTL | 1.1 ± 0.1 (1.0–1.2) | 1.0 ± 0.1 (0.8–1.2) | 1.2 ± 0.1 (1.1–1.2) |
| ODW | 1.3 ± 0.1 (1.2–1.4) | 1.3 ± 0.1 (1.2–1.5) | 1.6 ± 0.1 (1.5–1.6) |
| PL | 0.3 ± 0.1 (0.2–0.4) | 0.4 ± 0.1 (0.3–0.4) | 0.2 ± 0.1 (0.2–0.3) |
| **AL** | 0.3 ± 0.0 (0.3–0.4) | 0.3 ± 0.1 (0.2–0.4) | 0.4 ± 0.1 (0.4–0.5) |
| **GAP** | 0.5 ± 0.0 (0.5–0.5) | 0.5 ± 0.0 (0.4–0.6) | - |
| **A1** | 0.9 ± 0.1 (0.8–1.0) | 0.9 ± 0.1 (0.7–1.1) | 0.9 ± 0.2 (0.6–1.1) |
| **A2** | 0.8 ± 0.0 (0.7–0.9) | 0.8 ± 0.1 (0.7–1.0) | - |
| **P1** | 0.8 ± 0.0 (0.8–0.8) | 0.8 ± 0.1 (0.7–0.9) | 1.0 ± 0.1 (0.9–1.1) |
| **P2** | 0.6 ± 0.0 (0.6–0.7) | 0.7 ± 0.1 (0.5–0.8) | 0.8 ± 0.1 (0.7–0.9) |
| **N° AL** | 4.0 ± 0.0 (4.0–4.0) | 4.0 ± 1.1 (4.0–4.0) | 9.8 ± 1.3 (8.0–11.0) |
| **N° PL** | 11.9 ± 2.3 (9.0–15.0) | 10.9 ± 3.2 (9.0–14.0) | 12.8 ± 1.7 (11.0–15.0) |

The body ovoid in dorsal view, rounded anteriorly but truncate posteriorly, and longer than wide (BL/BW = 157%; Fig. 10); it is flattened in lateral view (BH/BL = 64%). Body length is 31% of total length and head width 92% of body width. The snout is bluntly rounded in dorsal view but slightly pointed in lateral view; snout length slightly less than eye diameter (NSD/ED = 86%). Large eyes are positioned dorsally and directed laterally; eye diameter 96% of END; interorbital distance 151% of IND and 63% of HWLE. The small naris is located dorsally and directed anterolaterally, visible in dorsal and lateral views; internarial distance 42% of HWLE. The spiracle is sinistral, visible in dorsal, lateral and ventral views, easily seen at approximately mid-body and below lateral midline and directed transversely; inner wall free from body; spiracle-tube length 15% of HWLE. The gut is coiled and visible through the skin; its axis is directed to the left side of the body. The medial vent tube is dextral 1.0 mm, attached to ventral fin, right wall displaced dorsally; vent tube length 200% of spiracle length. Tail length is 69% of total length; tail maximum height 92% of BH; tail muscle robust and deeper than fins, tail muscle maximum height 63% of MTH. The dorsal fin emerges at the thigh-body insertion and reaches its maximum depth at the central portion of the tail; dorsal fin deeper than the ventral fin along the first two thirds of the tail. Tail tip acuminate, lacks flagellum.

**Fig. 10.**
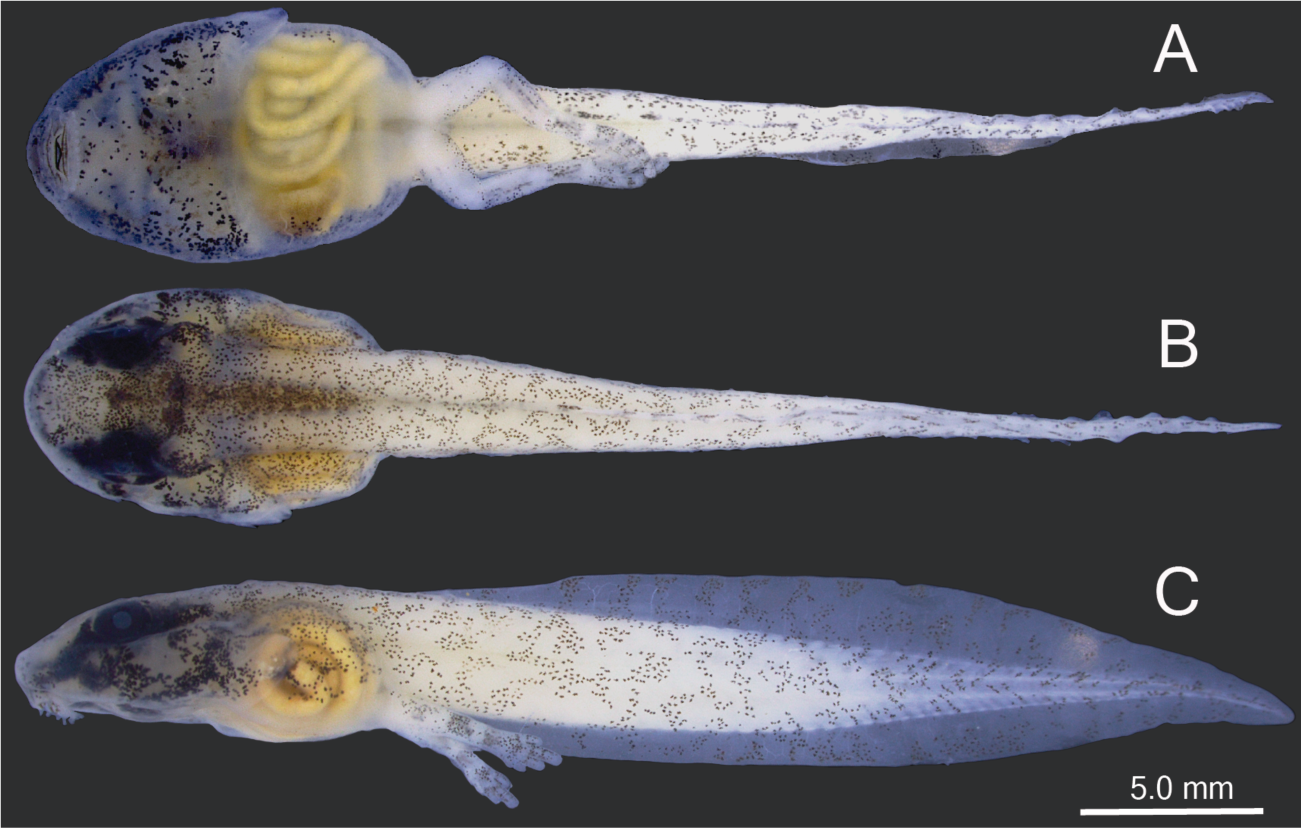
(A) Dorsal, (B) lateral and (C) ventral views of a preserved tadpole of *Allobates gasconi*, Gosner stage 36, from *Jainú West* (lot APL 23698). Photographs by A. S. Ferreira.

Oral disc is located anteroventrally and emarginated laterally (Fig. 11); oral disc width 36% of body width. Anterior labium has small pyramidal papillae only on lateral margin, 2 on each side. Posterior labium has a single marginal row of pyramidal papillae of variable length; short papillae are located close to the labium angle, all other papillae are elongate (Fig. 11); elongate papillae twice as long as the shorter ones. Submarginal papillae are absent. Upper jaw sheath is arch-shaped; lower jaw sheath “V”-shaped, deeper than upper sheath; cutting edge of each jaw sheath is serrated along its entire length. Labial keratodont row formula is (LKRF) 2(2)/2(1); tooth row A–1 complete; tooth row A–2 interrupted by a medial gap of ∼ 0.5 mm, each segment ∼ 0.4 mm long; tooth row P–1 interrupted by a small medial gap; P–1 slightly (0.2 mm) longer than P–2; P–3 absent. Vent tube and tooth row A-2 disappear at stage 41.

**Fig. 11.**
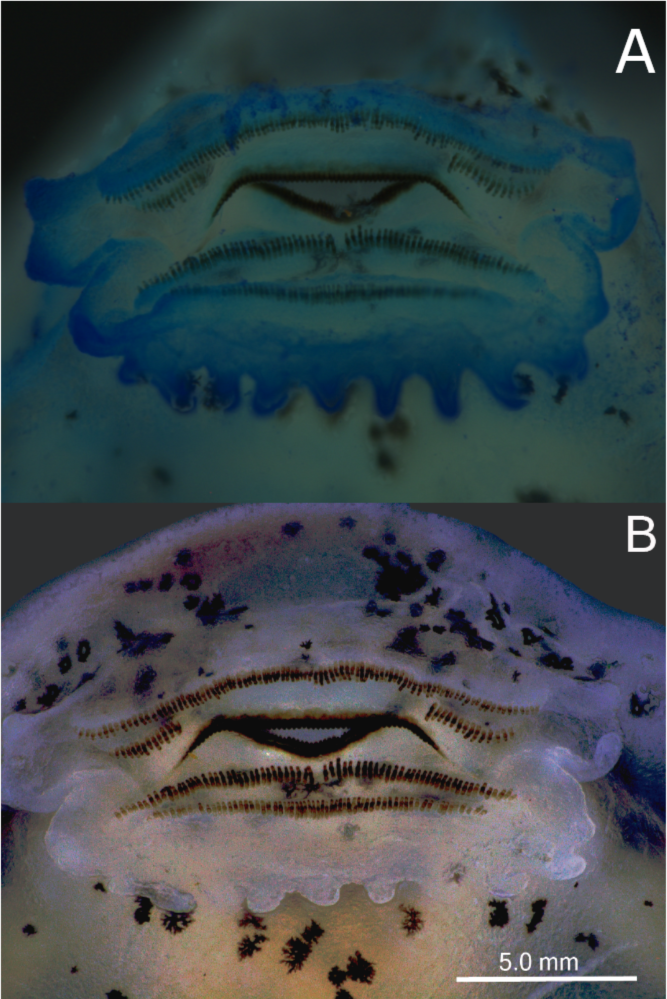
Detailed views of the oral disc of tadpoles of *Allobates gasconi*, Gosner stage 36, from *Jainú West*. (A) mouth dyed with blue dye and (B) mouth without dye. Photographs by A. S. Ferreira.

In life, dorsal and lateral surfaces of body are grayish brown with scattered irregular brown blotches. Ventral surface of body and tail musculature are the same as in preservative. At stage 36, fins are transparent with scattered irregular brown and silver blotches. In preservative, the dorsal skin of head and body is whitish-cream to yellowish-cream and covered with dark melanophores (Fig. 10). There is a higher concentration of melanophores in the center of the dorsum, but without an obvious hourglass-shaped mark. A dark brown dorsolateral stripe extends from the tip of the snout to the inguinal region in some specimens. Dark black blotches are present on the latero-frontal portion of head. The tail muscle is light cream and fins are translucent or pale with brown to dark brown blotches. Ventral surfaces are yellowish cream with brown irregular-shaped blotches surrounding the posterior labium until the beginning of pericardium and abdomen region; dark melanophores randomly distributed on posterior portion of ventral surface of head and tail; posterior region of ventral surface translucent and unpigmented, intestines visible through the skin.

#### Distribution and natural history notes

*Allobates gasconi* is a common diurnal, terrestrial frog that inhabits the leaf litter of seasonally flooded (*várzea*) forests along the middle and upper Juruá River in Brazil, and along the Yuyapichis River (affluent of the Pachita River, Ucayali Basin) in Peru. The specimen INPA-H 4889 from the paratype locality *Vai-Quem-Quer* (lower Juruá River) has morphological differences from the other individuals of the type series. In addition, we did not found *A. gasconi* in this locality, but there is another species with similar call and morphology (Ferrão & Ferreira, pers. comm.). Despite intense sampling effort in southwestern and western Brazilian Amazonia, *A. gasconi* has not been recorded in unflooded (*terra firme*) forests along the Juruá River or in unflooded or seasonally flooded forests along the Madeira, Purus and Solimões (upper Amazon) Rivers (unpublished data, A.P. Lima, M. Ferrão, A.S. Ferreira).

Breeding coincides with the rainy season from November to March. Males call while perched above the ground on shrub roots, the adaxial surface of live leaves and fallen logs, and within rolled leaves in the leaf litter. They are vocally most active at dawn (0530–0830 h) and dusk (1630–1800 h), and throughout the day on rainy days. Breeding males are territorial and exhibit aggressive behavior; in addition to responding to advertisement-call playback, two males were encountered in wrestling bouts. Territory size, however, remains unknown. When a female approaches a calling male, he emits low courtship calls in her direction, which attract her. We found clutches of 20–29 eggs deposited on the adaxial surface of live leaves of small shrubs and in dead rolled leaves in the leaf litter. Each oviposition site contained only one clutch. Eggs are enclosed by a layer of transparent jelly, which becomes cloudy during embryonic development (Fig. 12). Adults were not observed transporting tadpoles, nor were tadpoles found in temporary ponds, which leads us to hypothesize that tadpoles complete larval development in the flooding forest (*várzea*) of the Juruá River. Syntopic diurnal species include *Allobates femoralis*, *Ameerega trivittata* (Spix, 1824) and two probably undescribed species of *Allobates*.

**Fig. 12.**
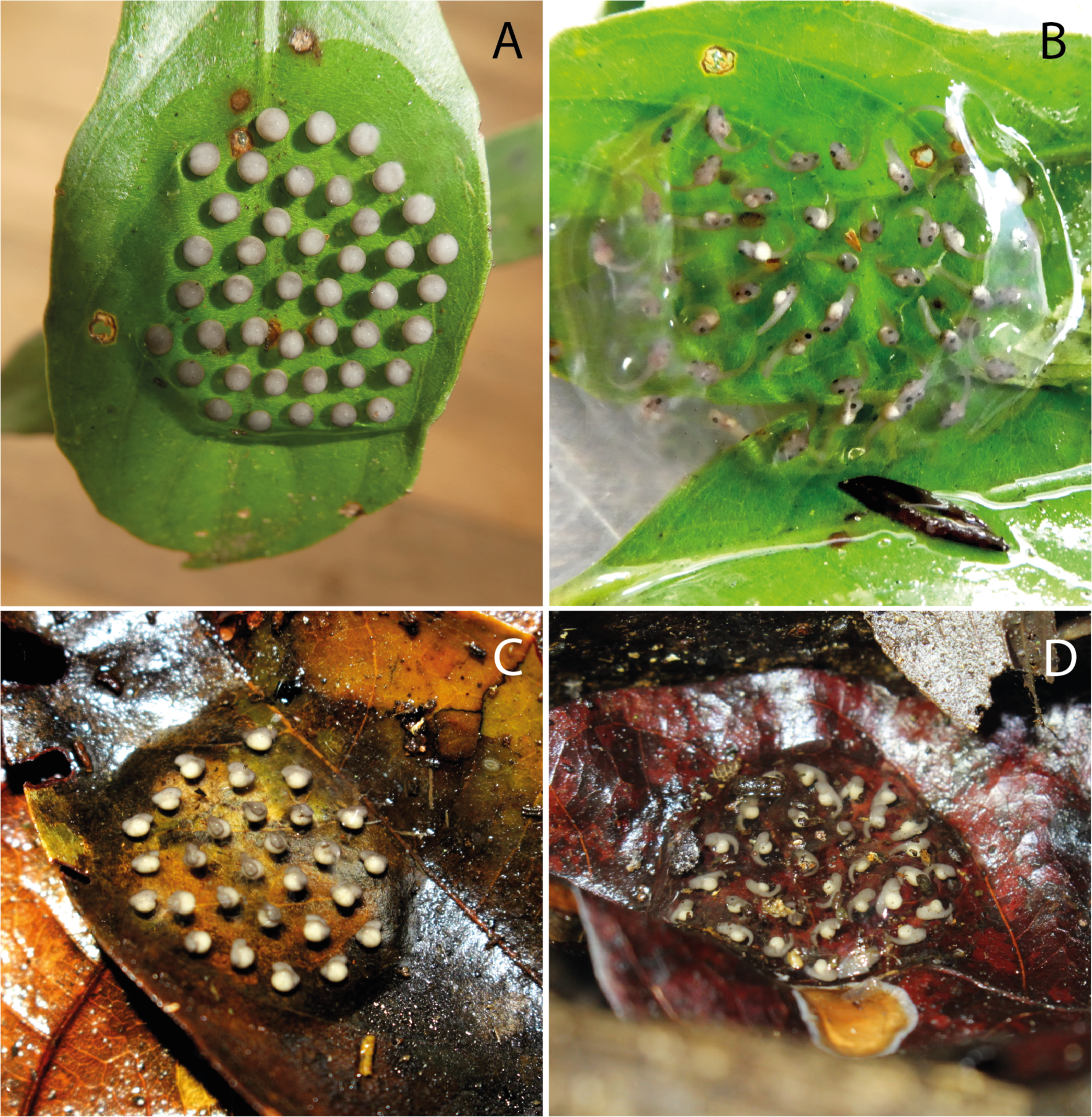
*Allobates gasconi* eggs and embryos from *Jainú West* and *Altamira*. (A) Freshly laid clutch on a green leaf contains 42 eggs, each enclosed within transparent jelly and with a pigmented animal pole. (B) Embryos of the clutch in A approximately 12 days after deposition. (C) Clutch laid on humid leaf containing 28 eggs approximately 3 days after deposition. (D) Clutch containing 31 embryos approximately 6 days after deposition. Photographs by A. S. Ferreira.

## Discussion

Uncertain taxonomy at the species level compromises the advance of knowledge in most fields of biology, but especially for the anuran biota of megadiverse regions. Several taxa have problematic taxonomic histories due to cryptic diversity, incomplete or unreliable species diagnoses, inexact type localities, the absence of genetic and acoustic data, poorly preserved type material or composite type series − and often a combination of these factors (Lima et al., 2009; Simões et al., 2013; Ferrão et al., 2020; Jaramillo et al., 2021; Fouquet et al., 2022). By redescribing the nurse frog *Allobates gasconi* through integrative taxonomy and by incorporating data from newly collected specimens from topotypic Amazonian localities, we show that the species has been characterized incorrectly, beginning with its oversimplified description two decades ago that made the diagnosis difficult (Morales, 2002), a circumstance that initiated a cascade of errors.

First, Victor Morales misidentified as *Allobates gasconi* a specimen from the Ituxí River (OMNH 36637, field number JPC 14139; Morales, 2002, pp. 58), which subsequently was included as a representative sample of that species in a phylogenetic analysis by Grant et al. (2006). Next, Lima et al. (2014) incorrectly identified as *A. gasconi* individuals from the Juruá River included in their phylogenetic tree, likely due to the short geographic distance between their sampling locality and the type locality of *A. gasconi*. Subsequently, Melo-Sampaio et al. (2018) reviewed the type series of *A. gasconi* and interpreted the observed phenotypic variation as population variation. More recently, Vacher et al. (2020) assigned specimens from the Madeira River to *A. gasconi*. Our phylogenetic inference shows that *A. gasconi* sensu stricto is not conspecific with individuals identified as *A. gasconi* by Grant et al. (2006), Lima et al. (2014), Melo-Sampaio et al. (2018) and Vacher et al. (2020). The initial misidentification, the lack of topotypic molecular and acoustic data, and the poor preservation of the holotype and several paratypes of the original type series of *A. gasconi* have all contributed to these misidentifications.

Several molecular phylogenetic studies have included DNA sequences of *Allobates gasconi* sensu lato (Grant et al., 2006; Lima et al., 2014; Lima et al., 2015; Grant et al., 2017; Melo-Sampaio et al., 2018; Simões et al., 2019; Lima et al., 2020; Réjaud et al., 2020; Souza et al., 2020; Jaramillo et al., 2021; Ferrão et al., 2022). In most of these studies, *A. gasconi* sensu lato was recovered as sister to the *A. tapajos* species complex and as a member of the *A. caeruleodactylus* clade (sensu Réjaud et al., 2020). In our phylogeny, topotypic individuals of *A. gasconi* nest with an unidentified specimen from Peru (voucher GGU684; Réjaud et al., 2020); we regard these specimens as conspecific because of the very low pairwise genetic distance between them. Also in our phylogeny, *Allobates gasconi* sensu stricto is recovered as sister to the Peruvian *Allobates* sp. “Huanuco” and groups within the large *A. trilineatus* clade.

Several species of nurse frogs widely distributed in Amazonia represent species complexes (Lima et al., 2020; Vacher et al., 2020; Réjaud et al., 2020; Ferrão et al., 2022). DNA barcoding of specimens identified as *Allobates gasconi* from Ituxí, Juruá and Madeira rivers delimited three OTUs, each locality harboring a distinct one (Vacher et al., 2020). As none of these specimens is conspecific with *A. gasconi* sensu stricto, this species complex is constituted entirely by unnamed species awaiting formal description. This level of cryptic diversity is likely even higher because additional specimens sequenced by Melo-Sampaio et al. (2018) formed a distinct lineage within the major clade representing the species complex but were not included in the barcoding analysis (Melo-Sampaio et al., 2018; present study). On the other hand, there is no molecular, morphological or acoustic evidence of additional cryptic diversity in *A. gasconi* sensu stricto.

*Allobates gasconi* is not the only problematic taxon among the eleven species of nurse frogs described by Morales (2002). *Allobates conspicuus*, for example, has a similar taxonomic history. Specimens assigned to *A. conspicuus* by Morales (2002) and subsequently included in the phylogenetic analysis by Grant et al. (2006) instead represent *A. subfolionidificans* (Melo-Sampaio et al., 2018); the phylogenetic relationships of *A. conspicuus* remain unknown. Melo-Sampaio et al. (2018) showed that the type series of *A. vanzolinius* and *A. fuscellus* contain specimens of different species; moreover, no additional specimens of *A. fuscellus* and *A. vanzolinius* have been reported since their description. Similarly, the type series of *A. sumtuosus* likely comprises specimens of more than one species because it groups with specimens from opposite sides of Amazonia (Simões et al., 2013), and because this taxon likely represents a species complex (Vacher et al., 2020; Fernandes et al., 2021). Further taxonomic revisions that integrate multiple and diverse data sources like those used in the present study could illuminate the complex taxonomic history of these taxa and help to better delimit those that are poorly diagnosed.

*Allobates gasconi* sensu stricto is known only from riparian environments subject to flood-pulse regimes of the middle and upper Juruá River in Brazil and the Ucayali River in Peru (Réjaud et al., 2020). Only two paratype localities (Condor and Nova Vida) on the upper Juruá River were not sampled in the present study, but it is expected that *A. gasconi* also occurs in forests under flood regime in this region. Moreover, the morphology of examined paratypes from these localities corroborates that of other confirmed individuals of *A. gasconi* examined here. Its putative distribution in the lower Juruá River is refuted, at least in the region from *Vai-Quem-Quer* to the river mouth; individuals in this region that resemble *A. gasconi* are not close relatives. The current geographic distribution of *A. gasconi* is disjunct: known Brazilian and Peruvian populations are distant ∼170 km from each other and are not connected by continuous flooded forests. However, extensive sampling in intervening areas is needed to determine if the species is continuously distributed and to assess its conservation status, as well as to better understand the species’ evolutionary history.

## Acknowledgements

We thank several local assistants for their practical help in the field. We are grateful for Lúcio H. Costa, pilot of the boat ‘Nadia’ for his dedication and good humor during the expedition. Collection permit N° 1337-1 was granted to A.P.L. by ICMBio.

## Appendix

Specimens examined

### Allobates bacurau

Adults. BRAZIL: AMAZONAS: Estrada do Miriti, Manicoré: INPA-H 35398 (holotype), 35397, 35399–35409 (paratypes).

### Allobates caldwellae

Adults. BRAZIL: AMAZONAS: Careiro: RAPELD M1 at km 32 of the BR-319 Highway: INPA-H 41047 (holotype), 41045–41046, 41049–41053, 41055–41057, 41059, 41061, 41063, 41067–41069 (paratopotypes); km 12 of the BR-319 Highway. INPA-H 41054, 41058, 41064, 41066 (paratypes); RAPELD M2, at km 100 of the BR-319 Highway: INPA-H 41048, 41060, 41062, 41065 (paratypes).

### Allobates femoralis

Adults. BRAZIL: PARÁ: Treviso: INPA-H 11657–11671, 15232, 30769–30778; Itaituba: INPA-H 26342–26354.

### Allobates fuscellus

Adults. BRAZIL: AMAZONAS: Ipixúna: Penedo, east bank of Juruá River: INPA-H 2532 (holotype), 2531 (paratopotype). Itamarati: Jainú, Juruá River: INPA-H 3114, 3250, 3270, 3514 (paratypes).

### Allobates hodli

Adults. BRAZIL: ACRE: Fazenda Experimental Catuaba: INPA-H 11621–11640 (paratypes); RONDÔNIA: Cachoeira do Jirau: INPA-H 16555 (holotype), 16541–16554, 16556–16569 (paratopotypes); Near Fortaleza do Abunã: INPA-H 16578, 16584–16587, 16589, 16591–92, 16597, 16602–03, 16605–16607, 16611–16614, 16620–16624, 16626, 16628, 16631, 16633, 16636–37, 16639–16641, 16643, 16645–46, 16648; Near Mutum-Paraná: INPA-H 16596, 16730, 16739, 16756, 16758, 16767, 16771, 16777–78, 16788, 16805, 16818–19 (paratypes). Tadpoles. BRAZIL: RONDÔNIA: Abunã: INPA-H 23693.

### Allobates marchesianus

Adults. BRAZIL: AMAZONAS: Missão Taracuá: INPA-H 7959–7990 (topotypes); São Gabriel da Cachoeira, 175 km E Missão Taracuá: INPA-H 7991, 7993, 8000–8007. Tadpoles. BRAZIL: AMAZONAS: Missão Taracuá: INPA-H 7943–7950, 7992, 7998, 8084 (topotypes).

### Allobates nidicola

Adults. BRAZIL: AMAZONAS: km 12 on road to Autazes: INPA-H 8093 (holotype), 7253–7259, 7261, 7262, 8094 (paratypes), INPA-H 28122, 28124, 28127, 28129, 28131, 28144, 28159, 28163, 28166, 28169, 28171, 28172, 28174, 28179, 28184, 28185 (topotypes). Tadpoles. BRAZIL: AMAZONAS: km 12 of road to *<u>Autazes:</u>* INPA-H 8021–8033, 8137–8139.

### Allobates paleovarzensis

Adults. BRAZIL: AMAZONAS: Careiro da Várzea: INPA-H 20904 (holotype), 20861–20903, 20905 (paratypes).

### Allobates subfolionidificans

Adults. BRAZIL: ACRE: Parque Zoobotânico of Universidade Federal do Acre: INPA-H 13760 (holotype), 11958–11974, 13749–13754, 13756–13759, 13761–62 (paratypes). Tadpoles. BRAZIL: ACRE: Parque Zoobotânico of Universidade Federal do Acre: INPA-H 14822–23.

### Allobates sumtuosus

Adults. BRAZIL: AMAZONAS: Reserva Florestal Adolpho Ducke: INPA-H 31949–31951; PARÁ: Reserva Biológica do Rio Trombetas: INPA-H 31952–56, 31958–60 (topotypes).

### Allobates tinae

Adults. BRAZIL: ACRE: Boca do Acre: INPA-H 40976, 41022, 41027, 41037, 41040; RONDÔNIA: Porto Velho, west bank of upper Madeira River: INPA-H 41012–21, 41029–36, 41041–44.

### Allobates velocicantus

Adults. BRAZIL: ACRE: Mâncio Lima: INPA-H 41342 (holotype), 41338–41341, 41343–41349 (paratopotypes); Cruzeiro do Sul: road connecting Cruzeiro do Sul to Guajará: INPA-H 41350 (paratype). Tadpoles: BRAZIL: ACRE: Mâncio Lima: INPA-H 41351.

### Allobates vanzolinius

Adults. BRAZIL: AMAZONAS: Vai-Quem-Quer, Rio Juruá. INPA-H 4896 (holotype), 4903, 4904, 4905, 4912 (paratypes); Jainú, Rio Juruá. INPA-H 3381, 3413 (paratypes).

## Availability of data and material

All new DNA sequences and call recordings have been deposited in public repositories.

## Declarations

### Ethics approval

Experiments have been conducted in accordance with relevant national legislation on the use of animals for research.

## Author Contributions

Anthony S. Ferreira, Miquéias Ferrão and Albertina P. Lima planned and organized the field expedition, conducted the specimen collection, analyzed the data, prepared figures and/or tables, authored or reviewed drafts of the article, and approved the final draft.

Miquéias Ferrão and Albertina P. Lima were also involved in obtaining funding. Antonio S. Cunha-Machado performed molecular data acquisition, authored or reviewed drafts of the article, and approved the final draft.

William E. Magnusson participated in data acquisition, authored or reviewed drafts of the article, and approved the final draft.

James Hanken prepared figures and/or tables, authored or reviewed drafts of the article, and approved the final draft.

## References

Altig, R., & McDiarmid, R. W. (1999). Body plan: development and morphology, (pp. 24–51). In: McDiarmid, R. W., & Altig, R. (Eds.), Tadpoles: the biology of anuran larvae. University of Chicago Press. Chicago.

Barrio-Amorós, C. L., & Santos, J. C. (2009). Description of a new *Allobates* (Anura, Dendrobatidae) from the eastern Andean piedmont, Venezuela. Phyllomedusa, 8, 89–104.

Bauer, L. (1986). A new genus and a new specific name in the dart poison frog family (Dendrobatidae, Anura, Amphibia). Ripa, 1–12.

Bibron, G. (1840). Historia Fisica, Politica y Natural de la Isla de Cuba, Segunda Part. Historia Natural (De la Sagra). Tomo VIII. Atlas de Zoología. Arthus Bertrand. Paris.

Bioacoustics Research Program. (2015). Raven Pro: interactive sound analysis software.v. 1.6. Ithaca: The Cornell Lab of Ornithology. Available at: http://www.birds.cornell.edu/raven.

Boulenger, G. A. (1884). On a collection of frogs from Yurimaguas, Huallaga River, Northern Peru. Proceedings of the Zoological Society of London, 1883, 635–638. DOI:10.1111/j.1469-7998.1883.tb06669.x

Caldwell, J. P., & Lima, A. P. (2003). A new Amazonian species of *Colostethus* (Anura: Dendrobatidae) with a nidicolous tadpole. Herpetologica, 59, 219–234. DOI:10.1655/0018-0831(2003)059[0219:ANASOC]2.0.CO;2

Cope, E. D. (1866). Fourth contribution to the herpetology of tropical America. Proceedings of the Academy of Natural Sciences of Philadelphia, 18, 123–132.

Fernandes, I. Y., Moraes, L. J. C. L., Menin, M., Farias, I. P., Lima, A. P., & Kaefer, I. L. (2021). Unlinking the speciation steps: geographical factors drive changes in sexual signals of an Amazonian nurse-frog through body size variation. Evolutionary Biology, 48, 81–93.

Ferrão, M., Hanken, J., & Lima, A. P. (2022). A new nurse frog of the *Allobates tapajos* species complex (Anura, Aromobatidae) from the upper Madeira River, Brazilian Amazonia. PeerJ, 10, e13751.

Fouquet, A., Peloso, P., Jairam, R., Lima, A. P., Mônico, A. T., Ernst, R., & Kok, P. J. R. (2022). Back from the deaf: integrative taxonomy revalidates an earless and mute species, *Hylodes grandoculis* van Lidth de Jeude, 1904, and confirms a new species of *Pristimantis* Jiménez de la Espada, 1870 (Anura: Strabomantidae) from the Eastern Guiana Shield. Organisms Diversity & Evolution, 22, 1065–1098 DOI:10.1007/s13127-022-00564-w

Gagliardi-Urrutia, G., Castroviejo-Fisher, S., Rojas-Runjaic, F. J. M., Jaramillo, A. F., Solís, S., & Simões, P. I. (2021). A new species of nurse-frog (Aromobatidae, *Allobates*) from the Amazonian forest of Loreto, Peru. Zootaxa, 5026, 375–404. DOI:10.11646/35ootaxa.5026.3.3

Gascon, C., Malcolm, J. R., Patton, J. L., Silva, M. N. F., Bogarti, J. P., Lougheed, S. et al. (2000). Riverine barriers and the geographic distribution of Amazonian species. Proceedings of the National Academy of Sciences of the United States of America, 97, 13672–13677.

Gosner, K. L. (1960). A simplified table for staging anuran embryos and larvae with notes on identification. Herpetologica, 16, 183–190.

Grant, T., Frost, D. F., Caldwell, J. P., Gagliardo, R., Haddad, C. F. B., Kok, P. J. R., et al. (2006). Phylogenetic systematics of dart-poison frogs and their relatives (Anura: Athesphatanura: Dendrobatidae). Bulletin of the American Museum of Natural History, 299, 1–262. DOI:10.1206/0003-0090(2006)299[1:PSODFA]2.0.CO;2

Grant, T., Rada, M., Anganoy-Criollo, M., Batista, A., Dias, P. H., Jeckel, A. M., et al. (2017). Phylogenetic systematics of dart-poison frogs and their relatives revisited (Anura: Dendrobatoidea). South American Journal of Herpetology, 12, 1–90. DOI:10.2994/SAJH-D-17-00017.1

Grant, T., & Rodríguez, L. O. (2001). Two new species of frogs of the genus *Colostethus* (Dendrobatidae) from Peru and a redescription of *C. trilineatus* (Boulenger, 1883). American Museum Novitates, 3355, 1–24.

Hoang, D. T., Chernomor, O., Haeseler, A. V., Minh, B. Q., & Vinh, L. S. (2018). UFBoot2: improving the ultrafast bootstrap approximation. Molecular Biology and Evolution, 35, 518–522. DOI:10.1093/molbev/msx281

Jaramillo, A. F., Gagliard-Urrutia, G., Simões, P. I. & Castroviejo-Fisher, S. (2021). Redescription and phylogenetics of *Allobates trilineatus* (Boulenger 1884 “1883”) (Anura: Aromobatidae) based on topotypic specimens. Zootaxa, 4951, 201–235. DOI:10.11646/35ootaxa.4951.2.1

Kaiser, H., Coloma, L. A., & Gray, H. M. (1994). A new species of *Colostethus* (Anura: Dendrobatidae) from Martinique, French Antilles. Herpetologica, 50, 23–32.

Kalyaanamoorthy, S., Minh, B. Q., Wong, T. K. F., von Haeseler, A., & Jermiin, L. S. (2017). ModelFinder: fast model selection for accurate phylogenetic estimates. Nature Methods, 4, 587–589. DOI:10.1038/nmeth.4285.

Kaplan, M. (1997). A new species of *Colostethus* from the Sierra Nevada de Santa Marta (Colombia) with comments on intergeneric relationships within the Dendrobatidae. Journal of Herpetology, 31, 369–375.

Katoh, K., & Standley, D. M. (2013). MAFFT Multiple sequence alignment software version 7: Improvements in performance and usability. Molecular Biology and Evolution, 30, 772–780. DOI:10.1093/molbev/mst010

Kearse, M., Moir, R., Wilson, A., Stones-Havas, S., Cheung, M., Sturrock, S., et al. (2012). Geneious Basic: an integrated and extendable desktop software platform for the organization and analysis of sequence data. Bioinformatics, 28, 1647–1649. DOI:10.1093/bioinformatics/bts199

Kimura, M. (1980). A simple method for estimating evolutionary rates of base substitutions through comparative studies of nucleotide sequences. Journal of Molecular Evolution, 16, 111–120. DOI:10.1007/bf01731581

Köhler, J., Jansen, M., Rodríguez, A., Kok, P. J. R., Toledo, L. F., Emmrich, M., et al. (2017). The use of bioacoustics in anuran taxonomy: theory, terminology, methods and recommendations for best practice. Zootaxa, 4251, 1–124. DOI:10.11646/36ootaxa.4251.1.1

Köhler, J., Castillo-Urbina, E., Aguilar-Puntriano, C., Vences, M., & Glaw, F. (2022). Rediscovery, redescription and identity of *Pristimantis nebulosus* (Henle, 1992), and description of a new terrestrial-breeding frog from montane rainforests of central Peru (Anura, Strabomantidae). Zoosystematics and Evolution, 98(2), 213–232. DOI:10.3897/zse.98.84963

Kok, P. J. R., Hölting, M., & Ernst, R. (2013). A third microendemic to the Iwokrama Mountains of central Guyana: a new “cryptic” species of *Allobates* Zimmerman and Zimmerman, 1988 (Anura: Aromobatidae). Organisms, Diversity & Evolution, 13, 621–638.

Kok, P. J. R., MacCulloch, R. D., Gaucher, P. Poelman, E. H., Bourne, G. R., Lathrop, A., & Lenglet, G. L. (2006). A new species of *Colostethus* (Anura, Dendrobatidae) from French Guiana with a redescription of *Colostethus beebei* (Noble, 1923) from its type locality. Phyllomedusa, 5, 43–66.

La Marca, E. (1992). Catalogo taxonomico, biogeografico y bibliografico de las ranas de Venezuela. Cuadernos Geográficos, 9, 1–197.

La Marca, E., Manzanilla, J., & Mijares-Urrutia, A. (2004). Revisión taxonómica del Colostethus del Norte de Venezuela confuindido durante largo tiempo con *C. brunneus*. Herpetotropicos, 1, 40–50.

Lima, A. P., Caldwell, J. P., Biavati, G., & Montanarin, A. (2010). A new species of *Allobates* (Anura: Aromobatidae) from Paleovárzea Forest in Amazonas, Brazil. Zootaxa, 2337, 1–17. DOI:10.11646/zootaxa.2337.1.1

Lima, A. P., Caldwell, J. P., & Strussmann, C. (2009) Redescription of *Allobates brunneus* (Cope) 1887 (Anura: Aromobatidae: Allobatinae), with a description of the tadpole, call, and reproductive behavior. Zootaxa, 1988, 1–16. DOI:10.11646/zootaxa.1988.1.1

Lima, A. P., Ferrão, M., & Silva, D. (2020). Not as widespread as thought: integrative taxonomy reveals cryptic diversity in the Amazonian nurse frog *Allobates tinae* Melo-Sampaio, Oliveira & Prates, 2018 and description of a new species. Journal of Zoological Systematics and Evolutionary Research, 58, 1173–1194. DOI:10.1111/jzs.12406

Lima, A. P., Sanchez, D. E. A., & Souza, J. R. D. (2007). A new Amazonian species of the frog genus *Colostethus* (Dendrobatidae) that lays its eggs on undersides of leaves. Copeia, 2007, 114–122. DOI:10.1643/00458511(2007)7[114:ANASOT] 2.0.CO;2

Lima, A. P., Simões, P. I. & Kaefer, I. L. (2014). A new species of *Allobates* (Anura: Aromobatidae) from the Tapajós River basin, Pará State, Brazil. Zootaxa, 3889, 355–387. DOI:10.11646/37ootaxa.3889.3.2

Lima, A. P., Simões, P. I., & Kaefer, I. L. (2015). A new species of *Allobates* (Anura: Aromobatidae) from Parque Nacional da Amazônia, Pará State, Brazil. Zootaxa, 3980, 501–525. DOI:10.11646/37ootaxa.3980.4.3

Lyra, M., Lourenço, A. C. C., Pinheiro, P. D. P., Pezzuti, T. L., Baêta, D., Barlow, A., et al. (2020). High-throughput DNA sequencing of museum specimens sheds light on the long-missing species of the *Bokermannohyla claresignata* group (Anura: Hylidae: Cophomantini). Zoological Journal of the Linnean Society. DOI:10.1093/zoolinnean/zlaa033

Maia, G. F., Lima, A. P., & Kaefer, I. L. (2017). Not just the river: genes, shapes, and sounds reveal population-structured diversification in the Amazonian frog *Allobates tapajos* (Dendrobatoidea). Biological Journal of the Linnean Society, 121, 95–108. DOI:10.1093/biolinnean/blw017

Melin, D. E. (1941). Contributions to the knowledge of the Amphibia of South America. Göteborgs Kungl. Vetenskaps-och Vitterhets-samhälles. Handlingar. Serien B, Matematiska och Naturvetenskapliga Skrifter, 1, 1–71.

Melo-Sampaio, P. R., Oliveira, R. M., & Prates, I. (2018). A new nurse frog from Brazil (Aromobatidae: *Allobates*) with data on the distribution and phenotypic variation of western Amazonian species. South American Journal of Herpetology, 13, 131–149. DOI:10.2994/SAJH-D-17-00098.1

Melo-Sampaio, P. R., Prates, I., Peloso, P. L. V., Recoder, R., Dal Vechio, F., Marques-Souza, S., & Rodrigues, M. T. (2020). A new nurse frog from Southwestern Amazonian highlands, with notes on the phylogenetic affinities of *Allobates alessandroi* (Aromobatidae). Journal of Natural History, 54, 43–62. DOI:10.1080/00222933.2020.1727972

Melo-Sampaio, P. R., Souza, M. B., & Peloso, P. L. V. (2013). A new riparian species of *Allobates* Zimmermann and Zimmermann, 1988 (Anura: Aromobatidae) from southwestern Amazonia. Zootaxa, 3716, 336–348. DOI:10.11646/38ootaxa.3716.3.2

Moraes, L. J. C. L., & Lima, A. P. (2021). A new nurse frog (Allobates, Aromobatidae) with acricket-like advertisement call from eastern Amazonia. Herpetologica, 77, 146–163. DOI:10.1655/Herpetologica-D-20-00010.1.

Moraes, L. J. C. L., Pavan, D., & Lima, A. P. (2019). A new nurse frog of *Allobates* masniger-nidicola complex (Anura, Aromobatidae) from the east bank of Tapajós River, eastern Amazonia. Zootaxa, 4648, 401–434. DOI:10.11646/zootaxa.4648.3.1.

Morales, V. R. (2002). Sistemática y biogeografia del grupo *trilineatus* (Amphibia, Anura, Dendrobatidae, Colostethus), com descripción de once nuevas. Publicaciones de la Asociación de Amigos Doñana, 13, 1–59.

Myers, C. W., Paolillo-O, A., & Daly, J. W. (1991). Discovery of a defensively malodorous and nocturnal frog in the family Dendrobatidae: Phylogenetic significance of a new genus and species from Venezuelan Andes. American Museum Novitates, 3002, 1–33.

Palumbi, S. R. (1996). Nucleic acids II: the polymerase chain reaction, (pp. 205–247). In: Hillis, D. M., Moritz, C., & Mable, B.K. (Eds.). Molecular systematics. Sunderland: Sinauer & Associates.

R Core Team. (2022). R: A language and environment for statistical computing. Vienna, Austria. Available at https://www.R-project.org/. Retrieved November 1, 2022.

Rancilhac, L., Bruy, T., Scherz, M. D., Pereira, E. A., Preick, M., Straube, N., et al. (2020). Target-enriched DNA sequencing from historical type material enables a partial revision of the Madagascar giant stream frogs (genus *Mantidactylus*). Journal of Natural History, 54, 1–4. DOI:10.1080/00222933.2020.1748243

Réjaud, A., Rodrigues, M. T., Crawford, A. J., Castroviejo-Fisher, S., Jaramillo, A. F., Chaparro, J. C., et al. (2020). Historical biogeography identifies a possible role of Miocene wetlands in the diversification of the Amazonian rocket frogs (Aromobatidae: Allobates). Journal of Biogeography, 47, 2472–2482. DOI:10.1111/jbi.13937

Rivero, J. A. (1980). Notas sobre los anfibios de Venezuela III. Nuevos *Colostethus* de los Andes Venezolanos. Memoria. Sociedad de Ciencias Naturales La Salle. Caracas, 38, 95–111.

Schulze, A., Jansen, M., & Köhler, G. (2015). Tadpole diversity of Bolivia’s lowland anuran communities: molecular identification, morphological characterization, and ecological assignment. Zootaxa, 4016, 1–111. DOI:10.11646/39ootaxa.4016.1.1

Shine, R. (1979). Sexual selection and sexual dimorphism in the Amphibia. Copeia, 1979, 297–306. DOI:10.2307/1443418

Simões, P. I. (2016). A new species of nurse-frog (Aromobatidae, Allobates) from the Madeira River basin with a small geographic range. Zootaxa, 4083, 501–525.

Simões, P. I., Gagliardi-Urrutia, L. A. G., Rojas-Runjaic, F. J. M., Castroviejo-Fisher, S. (2018). A new species of nurse-frog (Aromobatidae, *Allobates*) from the Juami River basin, northwestern Brazilian Amazonia. Zootaxa, 4387, 109–133.

Simões, P. I., Kaefer, I. L., Farias, I. P., & Lima, A. P. (2013). An integrative appraisal of the diagnosis and distribution of *Allobates sumtuosus* (Morales, 2002) (Anura, Aromobatidae). Zootaxa, 3746, 401–421. DOI:10.11646/39ootaxa.3746.3.1

Simões, P. I., Lima, A. P., & Farias, I. P. (2010). The description of a cryptic species related to the pan-Amazonian frog *Allobates femoralis* (Boulenger 1883) (Anura: Aromobatidae). Zootaxa, 2406, 1–28.

Simões, P. I., Rojas, D., & Lima, A. P. (2019). A name for the nurse-frog (*Allobates*, Aromobatidae) of Floresta Nacional de Carajás, Eastern Brazilian Amazonia. Zootaxa, 4550, 71–100. DOI:10.11646/40ootaxa.4550.1.3

Souza, J. R. D., Ferrão, M., Hanken, J., & Lima, A. P. (2020). A new nurse frog (Anura: *Allobates*) from Brazilian Amazonia with a remarkably fast multi-noted advertisement call. PeerJ, 8, e9979. DOI:10.7717/peerj.9979

Sueur, J., Aubin, T., & Simonis, C. (2008). Seewave, a free modular tool for sound analysis and synthesis. Bioacoustics, 18, 213–226. DOI:10.1080/09524622.2008.9753600

Tamura, K., Stecher, G., Peterson, D., Filipski, A., & Kumar, S. (2021). MEGA11: molecular evolutionary genetics analysis version 6.0. Molecular Biology and Evolution, 30, 2725–2729. DOI:10.1093/molbev/mst197

Trifinopoulos, J., Nguyen, L. T., von Haeseler, A., & Minh, B. Q. (2016). W-IQ-TREE: a fast online phylogenetic tool for maximum likelihood analysis. Nucleic Acids Research, 44, 232–235. DOI:10.1093/nar/gkw256

Vacher, J-P., Chave, J., Ficetola, F. G., Sommeria-Klein, G., Tao, S., Thébaud, C., et al. (2020). Large-scale DNA-based survey of frogs in Amazonia suggests a vast underestimation of species richness and endemism. Journal of Biogeography, 47, 1781–1791. DOI:10.1111/jbi.1

Wagler, J. (1830). Natürliches System der Amphibien, mit vorangehender Classification der Säugthiere und Vogel. Ein Beitrag zur vergleichenden Zoologie. München, Stuttgart and Tübingen: J.G. Cotta.

Zhang, J., Kapli, P., Pavlidis, P., & Stamatakis, A. (2013). A general species delimitation method with applications to phylogenetic placements. Bioinformatics, 29, 2869–2876.

Zimmerman, H., & Zimmerman, E. (1988). Etho-Taxonomie und zoogeographische Artengruppenbildung bei Pfeilgiftfröschen (Anura: Dendrobatidae). Salamandra, 24, 125–160.

