## Supplementtal files for "Integrative reappraisal of the Amazonian nurse frog *Allobates gasconi* (Morales 2002) based on topotypical data, with implications for the systematics and taxonomy of a large species complex"

### Supplementary files

**Table S1.** Museum voucher and GenBank accession number of specimens used in phylogenetic analyses. Accession numbers in bold font denote sequences generated by this study.

| Species | Voucher | 12S | 16S | ND1 | COI | Cytb |
|---|---|---|---|---|---|---|
| *Allobates* aff. *granti* | 125PG | JN690205 | JN690931 | | | |
| *Allobates* aff. *magnussoni* | 977126 | MT627173 | MT627173 | MT627173 | MT627173 | MT627173 |
| *Allobates* aff. *melanolaemus* | MTR28013 | MT627203 | MT627203 | MT627203 | MT627203 | MT627203 |
| *Allobates* aff. *olfersioides 1* | MTR17821 | | KDQF01003353 | | | |
| *Allobates* aff. *olfersioides 2* | MTR16435 | MT627202 | MT627202 | MT627202 | MT627202 | MT627202 |
| *Allobates* aff. *olfersioides 3* | JFT959 | | KDQF01002701 | | | |
| *Allobates* aff. *tapajos 1* | MTR10084 | MT627197 | MT627197 | MT627197 | MT627197 | MT627197 |
| *Allobates* aff. *tapajos 2* | AF1906 | MT627175 | MT627175 | MT627175 | MT627175 | MT627175 |
| *Allobates* aff. *tinae 1* | MPEG13397 | DQ502213 | DQ502213 | | DQ502900 | DQ502648 |
| *Allobates* aff. *trilineatus 1* | FGZC3247 | MT627185 | MT627185 | MT627185 | MT627185 | MT627185 |
| *Allobates* aff. *trilineatus 2* | JMP2313 | MT627195 | MT627195 | MT627195 | MT627195 | MT627195 |
| *Allobates* aff. *undulatus* | AMNHA159139 | DQ283044 | DQ283044 | | DQ502756 | DQ502459 |
| *Allobates algorei* | TNHCFS5551 | HQ290950 | HQ290950 | HQ290950 | | HQ290530 |
| *Allobates amissibilis* | PK3798 | MT627204 | MT627204 | MT627204 | MT627204 | MT627204 |
| *Allobates bacurau* | INPAH35401 | | KU195701 | | | |
| *Allobates caeruleodactylus* | MTR10227 | MT627199 | MT627199 | MT627199 | MT627199 | MT627199 |
| *Allobates caldwellae* | MPEG13826 | DQ502099 | DQ502099 | | | DQ502531 |
| *Allobates carajas* | BM163 | MT627183 | MT627183 | MT627183 | MT627183 | MT627183 |
| *Allobates chalcopis* | Alca1 | MT627182 | MT627182 | MT627182 | MT627182 | |
| *Allobates conspicuus/subfolionidificans* | FGZC3279 | MT627186 | MT627186 | MT627186 | MT627186 | MT627186 |
| *Allobates crombiei* | AF1097 | MT627174 | MT627174 | MT627174 | MT627174 | MT627174 |
| *Allobates femoralis* | AF3224 | MT627179 | MT627179 | MT627179 | MT627179 | MT627179 |

| | | | | | | |
|---|---|---|---|---|---|---|
| *Allobates femoralis* SS | AfemShucv3a | DQ523001 | DQ523072 | | | DQ523142 |
| *Allobates flaviventris* | HJ545 | MT627192 | MT627192 | MT627192 | MT627192 | MT627192 |
| *Allobates fratisenescus* | QCAZ54377 | | MF624172 | | | MF614174 |
| *Allobates goianus* | SAMA8574 | MT627207 | MT627207 | MT627207 | MT627207 | MT627207 |
| *Allobates granti* | AF1998 | MT627176 | MT627176 | MT627176 | MT627176 | MT627176 |
| *Allobates grillicantus* | MPEG43046 | | MW220039 | | | |
| *Allobates grillisimilis* | MTR12749 | MT627200 | MT627200 | MT627200 | MT627200 | MT627200 |
| *Allobates hodli* | ABU2194 | | KX044279 | | | |
| *Allobates humilis/pittieri* | CVULA5690 | KJ940454 | KJ940454 | | | |
| *Allobates insperatus/juami* | JMP1703 | MT627193 | MT627193 | MT627193 | MT627193 | MT627193 |
| *Allobates juanii/ranoides* | ARA2394 | DQ502271 | DQ502271 | | DQ502933 | DQ502702 |
| *Allobates kamilae* | HJ285 | MT627189 | MT627189 | MT627189 | MT627189 | MT627189 |
| *Allobates kingsburyi* | QCAZ16523 | AY364549 | HQ290963 | | | HQ290541 |
| *Allobates magnussoni* | BM168 | MT627184 | MT627184 | MT627184 | MT627184 | MT627184 |
| *Allobates marchesianus* | AJC2498 | MT627180 | MT627180 | MT627180 | MT627180 | MT627180 |
| *Allobates masniger* | MTR10155 | MT627198 | MT627198 | MT627198 | MT627198 | MT627198 |
| *Allobates nidicola* | MPEG13821 | DQ502101 | DQ502101 | | | DQ502533 |
| *Allobates melanolaemus* | NMP6V711404 | | MT524148 | | | |
| *Allobates niputidea* | MUJ3520 | DQ502272 | DQ502272 | | DQ502934 | DQ502703 |
| *Allobates nunciatus* | MPEG36777 | MT627196 | MT627196 | MT627196 | MT627196 | MT627196 |
| *Allobates olfersioides* | MNRJ79897 | MF624178 | MF624178 | | | MF614175 |
| *Allobates ornatus* | MHNSM22863 | | EU342550 | | | |
| *Allobates pacaas* | MZUSP158938 | | MT076999 | | | |
| *Allobates paleovarzensis* | JMP2196 | MT627194 | MT627194 | MT627194 | MT627194 | |
| *Allobates* sp. Huanuco | FGZC3348 | MT627187 | MT627187 | MT627187 | MT627187 | MT627187 |
| *Allobates* sp. Neblina | MTR15537 | MT627201 | MT627201 | MT627201 | MT627201 | MT627201 |

| | | | | | | |
|---|---|---|---|---|---|---|
| *Allobates* sp. Ucuyali | GGU684 | | MT524137 | | | |
| *Allobates sumtuosus* | AF2212 | MT627177 | MT627177 | MT627177 | MT627177 | MT627177 |
| *Allobates sieggreenae* | MCP14533 | | MW293942 | | | |
| *Allobates talamancae* | QCAZ35236 | MT627205 | MT627205 | MT627205 | MT627205 | MT627205 |
| *Allobates tapajos* | MJH3973 | DQ502110 | DQ502110 | | DQ502820 | DQ502542 |
| *Allobates tinae* | HJ298 | MT627190 | MT627190 | MT627190 | MT627190 | MT627190 |
| *Allobates trilineatus* | AF4493 | | MT524111 | | | |
| *Allobates undulatus* | AJC3040 | MT627181 | MT627181 | MT627181 | MT627181 | MT627181 |
| *Allobates velocicantus* | MCP10187/88 | MF624181 | MF624181 | | | MF614178 |
| *Allobates zaparo* | USNM546405 | DQ502026 | DQ502026 | | DQ502752 | DQ502455 |
| *Ameerega hahneli* | AF2673 | MT627178 | MT627178 | MT627178 | MT627178 | MT627178 |
| *Anomaloglossus stepheni* | MJH3928 | DQ502107 | DQ502107 | | DQ502818 | DQ502539 |
| *Aromobates saltuensis/nocturnus* | TNHCFS5541/AMNHA130042 | HQ290970 | HQ290970 | HQ290970 | DQ502860 | DQ502592 |
| *Leucostethus fugax* | QCAZ16513 | HQ290958 | HQ290958 | HQ290958 | | HQ290538 |
| *Colostethus brachistriatus* | CZPDUV4603 | MF624204 | MF624204 | | MF614304 | MF614198 |
| *Dendrobates auratus* | MVZHerp149723 | JX564862 | JX564862 | JX564862 | JX564862 | JX564862 |
| *Epipedobates boulengeri* | UMMZ227952/QCAZ16574 | HQ290997 | HQ290997 | HQ290997 | DQ502742 | DQ502447 |
| *Mannophryne collaris* | FS5523 | MT627188 | MT627188 | MT627188 | MT627188 | MT627188 |
| *Phyllobates terribilis* | TNHC64420/AMNHA118566 | HQ291006 | HQ291006 | HQ291006 | DQ502861 | DQ502593 |
| *Rheobates palmatus* | RHEOPALM | MT627206 | MT627206 | MT627206 | MT627206 | MT627206 |
| *Silverstoneia nubicola/erasmios* | TNHCFS4942/MAR336 | HQ290966 | HQ290966 | HQ290966 | MF614333 | MF614237 |
| *Allobates gasconi* SL1 | APL14410 | | KJ747333 | | | |
| *Allobates gasconi* SL1 | APL14411 | | KJ747334 | | | |
| *Allobates gasconi* SL1 | APL14416 | | KJ747335 | | | |
| *Allobates gasconi* SL2 | APL23940 | | **OQ297604** | | | |

| | | | | | | |
|---|---|---|---|---|---|---|
| *Allobates gasconi* SL2 | APL24058 | | **OQ297605** | | | |
| *Allobates gasconi* SL2 | APL24068 | | **OQ297606** | | | |
| *Allobates gasconi* SL2 | APL24070 | | **OQ297607** | | | |
| *Allobates gasconi* SL2 | APL24071 | | **OQ297608** | | | |
| *Allobates gasconi* SL3 | HJ480 | MT627191 | MT627191 | MT627191 | MT627191 | MT627191 |
| *Allobates gasconi* SL3 | HJ299 | | KDQF01002640 | | | |
| *Allobates gasconi* SL4 | MCP13630 | | KY886577 | | | KY886618 |
| *Allobates gasconi* SL4 | MNRJ91665 | | KY886576 | | | KY886617 |
| *Allobates gasconi* SL4 | MNRJ91679 | | KY886578 | | | KY886619 |
| *Allobates gasconi* SL4 | MNRJ91683 | | KY886574 | | | KY886615 |
| *Allobates gasconi* SL4 | MNRJ91684 | | KY886575 | | | KY886616 |
| *Allobates gasconi* SL5 | MNRJ 91681 | | KY886572 | | | |
| *Allobates gasconi* SL5 | MNRJ 91682 | | KY886573 | | | KY886614 |
| *Allobates gasconi* SL5 | OMNH36636 | DQ502209 | DQ502209 | | DQ502898 | DQ502644 |
| *Allobates gasconi* SL5 | MPEG13003 | DQ502052 | DQ502052 | | DQ502777 | DQ502483 |
| *Allobates gasconi* SL5 | MNRJ 90229 | | KY886570 | | | KY886612 |
| *Allobates gasconi* SL5 | MNRJ 90230 | | KY886571 | | | KY886613 |
| *Allobates gasconi* sensu stricto | APL23345 | | **ON997555** | | | |
| *Allobates gasconi* sensu stricto | APL23342 | | **ON997554** | | | |
| *Allobates gasconi* sensu stricto | APL23346 | | **ON997547** | | | |
| *Allobates gasconi* sensu stricto | APL23397 | | **ON997551** | | | |
| *Allobates gasconi* sensu stricto | APL23336 | | **ON997561** | | | |
| *Allobates gasconi* sensu stricto | APL23337 | | **ON997553** | | | |
| *Allobates gasconi* sensu stricto | APL23338 | | **ON997560** | | | |
| *Allobates gasconi* sensu stricto | APL23698 | | **ON997550** | | | |
| *Allobates gasconi* sensu stricto | APL23354 | | **ON997552** | | | |

| | | |
|---|---|---|
| *Allobates gasconi* sensu stricto | APL23355 | **ON997562** |
| *Allobates gasconi* sensu stricto | APL23356 | **ON997563** |
| *Allobates gasconi* sensu stricto | APL23357 | **ON997549** |
| *Allobates gasconi* sensu stricto | APL23360 | **ON997559** |
| *Allobates gasconi* sensu stricto | APL23411 | **ON997558** |
| *Allobates gasconi* sensu stricto | APL23418 | **ON997557** |
| *Allobates gasconi* sensu stricto | APL23422 | **ON997548** |
| *Allobates gasconi* sensu stricto | APL23433 | **ON997556** |
| *Allobates gasconi* sensu stricto | APL23436 | **ON997564** |
| *Allobates gasconi* sensu stricto | APL23703 | **ON997546** |
| *Allobates gasconi* sensu stricto | APL23700 | **ON997545** |

**Table S2**. Morphometric measurements (in mm) of adults of the type series and recently collected specimens of *A. gasconi* sensu stricto (SS). Measurement acronyms are defined in the text. Abbreviations: INPAH, museum voucher catalog numbers; FN, field numbers; SP, species; TS, *A. gasconi* from type series; RC, *A. gasconi* rerenttly collected; M, male; F, female.

| INPAH/FN | SP | Sex | SVL | HL | HW | SL | EN | IN | EL | IO | TYM | FAL | UAL | HANDI | HANDII | HANDIII | HANDIV | WFD | TL | FL | THL | DPT | WTT | WTD | WPF |
|---|---|---|---|---|---|---|---|---|---|---|---|---|---|---|---|---|---|---|---|---|---|---|---|---|---|
| 3083 | TS | M | 15.2 | 4.0 | 4.2 | 2.3 | 1.6 | 2.2 | 2.0 | 4.1 | 0.6 | 3.4 | 3.8 | 2.6 | 2.8 | 3.8 | 2.5 | 0.4 | 7.5 | 6.9 | 6.5 | 0.3 | 0.3 | 0.3 | 0.3 |
| 3093 | TS | M | 14.8 | 4.3 | 4.4 | 2.1 | 1.4 | 2.2 | 2.0 | 3.7 | 0.8 | 3.5 | 3.6 | 2.4 | 2.7 | 3.6 | 2.2 | 0.4 | 7.0 | 7.1 | 7.2 | 0.4 | 0.3 | 0.5 | 0.3 |
| 3249 | TS | M | 15.5 | 5.0 | 4.8 | 2.2 | 1.6 | 2.4 | 2.0 | 4.9 | 0.7 | 2.3 | 3.6 | 3.0 | 2.7 | 3.6 | 2.3 | 0.5 | 6.5 | 6.5 | 7.1 | 0.6 | 0.3 | 0.5 | 0.4 |
| 3513 | TS | M | 15.7 | 4.3 | 4.8 | 2.1 | 1.4 | 2.3 | 2.1 | 4.2 | 0.8 | 2.9 | 3.1 | 2.4 | 2.5 | 3.6 | 2.3 | 0.5 | 7.5 | 6.5 | 7.1 | 0.6 | 0.4 | 0.3 | 0.6 |
| 3541 | TS | M | 15.6 | 4.2 | 4.5 | 2.3 | 1.5 | 2.3 | 2.1 | 4.1 | 0.6 | 3.6 | 3.4 | 2.3 | 2.5 | 3.3 | 2.0 | 0.4 | 6.8 | 6.5 | 7.1 | 0.5 | 0.3 | 0.3 | 0.3 |
| 3512 | TS | M | 15.9 | 4.5 | 5.0 | 2.5 | 1.4 | 2.2 | 2.0 | 4.4 | 0.6 | 3.1 | 3.4 | 2.8 | 2.8 | 3.8 | 2.6 | 0.5 | 6.6 | 6.0 | 7.7 | 0.5 | 0.3 | 0.6 | 0.4 |
| 3055 | TS | F | 16.0 | 4.2 | 4.0 | 2.2 | 1.6 | 2.1 | 2.1 | 4.0 | 0.8 | 3.2 | 3.6 | 2.8 | 2.6 | 3.6 | 2.2 | 0.4 | 7.2 | 6.5 | 6.6 | 0.5 | 0.4 | 0.3 | 0.3 |
| 3079 | TS | F | 16.7 | 4.5 | 4.7 | 2.1 | 1.6 | 2.5 | 2.2 | 4.6 | 0.8 | 3.6 | 3.8 | 2.5 | 2.6 | 3.8 | 2.4 | 0.4 | 7.5 | 7.2 | 7.5 | 0.4 | 0.3 | 0.6 | 0.3 |
| 3491 | TS | F | 17.1 | 4.3 | 4.6 | 2.2 | 1.6 | 2.4 | 2.0 | 4.5 | 0.8 | 3.4 | 3.6 | 2.4 | 2.4 | 3.5 | 2.1 | 0.4 | 7.0 | 7.0 | 7.3 | 0.4 | 0.4 | 0.6 | 0.3 |
| 23336 | RC | M | 16.4 | 4.6 | 4.9 | 2.6 | 1.9 | 2.5 | 2.2 | 4.8 | 0.9 | 3.6 | 4.1 | 2.9 | 2.9 | 4.1 | 2.4 | 0.6 | 8.0 | 7.4 | 7.7 | 0.6 | 0.5 | 0.6 | 0.5 |
| 23337 | RC | M | 16.3 | 4.8 | 4.9 | 2.5 | 2.0 | 2.4 | 2.3 | 4.5 | 0.9 | 3.8 | 4.1 | 2.8 | 2.5 | 3.7 | 2.4 | 0.5 | 7.2 | 7.3 | 7.1 | 0.7 | 0.5 | 0.7 | 0.4 |
| 23338 | RC | M | 17.0 | 4.5 | 5.0 | 2.5 | 1.9 | 2.5 | 2.2 | 4.7 | 0.9 | 3.6 | 3.8 | 2.9 | 2.6 | 4.1 | 2.6 | 0.6 | 7.0 | 7.5 | 7.2 | 0.6 | 0.6 | 0.7 | 0.5 |
| 23339 | RC | M | 15.5 | 4.5 | 4.7 | 2.5 | 1.8 | 2.4 | 2.1 | 4.4 | 0.8 | 3.3 | 3.8 | 2.8 | 2.6 | 3.6 | 2.2 | 0.5 | 7.3 | 7.2 | 7.1 | 0.5 | 0.3 | 0.6 | 0.4 |
| 23341 | RC | M | 14.8 | 4.7 | 4.8 | 2.4 | 1.9 | 2.4 | 2.0 | 4.4 | 0.8 | 3.3 | 3.8 | 2.7 | 2.5 | 3.6 | 2.0 | 0.6 | 7.2 | 6.6 | 6.6 | 0.6 | 0.3 | 0.5 | 0.5 |
| 23342 | RC | M | 16.0 | 4.4 | 4.9 | 2.5 | 1.9 | 2.7 | 2.2 | 4.5 | 0.8 | 4.4 | 4.5 | 3.3 | 2.9 | 4.3 | 2.6 | 0.6 | 8.2 | 7.3 | 8.1 | 0.6 | 0.4 | 0.6 | 0.4 |
| 23343 | RC | M | 15.9 | 4.3 | 4.9 | 2.5 | 1.8 | 2.3 | 2.2 | 4.5 | 0.9 | 3.8 | 4.1 | 3.0 | 2.8 | 4.3 | 2.4 | 0.5 | 7.3 | 7.2 | 7.4 | 0.6 | 0.7 | 0.7 | 0.4 |
| 23344 | RC | M | 15.9 | 4.4 | 4.8 | 2.8 | 2.0 | 2.4 | 2.0 | 4.6 | 0.8 | 3.3 | 3.5 | 2.8 | 2.6 | 3.6 | 2.2 | 0.5 | 7.0 | 6.6 | 6.5 | 0.6 | 0.4 | 0.6 | 0.5 |
| 23349 | RC | M | 16.4 | 4.2 | 4.7 | 2.5 | 2.0 | 2.4 | 2.2 | 4.7 | 0.9 | 3.8 | 3.9 | 2.8 | 2.6 | 3.8 | 2.5 | 0.5 | 6.6 | 6.7 | 6.8 | 0.5 | 0.6 | 0.7 | 0.4 |
| 23351 | RC | M | 16.2 | 4.5 | 4.5 | 2.5 | 1.9 | 2.4 | 2.1 | 4.4 | 0.8 | 3.7 | 4.1 | 2.8 | 2.9 | 4.1 | 2.8 | 0.6 | 7.4 | 7.0 | 6.6 | 0.5 | 0.5 | 0.7 | 0.4 |
| 23352 | RC | M | 16.1 | 4.5 | 4.8 | 2.4 | 1.7 | 2.5 | 2.1 | 4.8 | 0.8 | 3.9 | 4.1 | 3.0 | 2.8 | 4.1 | 2.7 | 0.5 | 7.6 | 7.7 | 7.3 | 0.5 | 0.5 | 0.7 | 0.3 |
| 23354 | RC | M | 16.4 | 4.7 | 5.2 | 2.8 | 2.0 | 2.4 | 2.2 | 4.5 | 0.8 | 2.9 | 3.8 | 2.8 | 2.7 | 3.9 | 2.6 | 0.4 | 7.8 | 7.7 | 7.2 | 0.6 | 0.5 | 0.6 | 0.4 |
| 23355 | RC | M | 16.4 | 4.5 | 5.2 | 2.3 | 1.9 | 2.4 | 2.2 | 4.8 | 0.9 | 3.9 | 3.8 | 3.1 | 3.0 | 4.2 | 2.8 | 0.5 | 7.6 | 8.1 | 7.5 | 0.5 | 0.4 | 0.6 | 0.4 |

| | | | | | | | | | | | | | | | | | | | | | | | | | |
|---|---|---|---|---|---|---|---|---|---|---|---|---|---|---|---|---|---|---|---|---|---|---|---|---|---|
| 23356 | RC | M | 17.0 | 4.8 | 5.0 | 2.5 | 1.9 | 2.4 | 2.1 | 4.5 | 0.8 | 3.9 | 4.0 | 2.9 | 2.9 | 4.2 | 2.9 | 0.6 | 7.7 | 7.1 | 7.7 | 0.6 | 0.6 | 0.6 | 0.4 |
| 23357 | RC | M | 16.5 | 4.2 | 5.0 | 2.6 | 1.9 | 2.6 | 2.4 | 4.7 | 0.8 | 3.8 | 3.9 | 2.9 | 2.6 | 3.7 | 2.7 | 0.4 | 7.6 | 7.2 | 7.1 | 0.5 | 0.4 | 0.7 | 0.5 |
| 23359 | RC | M | 17.1 | 5.7 | 6.0 | 2.5 | 2.1 | 2.6 | 2.2 | 5.3 | 0.7 | 3.9 | 4.4 | 3.1 | 3.1 | 4.4 | 3.1 | 0.5 | 9.2 | 8.9 | 8.4 | 0.7 | 0.5 | 0.7 | 0.4 |
| 23360 | RC | M | 16.3 | 4.6 | 4.7 | 2.8 | 1.9 | 2.3 | 2.0 | 4.5 | 0.8 | 3.6 | 4.1 | 2.8 | 2.4 | 3.8 | 2.4 | 0.4 | 7.0 | 7.5 | 6.9 | 0.6 | 0.4 | 0.6 | 0.4 |
| 23411 | RC | M | 15.8 | 5.5 | 5.3 | 2.8 | 1.9 | 2.6 | 2.4 | 5.5 | 0.8 | 3.8 | 4.1 | 3.0 | 3.1 | 4.1 | 3.0 | 0.6 | 8.7 | 8.0 | 8.8 | 0.8 | 0.5 | 0.6 | 0.4 |
| 23418 | RC | M | 15.4 | 4.7 | 4.9 | 2.6 | 2.0 | 2.3 | 2.1 | 4.9 | 0.8 | 3.8 | 4.2 | 2.9 | 2.8 | 3.8 | 2.4 | 0.5 | 7.8 | 7.0 | 7.5 | 0.5 | 0.3 | 0.7 | 0.3 |
| 23419 | RC | M | 15.7 | 4.2 | 4.7 | 2.1 | 1.4 | 2.3 | 2.0 | 4.7 | 0.7 | 3.8 | 3.8 | 2.9 | 2.9 | 4.1 | 2.8 | 0.5 | 7.6 | 7.4 | 7.5 | 0.7 | 0.4 | 0.5 | 0.4 |
| 23422 | RC | M | 16.5 | 4.6 | 5.2 | 2.9 | 2.0 | 2.3 | 2.2 | 5.0 | 0.6 | 3.6 | 3.8 | 2.9 | 2.8 | 3.9 | 2.6 | 0.5 | 9.0 | 8.2 | 8.4 | 0.4 | 0.5 | 0.6 | 0.4 |
| 23433 | RC | M | 15.3 | 5.0 | 4.9 | 2.2 | 1.6 | 2.2 | 2.2 | 4.9 | 0.9 | 3.3 | 4.1 | 2.9 | 2.8 | 3.9 | 2.8 | 0.4 | 7.9 | 7.5 | 7.7 | 0.4 | 0.3 | 0.6 | 0.3 |
| 23436 | RC | M | 17.0 | 4.0 | 5.2 | 2.8 | 1.6 | 2.3 | 1.9 | 4.3 | 0.8 | 3.6 | 3.8 | 2.4 | 2.5 | 3.4 | 2.3 | 0.6 | 7.6 | 7.0 | 7.0 | 0.4 | 0.3 | 0.6 | 0.3 |
| 23397 | RC | M | 16.0 | 4.6 | 4.3 | 2.4 | 1.9 | 2.3 | 2.2 | 4.5 | 0.9 | 3.5 | 3.8 | 2.7 | 2.6 | 3.6 | 2.5 | 0.4 | 7.3 | 6.7 | 7.1 | 0.6 | 0.6 | 0.6 | 0.4 |
| 23345 | RC | F | 16.7 | 4.3 | 4.9 | 2.8 | 1.9 | 2.6 | 2.1 | 4.3 | 0.8 | 3.6 | 3.8 | 2.8 | 2.7 | 3.8 | 2.5 | 0.6 | 7.6 | 7.5 | 7.3 | 0.6 | 0.5 | 0.8 | 0.3 |
| 23346 | RC | F | 16.0 | 4.7 | 4.8 | 2.8 | 1.8 | 2.4 | 2.1 | 4.6 | 0.8 | 3.7 | 3.8 | 2.9 | 2.7 | 3.9 | 2.6 | 0.6 | 7.4 | 7.0 | 7.0 | 0.5 | 0.5 | 0.7 | 0.4 |
| 23347 | RC | F | 16.2 | 4.9 | 4.5 | 2.6 | 2.0 | 2.4 | 2.3 | 4.6 | 0.9 | 3.5 | 4.1 | 2.8 | 2.6 | 3.9 | 2.5 | 0.5 | 7.3 | 7.4 | 7.0 | 0.6 | 0.7 | 0.7 | 0.3 |
| 23348 | RC | F | 16.5 | 4.9 | 4.7 | 2.8 | 1.9 | 2.6 | 2.3 | 4.7 | 0.9 | 3.6 | 3.8 | 2.9 | 2.9 | 3.9 | 2.8 | 0.6 | 7.2 | 7.1 | 7.2 | 0.6 | 0.3 | 0.7 | 0.3 |
| 23350 | RC | F | 18.0 | 4.5 | 5.1 | 2.5 | 1.9 | 2.5 | 2.3 | 4.9 | 0.9 | 4.3 | 4.4 | 3.1 | 2.9 | 4.2 | 2.7 | 0.5 | 8.1 | 8.1 | 7.8 | 0.6 | 0.6 | 0.8 | 0.3 |
| 23358 | RC | F | 18.0 | 4.4 | 5.0 | 2.8 | 2.0 | 2.5 | 2.5 | 4.8 | 0.9 | 3.6 | 4.1 | 3.2 | 2.9 | 4.1 | 2.8 | 0.5 | 8.0 | 7.6 | 7.7 | 0.5 | 0.4 | 0.7 | 0.3 |
| 23410 | RC | F | 16.4 | 4.9 | 5.1 | 2.6 | 1.8 | 2.3 | 2.2 | 4.9 | 0.6 | 3.8 | 3.8 | 3.2 | 3.0 | 4.1 | 2.8 | 0.6 | 8.0 | 8.0 | 7.5 | 0.5 | 0.3 | 0.6 | 0.3 |

**Table S3**. Morphometric measurements (in mm) of tadpoles of *A. gasconi* sensu stricto (SS). Abbreviations: FN, field number; GS, Gosner stage. Measurement acronyms are described in the text.

| FN | GS | TMW | BW | HWLE | IOD | IND | BL | TAL | TL | SS | BH | TMH | MTH | ED | END | NSD | STL | VTL | ODW | PL | AL | GAP | A1 | A2 | P1 | P2 |
|---|---|---|---|---|---|---|---|---|---|---|---|---|---|---|---|---|---|---|---|---|---|---|---|---|---|---|
| 23698 | 36 | 1.9 | 3.5 | 3.2 | 2.2 | 0.4 | 5.5 | 12.5 | 18.1 | 4.2 | 2.8 | 1.5 | 2.7 | 0.8 | 0.9 | 0.8 | 0.5 | 0.9 | 1.2 | 0.4 | 0.3 | - | 0.7 | 0.7 | 0.8 | 0.6 |
| 23698 | 36 | 1.9 | 3.5 | 3.3 | 2.2 | 0.4 | 5.9 | 13.1 | 19.0 | 4.2 | 2.8 | 1.5 | 2.6 | 0.9 | 0.8 | 0.9 | 0.5 | 0.9 | 1.2 | 0.3 | 0.2 | - | 0.7 | 0.7 | 0.7 | 0.5 |
| 23698 | 36 | 2.0 | 3.7 | 3.2 | 2.2 | 0.4 | 6.0 | 13.5 | 19.5 | 4.4 | 3.0 | 1.5 | 2.6 | 0.9 | 0.9 | 0.9 | 0.5 | 0.8 | 1.2 | 0.4 | 0.3 | - | 0.8 | 0.8 | 0.9 | 0.6 |
| 23698 | 36 | 1.5 | 3.2 | 3.1 | 2.1 | 0.4 | 6.0 | 11.5 | 17.5 | 4.0 | 2.8 | 1.5 | 2.5 | 0.8 | 0.9 | 0.8 | 0.5 | 0.8 | 1.3 | 0.4 | 0.3 | 0.4 | 0.8 | 0.8 | 0.8 | 0.7 |
| 23700 | 36 | 1.9 | 3.7 | 3.3 | 2.1 | 0.4 | 5.3 | 13.0 | 18.3 | 4.0 | 2.8 | 1.7 | 2.7 | 0.9 | 0.9 | 0.7 | 0.5 | 1.1 | 1.2 | 0.3 | 0.3 | 0.5 | 0.9 | 0.8 | 0.8 | 0.6 |
| 23700 | 36 | 1.9 | 3.1 | 3.2 | 2.0 | 0.4 | 5.5 | 12.5 | 18.0 | 4.0 | 2.7 | 1.7 | 2.6 | 0.8 | 0.9 | 0.6 | 0.5 | 1.1 | 1.3 | 0.4 | 0.3 | 0.5 | 0.9 | 0.8 | 0.8 | 0.6 |
| 23700 | 36 | 1.9 | 3.7 | 3.4 | 2.1 | 0.4 | 5.9 | 13.5 | 19.4 | 4.1 | 2.8 | 1.9 | 2.6 | 0.9 | 0.9 | 0.8 | 0.5 | 1.1 | 1.4 | 0.3 | 0.3 | 0.6 | 0.9 | 0.8 | 0.9 | 0.8 |
| 23700 | 36 | 1.8 | 3.8 | 3.5 | 2.1 | 0.4 | 5.7 | 13.0 | 18.7 | 4.1 | 2.8 | 1.8 | 2.6 | 0.9 | 0.9 | 0.8 | 0.5 | 1.1 | 1.4 | 0.3 | 0.3 | 0.5 | 1.0 | 0.9 | 0.8 | 0.7 |
| 23700 | 36 | 1.8 | 3.9 | 3.3 | 2.1 | 0.4 | 5.9 | 13.1 | 19.0 | 4.1 | 2.7 | 1.7 | 2.4 | 0.9 | 0.9 | 0.6 | 0.7 | 1.1 | 1.3 | 0.3 | 0.4 | 0.5 | 0.9 | 0.8 | 0.8 | 0.8 |
| 23700 | 36 | 2.0 | 3.9 | 3.9 | 2.1 | 0.4 | 5.9 | 14.0 | 19.5 | 4.2 | 3.0 | 1.7 | 2.7 | 0.9 | 0.9 | 0.8 | 0.5 | 1.0 | 1.5 | 0.4 | 0.3 | 0.6 | 1.1 | 1.0 | 0.9 | 0.8 |
| 23700 | 36 | 1.9 | 3.5 | 3.2 | 2.1 | 0.4 | 5.0 | 12.5 | 17.5 | 3.9 | 2.8 | 1.6 | 2.6 | 1.0 | 0.9 | 0.6 | 0.5 | 1.0 | 1.2 | 0.3 | 0.4 | 0.5 | 1.0 | 0.9 | 0.8 | 0.7 |
| 23700 | 36 | 1.9 | 3.7 | 3.3 | 2.1 | 0.4 | 5.6 | 13.4 | 19.0 | 4.1 | 3.1 | 1.7 | 2.7 | 0.9 | 0.9 | 0.7 | 0.5 | 1.1 | 1.4 | 0.4 | 0.4 | 0.6 | 0.9 | 0.8 | 0.8 | 0.6 |
| 23700 | 36 | 1.8 | 4.0 | 3.5 | 2.2 | 0.4 | 6.0 | 13.4 | 19.4 | 4.4 | 2.9 | 1.7 | 2.8 | 1.0 | 0.9 | 0.7 | 0.5 | 1.0 | 1.3 | 0.3 | 0.3 | 0.6 | 0.9 | 0.8 | 0.9 | 0.8 |
| 23700 | 36 | 1.8 | 3.8 | 3.4 | 2.1 | 0.4 | 5.8 | 12.5 | 18.3 | 4.3 | 3.0 | 1.7 | 2.7 | 0.9 | 1.0 | 0.8 | 0.5 | 1.2 | 1.4 | 0.4 | 0.3 | 0.5 | 0.8 | 0.8 | 0.9 | 0.7 |
| 23700 | 35 | 1.9 | 4.0 | 3.3 | 2.0 | 0.4 | 5.6 | 13.0 | 18.6 | 4.2 | 2.9 | 1.9 | 2.7 | 0.9 | 0.9 | 0.6 | 0.5 | 1.2 | 1.4 | 0.4 | 0.3 | 0.5 | 0.9 | 0.9 | 0.8 | 0.7 |
| 23700 | 35 | 1.8 | 3.7 | 3.2 | 2.1 | 0.4 | 5.5 | 12.2 | 17.7 | 4.1 | 2.7 | 1.7 | 2.6 | 0.9 | 0.9 | 0.9 | 0.5 | 1.1 | 1.4 | 0.4 | 0.3 | 0.5 | 1.0 | 0.8 | 0.8 | 0.7 |
| 23700 | 34 | 1.9 | 3.5 | 3.3 | 2.1 | 0.4 | 5.3 | 11.5 | 16.8 | 4.0 | 2.7 | 1.7 | 2.6 | 0.9 | 0.9 | 0.7 | 0.5 | 1.1 | 1.2 | 0.2 | 0.3 | 0.5 | 0.9 | 0.8 | 0.8 | 0.6 |
| 23700 | 35 | 1.9 | 3.9 | 3.6 | 2.2 | 0.4 | 5.7 | 13.3 | 19.0 | 4.1 | 2.9 | 1.7 | 2.7 | 1.0 | 0.9 | 0.8 | 0.5 | 1.2 | 1.3 | 0.2 | 0.3 | 0.5 | 1.0 | 0.9 | 0.8 | 0.6 |
| 23700 | 34 | 1.6 | 3.3 | 3.1 | 2.0 | 0.4 | 5.1 | 12.0 | 17.1 | 4.1 | 2.7 | 1.7 | 2.5 | 0.9 | 0.9 | 0.5 | 0.5 | 1.2 | 1.3 | 0.4 | 0.3 | 0.5 | 0.8 | 0.7 | 0.8 | 0.7 |
| 23700 | 35 | 1.9 | 3.7 | 3.4 | 2.1 | 0.4 | 5.6 | 12.5 | 18.1 | 4.1 | 2.9 | 1.8 | 2.6 | 0.9 | 0.9 | 0.6 | 0.5 | 1.1 | 1.2 | 0.2 | 0.3 | 0.5 | 0.8 | 0.8 | 0.8 | 0.6 |
| 23700 | 34 | 1.8 | 3.3 | 3.2 | 2.0 | 0.4 | 5.3 | 12.5 | 17.8 | 4.0 | 2.7 | 1.6 | 2.5 | 0.9 | 0.8 | 0.8 | 0.5 | 1.0 | 1.4 | 0.4 | 0.4 | 0.5 | 0.9 | 0.8 | 0.8 | 0.6 |
| 23703 | 40 | 2.2 | 4.6 | 3.6 | 2.4 | 1.5 | 6.0 | 16.5 | 22.5 | 4.5 | 3.2 | 1.7 | 2.9 | 1.1 | 1.0 | 0.5 | 0.5 | 1.1 | 1.5 | 0.2 | 0.4 | 0.6 | 1.1 | 1.0 | 0.9 | 0.7 |
| 23703 | 41 | 2.3 | 4.6 | 3.6 | 2.3 | 1.6 | 6.1 | 15.3 | 21.4 | 4.4 | 3.3 | 1.8 | 3.0 | 1.0 | 1.0 | 0.6 | 0.7 | - | 1.6 | 0.2 | 0.4 | - | 0.6 | - | 1.1 | 0.8 |

| | | | | | | | | | | | | | | | | | | | | | | | | | | |
|---|---|---|---|---|---|---|---|---|---|---|---|---|---|---|---|---|---|---|---|---|---|---|---|---|---|---|
| 23703 | 40 | 2.3 | 4.3 | 3.6 | 2.2 | 1.5 | 6.0 | 16.3 | 22.3 | 4.3 | 3.4 | 1.8 | 3.0 | 1.0 | 1.0 | 0.5 | 0.5 | 1.2 | 1.5 | 0.3 | 0.4 | - | 0.9 | - | 0.9 | 0.7 |
| 23703 | 41 | 2.1 | 4.4 | 3.4 | 2.4 | 1.5 | 6.1 | 15.8 | 21.9 | 4.3 | 3.1 | 1.9 | 3.0 | 1.0 | 1.0 | 0.5 | 0.5 | - | 1.6 | 0.3 | 0.5 | - | 1.0 | - | 1.0 | 0.9 |

**Table S4**. Acoustic parameters of the pulsed notes of *A. gasconi* sensu stricto (SS). Abbreviations: FN, field number; FNJV, call voucher recordings; AT, air temperature (ºC); CD, call duration (ms); NN, number of notes; ND, note duration (ms), INI, inter-note interval (ms); ICI, inter-call interval (ms); LF, lower frequency (Hz); UF, upper frequency (Hz); DF, dominant frequency (Hz); CR, call rater.

| FN | FNJV | AT | CD | NN | ND1 | ND2 | INI1 | INI2 | ICI | LF1 | LF2 | UF1 | UF2 | DF1 | DF2 | CR |
|---|---|---|---|---|---|---|---|---|---|---|---|---|---|---|---|---|
| 23354 | 58775 | 26.5 | 0.103 | 3 | 0.015 | 0.015 | 0.029 | 0.03 | 0.414 | 4978.4 | 5271.3 | 5509.5 | 5699.1 | 5297.2 | 5512.5 | 120 |
| 23354 | 58775 | 26.5 | 0.103 | 3 | 0.016 | 0.014 | 0.027 | 0.03 | 0.423 | 5128.2 | 5274.2 | 5514.1 | 5722.7 | 5297.2 | 5512.5 | 120 |
| 23354 | 58775 | 26.5 | 0.102 | 3 | 0.017 | 0.013 | 0.024 | 0.03 | 0.495 | 5138.6 | 5279.4 | 5472.4 | 5733.1 | 5297.2 | 5512.5 | 120 |
| 23354 | 58775 | 26.5 | 0.105 | 3 | 0.015 | 0.02 | 0.025 | 0.027 | 0.302 | 5143 | 5252.5 | 5487.2 | 5487.3 | 5297.2 | 5555.6 | 120 |
| 23354 | 58775 | 26.5 | 0.108 | 3 | 0.013 | 0.019 | 0.029 | 0.026 | 0.469 | 5112.1 | 5411.5 | 5550.2 | 5781.8 | 5297.2 | 5598.6 | 120 |
| 23354 | 58775 | 26.5 | 0.061 | 2 | 0.017 | 0.016 | 0.028 | - | 0.203 | 5127.4 | 5253.2 | 5575.9 | 5722.6 | 5575.9 | 5555.6 | 120 |
| 23354 | 58775 | 26.5 | 0.058 | 2 | 0.019 | 0.016 | 0.023 | - | 0.367 | 5023.8 | 5279.3 | 5534.9 | 5722.6 | 5297.2 | 5555.6 | 120 |
| 23354 | 58775 | 26.5 | 0.058 | 2 | 0.018 | 0.016 | 0.024 | - | 0.212 | 5133.3 | 5341.9 | 5534.9 | 5743.5 | 5297.2 | 5555.6 | 120 |
| 23354 | 58775 | 26.5 | 0.058 | 2 | 0.017 | 0.018 | 0.023 | - | 0.267 | 5133.3 | 5279.3 | 5540.1 | 5759.1 | 5340.2 | 5598.6 | 120 |
| 23354 | 58775 | 26.5 | 0.063 | 2 | 0.015 | 0.022 | 0.026 | - | 0.398 | 5122.9 | 5399.3 | 5503.6 | 5748.7 | 5297.2 | 5598.6 | 120 |
| 23359 | 58776 | 25.8 | 0.147 | 4 | 0.013 | 0.013 | 0.033 | 0.03 | 0.389 | 5283.1 | 5527.5 | 5767.5 | 5920.2 | 5469.4 | 5727.8 | 34 |
| 23359 | 58776 | 25.8 | 0.147 | 4 | 0.017 | 0.016 | 0.024 | 0.029 | 0.735 | 5104.3 | 5303.2 | 5751.9 | 5792.7 | 5426.4 | 5469.4 | 34 |
| 23359 | 58776 | 25.8 | 0.145 | 4 | 0.014 | 0.016 | 0.03 | 0.029 | 0.496 | 5114.5 | 5277.7 | 5731.5 | 5853.9 | 5426.4 | 5598.6 | 34 |
| 23359 | 58776 | 25.8 | 0.144 | 4 | 0.011 | 0.016 | 0.029 | 0.031 | 0.498 | 5196.1 | 5514.5 | 5787.6 | 5932.7 | 5469.4 | 5727.8 | 34 |
| 23359 | 58776 | 25.8 | 0.146 | 4 | 0.014 | 0.018 | 0.028 | 0.022 | 0.443 | 5137.2 | 5483.6 | 5769.5 | 5906.9 | 5469.4 | 5684.8 | 34 |
| 23359 | 58776 | 25.8 | 0.098 | 3 | 0.011 | 0.015 | 0.029 | 0.029 | 0.441 | 5138.2 | 5249.9 | 5772.8 | 5758.4 | 5340.2 | 5598.6 | 34 |
| 23359 | 58776 | 25.8 | 0.101 | 3 | 0.015 | 0.018 | 0.026 | 0.028 | 0.276 | 5305.1 | 5527 | 5725.9 | 5894.1 | 5469.4 | 5727.8 | 34 |
| 23359 | 58776 | 25.8 | 0.103 | 3 | 0.012 | 0.018 | 0.029 | 0.028 | 0.394 | 5149.6 | 5476 | 5741.2 | 5889 | 5426.4 | 5727.8 | 34 |
| 23359 | 58776 | 25.8 | 0.104 | 3 | 0.015 | 0.012 | 0.028 | 0.032 | 0.518 | 5159.8 | 5453 | 5736.1 | 5906.9 | 5426.4 | 5641.7 | 34 |
| 23359 | 58776 | 25.8 | 0.098 | 3 | 0.011 | 0.015 | 0.029 | 0.029 | 0.441 | 5138.2 | 5249.9 | 5772.8 | 5758.4 | 5340.2 | 5598.6 | 34 |
| 23355 | 58777 | 25.8 | 0.057 | 2 | 0.012 | 0.012 | 0.033 | - | 0.347 | 4794.6 | 5065.6 | 5365.4 | 5416 | 5168 | 5254.1 | 122 |

| | | | | | | | | | | | | | | | | |
|---|---|---|---|---|---|---|---|---|---|---|---|---|---|---|---|---|
| 23355 | 58777 | 25.8 | 0.059 | 2 | 0.012 | 0.016 | 0.031 | - | 0.516 | 5108.9 | 5184.8 | 5491.9 | 5640 | 5340.2 | 5469.4 | 122 |
| 23355 | 58777 | 25.8 | 0.059 | 2 | 0.015 | 0.015 | 0.029 | - | 0.26 | 5166.7 | 5296.8 | 5600.2 | 5676.1 | 5426.4 | 5512.5 | 122 |
| 23355 | 58777 | 25.8 | 0.057 | 2 | 0.015 | 0.013 | 0.029 | - | 0.282 | 5152.3 | 5311.2 | 5585.8 | 5658 | 5426.4 | 5469.4 | 122 |
| 23355 | 58777 | 25.8 | 0.055 | 2 | 0.014 | 0.011 | 0.03 | - | 0.205 | 5177.6 | 5254.2 | 5622.1 | 5673.2 | 5469.4 | 5512.5 | 122 |
| 23355 | 58777 | 25.8 | 0.097 | 3 | 0.012 | 0.012 | 0.029 | 0.031 | 0.616 | 5108.6 | 5218.5 | 5537.8 | 5673.2 | 5340.2 | 5469.4 | 122 |
| 23355 | 58777 | 25.8 | 0.103 | 3 | 0.014 | 0.01 | 0.031 | 0.036 | 0.469 | 5195.5 | 5251.7 | 5627.2 | 5673.2 | 5469.4 | 5512.5 | 122 |
| 23355 | 58777 | 25.8 | 0.1 | 3 | 0.012 | 0.012 | 0.031 | 0.032 | 0.593 | 5134.2 | 5287.4 | 5548 | 5652.7 | 5340.2 | 5469.4 | 122 |
| 23355 | 58777 | 25.8 | 0.099 | 3 | 0.011 | 0.011 | 0.032 | 0.033 | 0.281 | 5164.8 | 5313 | 5565.9 | 5662.9 | 5340.2 | 5512.5 | 122 |
| 23355 | 58777 | 25.8 | 0.097 | 3 | 0.014 | 0.01 | 0.028 | 0.033 | 0.329 | 5169.9 | 5323.2 | 5601.6 | 5678.3 | 5383.3 | 5512.5 | 122 |
| 23355 | 58777 | 25.8 | 0.146 | 4 | 0.012 | 0.011 | 0.031 | 0.036 | 0.675 | 5162.3 | 5382 | 5624.6 | 5698.7 | 5426.4 | 5512.5 | 122 |
| 23360 | 58778 | 26.8 | 0.103 | 3 | 0.01 | 0.014 | 0.032 | 0.03 | 0.419 | 4996.5 | 5117.2 | 5440.8 | 5512.1 | 5254.1 | 5254.1 | 126 |
| 23360 | 58778 | 26.8 | 0.101 | 3 | 0.012 | 0.01 | 0.032 | 0.037 | 0.289 | 4980 | 5002 | 5484.7 | 5523.1 | 5297.2 | 5340.2 | 126 |
| 23360 | 58778 | 26.8 | 0.107 | 3 | 0.019 | 0.016 | 0.024 | 0.038 | 0.561 | 4903.3 | 4985.5 | 5435.3 | 5566.9 | 5254.1 | 5297.2 | 126 |
| 23360 | 58778 | 26.8 | 0.101 | 3 | 0.013 | 0.012 | 0.03 | 0.032 | 0.664 | 5078.8 | 5007.2 | 5586.2 | 5585.9 | 5297.2 | 5297.2 | 126 |
| 23360 | 58778 | 26.8 | 0.107 | 3 | 0.012 | 0.015 | 0.033 | 0.037 | 0.505 | 4977.1 | 4856.4 | 5426.8 | 5605.1 | 5254.1 | 5297.2 | 126 |
| 23360 | 58778 | 26.8 | 0.053 | 2 | 0.013 | 0.012 | 0.029 | - | 0.22 | 4974.3 | 5210.2 | 5498.1 | 5542 | 5297.2 | 5340.2 | 126 |
| 23360 | 58778 | 26.8 | 0.055 | 2 | 0.011 | 0.013 | 0.031 | - | 0.291 | 4971.6 | 5020.9 | 5432.3 | 5492.7 | 5254.1 | 5297.2 | 126 |
| 23360 | 58778 | 26.8 | 0.058 | 2 | 0.01 | 0.013 | 0.038 | - | 0.185 | 5158.1 | 5075.8 | 5605.1 | 5624.3 | 5383.3 | 5624.3 | 126 |
| 23360 | 58778 | 26.8 | 0.051 | 2 | 0.01 | 0.01 | 0.034 | - | 0.161 | 4999 | 5004.5 | 5509.1 | 5542 | 5297.2 | 5297.2 | 126 |
| 23360 | 58778 | 26.8 | 0.058 | 2 | 0.013 | 0.012 | 0.033 | - | 0.513 | 5086.8 | 5212.9 | 5547.5 | 5602.4 | 5297.2 | 5469.4 | 126 |
| 23356 | 58779 | 25.7 | 0.156 | 4 | 0.017 | 0.012 | 0.029 | 0.038 | 0.524 | 5280.6 | 5316.9 | 5643.8 | 5731 | 5469.4 | 5512.5 | 88 |
| 23356 | 58779 | 25.7 | 0.171 | 4 | 0.015 | 0.021 | 0.033 | 0.033 | 0.586 | 5036.2 | 5237.8 | 5572 | 5633.7 | 5340.2 | 5512.5 | 88 |
| 23356 | 58779 | 25.7 | 0.172 | 4 | 0.017 | 0.02 | 0.03 | 0.036 | 0.714 | 5014.4 | 5065.2 | 5512 | 5584.7 | 5297.2 | 5383.3 | 88 |
| 23356 | 58779 | 25.7 | 0.163 | 4 | 0.013 | 0.02 | 0.033 | 0.031 | 0.684 | 5007.1 | 5232.3 | 5461.2 | 5606.5 | 5297.2 | 5383.3 | 88 |
| 23356 | 58779 | 25.7 | 0.16 | 4 | 0.013 | 0.015 | 0.033 | 0.033 | 0.415 | 5174.2 | 5166.9 | 5552 | 5610.1 | 5340.2 | 5383.3 | 88 |
| 23356 | 58779 | 25.7 | 0.104 | 3 | 0.013 | 0.013 | 0.029 | 0.029 | 0.57 | 5060.8 | 5215.2 | 5533.1 | 5687.5 | 5533.1 | 5512.5 | 88 |

| | | | | | | | | | | | | | | | | |
|---|---|---|---|---|---|---|---|---|---|---|---|---|---|---|---|---|
| 23356 | 58779 | 25.7 | 0.11 | 3 | 0.012 | 0.015 | 0.032 | 0.031 | 0.465 | 5193.2 | 5203.5 | 5570.8 | 5673.6 | 5383.3 | 5512.5 | 88 |
| 23356 | 58779 | 25.7 | 0.114 | 3 | 0.011 | 0.019 | 0.034 | 0.032 | 0.645 | 5020.2 | 5230.9 | 5470.7 | 5594.2 | 5254.1 | 5383.3 | 88 |
| 23356 | 58779 | 25.7 | 0.118 | 3 | 0.013 | 0.02 | 0.034 | 0.034 | 0.451 | 4982.2 | 5061.9 | 5493.4 | 5578.2 | 5124.9 | 5383.3 | 88 |
| 23356 | 58779 | 25.7 | 0.116 | 3 | 0.011 | 0.021 | 0.035 | 0.034 | 0.958 | 5061.9 | 5316.2 | 5472.9 | 5611.6 | 5297.2 | 5469.4 | 88 |
| 23357 | 58780 | 26.1 | 0.106 | 3 | 0.011 | 0.013 | 0.033 | 0.035 | 0.239 | 5175.2 | 5289.5 | 5632.4 | 5798.6 | 5426.4 | 5555.6 | 143 |
| 23357 | 58780 | 26.1 | 0.107 | 3 | 0.011 | 0.019 | 0.033 | 0.031 | 0.261 | 5147.1 | 5310.8 | 5609.5 | 5788.8 | 5426.4 | 5512.5 | 143 |
| 23357 | 58780 | 26.1 | 0.103 | 3 | 0.011 | 0.014 | 0.031 | 0.038 | 0.306 | 5157.5 | 5266.6 | 5601.7 | 5679.7 | 5426.4 | 5469.4 | 143 |
| 23357 | 58780 | 26.1 | 0.099 | 3 | 0.011 | 0.011 | 0.032 | 0.036 | 0.594 | 5255.9 | 5292.6 | 5641.7 | 5737.2 | 5426.4 | 5426.4 | 143 |
| 23357 | 58780 | 26.1 | 0.109 | 3 | 0.012 | 0.019 | 0.032 | 0.029 | 0.207 | 5141.9 | 5313.4 | 5625.1 | 5755 | 5426.4 | 5555.6 | 143 |
| 23357 | 58780 | 26.1 | 0.056 | 2 | 0.011 | 0.012 | 0.033 | - | 0.22 | 5285.3 | 5307.3 | 5649 | 5733.5 | 5426.4 | 5555.6 | 143 |
| 23357 | 58780 | 26.1 | 0.057 | 2 | 0.013 | 0.013 | 0.03 | - | 0.473 | 5253.8 | 5295.4 | 5612.3 | 5729.2 | 5469.4 | 5469.4 | 143 |
| 23357 | 58780 | 26.1 | 0.045 | 2 | 0.012 | 0.012 | 0.033 | - | 0.404 | 5264.2 | 5290.2 | 5586.3 | 5630.5 | 5426.4 | 5426.4 | 143 |
| 23357 | 58780 | 26.1 | 0.055 | 2 | 0.012 | 0.012 | 0.031 | - | 0.42 | 5266.8 | 5305.8 | 5646.1 | 5752.6 | 5426.4 | 5469.4 | 143 |
| 23357 | 58780 | 26.1 | 0.056 | 2 | 0.011 | 0.013 | 0.032 | - | 0.241 | 5257.9 | 5272.6 | 5632.7 | 5713.5 | 5426.4 | 5426.4 | 143 |
| 23341 | 58781 | 26.3 | 0.089 | 3 | 0.006 | 0.009 | 0.033 | 0.032 | 0.247 | 5066 | 5406.4 | 5670.6 | 5731.6 | 5426.4 | 5555.6 | 110 |
| 23341 | 58781 | 26.3 | 0.098 | 3 | 0.009 | 0.012 | 0.032 | 0.031 | 0.257 | 5419.9 | 5361.5 | 5963.5 | 5869.5 | 5555.6 | 5869.5 | 110 |
| 23341 | 58781 | 26.3 | 0.099 | 3 | 0.011 | 0.014 | 0.03 | 0.031 | 0.348 | 5449.4 | 5402.7 | 5894.9 | 5902.1 | 5894.9 | 5598.6 | 110 |
| 23341 | 58781 | 26.3 | 0.09 | 3 | 0.008 | 0.009 | 0.03 | 0.033 | 0.286 | 5436.3 | 5429.4 | 5771.6 | 5774.3 | 5598.6 | 5598.6 | 110 |
| 23341 | 58781 | 26.3 | 0.093 | 3 | 0.008 | 0.011 | 0.031 | 0.033 | 0.319 | 5538.5 | 5509.7 | 5840.2 | 6027 | 5641.7 | 5857 | 110 |
| 23341 | 58781 | 26.3 | 0.139 | 4 | 0.007 | 0.011 | 0.033 | 0.031 | 0.379 | 5239.3 | 5493.3 | 5696.6 | 5889.6 | 5512.5 | 5727.8 | 110 |
| 23341 | 58781 | 26.3 | 0.139 | 4 | 0.01 | 0.01 | 0.03 | 0.034 | 0.284 | 5386.6 | 5488.3 | 5818.5 | 5950.6 | 5641.7 | 5727.8 | 110 |
| 23341 | 58781 | 26.3 | 0.136 | 4 | 0.009 | 0.011 | 0.03 | 0.033 | 0.335 | 5417.1 | 5495.4 | 5833.7 | 5969.6 | 5641.7 | 5770.9 | 110 |
| 23341 | 58781 | 26.3 | 0.134 | 4 | 0.007 | 0.01 | 0.031 | 0.033 | 0.389 | 5366 | 5481 | 5710.9 | 5940.8 | 5512.5 | 5727.8 | 110 |
| 23341 | 58781 | 26.3 | 0.137 | 4 | 0.009 | 0.009 | 0.029 | 0.034 | 0.385 | 5423.5 | 5560 | 5883.3 | 6005.5 | 5684.8 | 5770.9 | 110 |
| 23341 | 58781 | 26.3 | 0.051 | 2 | 0.009 | 0.01 | 0.031 | - | 0.229 | 5437.4 | 5488.3 | 5884.5 | 6026.8 | 5684.8 | 5857 | 110 |
| 23341 | 58781 | 26.3 | 0.052 | 2 | 0.01 | 0.011 | 0.032 | - | 0.245 | 5478.1 | 5528.9 | 5915 | 6037 | 5727.8 | 5857 | 110 |

| | | | | | | | | | | | | | | | | |
|---|---|---|---|---|---|---|---|---|---|---|---|---|---|---|---|---|
| 23336 | 58782 | 24.4 | 0.104 | 3 | 0.019 | 0.012 | 0.026 | 0.033 | 0.265 | 5317.7 | 5540 | 5791 | 5869.9 | 5426.4 | 5727.8 | 134 |
| 23336 | 58782 | 24.4 | 0.109 | 3 | 0.018 | 0.013 | 0.033 | 0.034 | 0.471 | 5382.1 | 5506.3 | 5701.6 | 6176.9 | 5469.4 | 5684.8 | 134 |
| 23336 | 58782 | 24.4 | 0.098 | 3 | 0.015 | 0.009 | 0.029 | 0.035 | 0.394 | 5167.4 | 5273.2 | 5576.2 | 5721.5 | 5383.3 | 5555.6 | 134 |
| 23336 | 58782 | 24.4 | 0.099 | 3 | 0.014 | 0.011 | 0.029 | 0.034 | 0.429 | 5296.8 | 5524.5 | 5698.4 | 5845.5 | 5426.4 | 5641.7 | 134 |
| 23336 | 58782 | 24.4 | 0.101 | 3 | 0.021 | 0.012 | 0.023 | 0.031 | 0.439 | 5323.7 | 5524.5 | 5782.7 | 5884.9 | 5598.6 | 5641.7 | 134 |
| 23336 | 58782 | 24.4 | 0.055 | 2 | 0.018 | 0.011 | 0.026 | - | 0.258 | 5305.4 | 5518.8 | 5513.4 | 5830.8 | 5426.4 | 5684.8 | 134 |
| 23336 | 58782 | 24.4 | 0.058 | 2 | 0.021 | 0.013 | 0.024 | - | 0.432 | 5314.4 | 5524.2 | 5787.7 | 5820 | 5426.4 | 5641.7 | 134 |
| 23336 | 58782 | 24.4 | 0.057 | 2 | 0.018 | 0.012 | 0.027 | - | 0.352 | 5305.4 | 5468.6 | 5735.7 | 5814.6 | 5426.4 | 5641.7 | 134 |
| 23336 | 58782 | 24.4 | 0.056 | 2 | 0.015 | 0.012 | 0.029 | - | 0.201 | 5037.7 | 5231.4 | 5477 | 5607.9 | 5254.1 | 5426.4 | 134 |
| 23336 | 58782 | 24.4 | 0.062 | 2 | 0.016 | 0.016 | 0.03 | - | 0.349 | 5184.8 | 5516.5 | 5685 | 5798 | 5469.4 | 5684.8 | 134 |
| 23337 | 58783 | 24.4 | 0.128 | 3 | 0.02 | 0.028 | 0.027 | 0.027 | 0.408 | 5513.4 | 5630.1 | 6040.9 | 6076.4 | 5727.8 | 5900.1 | 118 |
| 23337 | 58783 | 24.4 | 0.118 | 3 | 0.023 | 0.023 | 0.023 | 0.027 | 0.492 | 5584.4 | 5660.5 | 5972.4 | 6099.2 | 5814 | 5814 | 118 |
| 23337 | 58783 | 24.4 | 0.118 | 3 | 0.021 | 0.021 | 0.028 | 0.028 | 0.368 | 5686.7 | 5694.3 | 6031.6 | 6153.3 | 5770.9 | 5986.2 | 118 |
| 23337 | 58783 | 24.4 | 0.117 | 3 | 0.019 | 0.022 | 0.027 | 0.029 | 0.238 | 5554.9 | 5681.7 | 6018.9 | 6148.2 | 5857 | 5943.2 | 118 |
| 23337 | 58783 | 24.4 | 0.115 | 3 | 0.014 | 0.02 | 0.034 | 0.029 | 0.263 | 5509.2 | 5765.3 | 5985.9 | 6153.3 | 5814 | 5986.2 | 118 |
| 23337 | 58783 | 24.4 | 0.064 | 2 | 0.014 | 0.016 | 0.034 | - | 0.406 | 5499.1 | 5728.6 | 5965.3 | 6158.9 | 5814 | 5986.2 | 118 |
| 23337 | 58783 | 24.4 | 0.056 | 2 | 0.015 | 0.012 | 0.029 | - | 0.253 | 5298.3 | 5344.9 | 5708.9 | 5820 | 5426.4 | 5598.6 | 118 |
| 23337 | 58783 | 24.4 | 0.066 | 2 | 0.018 | 0.02 | 0.028 | - | 0.333 | 5352.1 | 5594.1 | 5823.6 | 5816.4 | 5641.7 | 5727.8 | 118 |
| 23337 | 58783 | 24.4 | 0.067 | 2 | 0.018 | 0.019 | 0.031 | - | 0.239 | 5526 | 5635.4 | 5972.4 | 6049.5 | 5814 | 5900.1 | 118 |
| 23337 | 58783 | 24.4 | 0.071 | 2 | 0.021 | 0.023 | 0.027 | - | 0.447 | 5563.6 | 5635.4 | 5968.8 | 6031.6 | 5814 | 5814 | 118 |
| 23338 | 58784 | 25.1 | 0.107 | 3 | 0.018 | 0.017 | 0.028 | 0.029 | 0.428 | 5212.3 | 5613 | 5861.5 | 5907.1 | 5727.8 | 5727.8 | 103 |
| 23338 | 58784 | 25.1 | 0.11 | 3 | 0.017 | 0.02 | 0.026 | 0.026 | 0.264 | 5252.9 | 5607.9 | 5831 | 6084.6 | 5598.6 | 5727.8 | 103 |
| 23338 | 58784 | 25.1 | 0.11 | 3 | 0.02 | 0.02 | 0.022 | 0.027 | 0.243 | 5171.7 | 5390.4 | 5713.1 | 6032.3 | 5426.4 | 5857 | 103 |
| 23338 | 58784 | 25.1 | 0.108 | 3 | 0.019 | 0.017 | 0.025 | 0.033 | 0.282 | 5131.2 | 5374.6 | 5760 | 5968 | 5426.4 | 5814 | 103 |
| 23338 | 58784 | 25.1 | 0.105 | 3 | 0.019 | 0.013 | 0.024 | 0.036 | 0.318 | 5141.3 | 5384.8 | 5800.6 | 5886.8 | 5555.6 | 5598.6 | 103 |
| 23338 | 58784 | 25.1 | 0.063 | 2 | 0.017 | 0.016 | 0.03 | - | 0.364 | 5491.3 | 5486.2 | 5897 | 5978.1 | 5727.8 | 5727.8 | 103 |

| | | | | | | | | | | | | | | | | |
|---|---|---|---|---|---|---|---|---|---|---|---|---|---|---|---|---|
| 23338 | 58784 | 25.1 | 0.065 | 2 | 0.019 | 0.018 | 0.028 | - | 0.368 | 5486.2 | 5486.2 | 5897 | 5922.3 | 5727.8 | 5727.8 | 103 |
| 23338 | 58784 | 25.1 | 0.063 | 2 | 0.017 | 0.015 | 0.032 | - | 0.419 | 5511.6 | 5501.4 | 5932.5 | 5973 | 5727.8 | 5727.8 | 103 |
| 23338 | 58784 | 25.1 | 0.065 | 2 | 0.019 | 0.017 | 0.029 | - | 0.367 | 5121.1 | 5161.6 | 5795.5 | 5831 | 5555.6 | 5555.6 | 103 |
| 23338 | 58784 | 25.1 | 0.159 | 4 | 0.017 | 0.019 | 0.026 | 0.029 | 0.355 | 5237.7 | 5607.9 | 5831 | 6059.3 | 5684.8 | 6059.3 | 103 |
| 23338 | 58784 | 25.1 | 0.158 | 4 | 0.021 | 0.02 | 0.022 | 0.029 | 0.37 | 5110.9 | 5389.8 | 5668.8 | 6013.6 | 5668.8 | 5598.6 | 103 |
| 23338 | 58784 | 25.1 | 0.152 | 4 | 0.012 | 0.01 | 0.032 | 0.042 | 0.85 | 5258 | 5359.4 | 5810.8 | 5983.2 | 5555.6 | 5684.8 | 103 |
| 23338 | 58784 | 25.1 | 0.16 | 4 | 0.015 | 0.016 | 0.03 | 0.033 | 0.356 | 5288.4 | 5501.4 | 5841.2 | 6008.5 | 5598.6 | 5814 | 103 |
| 23338 | 58784 | 25.1 | 0.164 | 4 | 0.019 | 0.019 | 0.026 | 0.03 | 0.267 | 5242.8 | 5643.4 | 5805.7 | 6018.7 | 5555.6 | 5814 | 103 |
| 23343 | 58785 | 26.6 | 0.1 | 3 | 0.009 | 0.013 | 0.032 | 0.034 | 0.391 | 5384.8 | 5499.6 | 5883.3 | 6005.2 | 5641.7 | 5814 | 110 |
| 23343 | 58785 | 26.6 | 0.099 | 3 | 0.012 | 0.009 | 0.029 | 0.034 | 0.493 | 5316.3 | 5620.5 | 6092.2 | 5995.8 | 5598.6 | 5814 | 110 |
| 23343 | 58785 | 26.6 | 0.101 | 3 | 0.01 | 0.008 | 0.038 | 0.037 | 0.332 | 5228.1 | 5611.8 | 5765.9 | 6017 | 5469.4 | 5814 | 110 |
| 23343 | 58785 | 26.6 | 0.106 | 3 | 0.012 | 0.014 | 0.032 | 0.035 | 0.85 | 5193.8 | 5698.3 | 5771.9 | 6017.8 | 5426.4 | 5814 | 110 |
| 23343 | 58785 | 26.6 | 0.099 | 3 | 0.009 | 0.011 | 0.033 | 0.032 | 0.361 | 6185 | 5202.4 | 6938 | 6020 | 6503 | 5684.8 | 110 |
| 23343 | 58785 | 26.6 | 0.059 | 2 | 0.01 | 0.017 | 0.032 | - | 0.35 | 5237.8 | 5549.8 | 5944.2 | 5958.6 | 5469.4 | 5814 | 110 |
| 23343 | 58785 | 26.6 | 0.059 | 2 | 0.014 | 0.014 | 0.032 | - | 0.291 | 6256.2 | 5338.2 | 6822.8 | 5897.6 | 6546.1 | 5641.7 | 110 |
| 23343 | 58785 | 26.6 | 0.059 | 2 | 0.011 | 0.014 | 0.034 | - | 0.26 | 5130.2 | 5273.6 | 5664.5 | 5876.1 | 5469.4 | 5469.4 | 110 |
| 23343 | 58785 | 26.6 | 0.063 | 2 | 0.017 | 0.014 | 0.032 | - | 1.048 | 4874.4 | 4982 | 5261.7 | 5283.2 | 5081.8 | 5124.9 | 110 |
| 23343 | 58785 | 26.6 | 0.059 | 2 | 0.022 | 0.018 | 0.019 | - | 0.505 | 5211.7 | 5358.8 | 5642.8 | 5774.7 | 5383.3 | 5555.6 | 110 |
| 23418 | 58786 | 26.6 | 0.061 | 2 | 0.019 | 0.02 | 0.023 | - | 0.319 | 5289.6 | 5382.8 | 5641 | 5841.8 | 5512.5 | 5512.5 | 135 |
| 23418 | 58786 | 26.6 | 0.059 | 2 | 0.018 | 0.017 | 0.025 | - | 0.302 | 5412 | 5472.9 | 5822.8 | 6000.3 | 5512.5 | 5857 | 135 |
| 23418 | 58786 | 26.6 | 0.06 | 2 | 0.017 | 0.018 | 0.025 | - | 0.297 | 6487.1 | 5589.5 | 6877.6 | 6010.4 | 6761.4 | 5857 | 135 |
| 23418 | 58786 | 26.6 | 0.062 | 2 | 0.021 | 0.017 | 0.024 | - | 0.364 | 5412 | 5594.6 | 5964.8 | 6030.7 | 5964.8 | 5900.1 | 135 |
| 23418 | 58786 | 26.6 | 0.06 | 2 | 0.017 | 0.019 | 0.024 | - | 0.26 | 5397.2 | 5612.3 | 5669.7 | 5963.8 | 5669.7 | 5770.9 | 135 |
| 23418 | 58786 | 26.6 | 0.109 | 3 | 0.016 | 0.02 | 0.026 | 0.025 | 0.524 | 5390 | 5468.9 | 5719.9 | 5992.4 | 5719.9 | 5992.4 | 135 |
| 23418 | 58786 | 26.6 | 0.101 | 3 | 0.017 | 0.016 | 0.024 | 0.028 | 0.341 | 5404.3 | 5605.2 | 5892 | 6071.3 | 5512.5 | 5857 | 135 |
| 23418 | 58786 | 26.6 | 0.109 | 3 | 0.017 | 0.019 | 0.026 | 0.029 | 0.566 | 5422.1 | 5594.6 | 5914.1 | 6010.4 | 5512.5 | 5857 | 135 |

| | | | | | | | | | | | | | | | | |
|---|---|---|---|---|---|---|---|---|---|---|---|---|---|---|---|---|
| 23418 | 58786 | 26.6 | 0.106 | 3 | 0.017 | 0.017 | 0.026 | 0.028 | 0.589 | 5375.7 | 5612.3 | 5691.2 | 6028.3 | 5512.5 | 5857 | 135 |
| 23418 | 58786 | 26.6 | 0.102 | 3 | 0.021 | 0.015 | 0.022 | 0.031 | 0.347 | 5437.4 | 5650.3 | 5980 | 6127 | 5727.8 | 5900.1 | 135 |
| 23411 | 58787 | 25.7 | 0.063 | 2 | 0.014 | 0.018 | 0.031 | - | 0.258 | 5446.5 | 5733.7 | 5974.1 | 6096.2 | 5814 | 5943.2 | 186 |
| 23411 | 58787 | 25.7 | 0.06 | 2 | 0.015 | 0.016 | 0.029 | - | 0.258 | 5514 | 5747.5 | 5950.5 | 6143.4 | 5814 | 5943.2 | 186 |
| 23411 | 58787 | 25.7 | 0.056 | 2 | 0.014 | 0.011 | 0.031 | - | 0.267 | 5417.5 | 5539.3 | 5904.8 | 6031.7 | 5641.7 | 5857 | 186 |
| 23411 | 58787 | 25.7 | 0.065 | 2 | 0.014 | 0.021 | 0.03 | - | 0.243 | 5425 | 5852.1 | 5931.1 | 6078.2 | 5770.9 | 5943.2 | 186 |
| 23411 | 58787 | 25.7 | 0.066 | 2 | 0.014 | 0.02 | 0.033 | - | 0.218 | 5514 | 5686.5 | 5925.1 | 6087.6 | 5641.7 | 5943.2 | 186 |
| 23411 | 58787 | 25.7 | 0.11 | 3 | 0.013 | 0.019 | 0.03 | 0.027 | 0.28 | 5378.3 | 5862.9 | 5841.3 | 6071 | 5641.7 | 5943.2 | 186 |
| 23411 | 58787 | 25.7 | 0.103 | 3 | 0.014 | 0.01 | 0.031 | 0.029 | 0.192 | 5407 | 5575.7 | 5909.5 | 6078.2 | 5641.7 | 5857 | 186 |
| 23411 | 58787 | 25.7 | 0.11 | 3 | 0.017 | 0.014 | 0.035 | 0.036 | 0.323 | 5453.7 | 5758.8 | 5913.1 | 6067.5 | 5684.8 | 5900.1 | 186 |
| 23411 | 58787 | 25.7 | 0.106 | 3 | 0.017 | 0.012 | 0.029 | 0.036 | 0.349 | 5600.8 | 5643.9 | 5988.5 | 6160.8 | 5857 | 5900.1 | 186 |
| 23411 | 58787 | 25.7 | 0.105 | 3 | 0.017 | 0.013 | 0.034 | 0.035 | 0.257 | 5600.3 | 5643.4 | 6120.5 | 6145.9 | 5857 | 5900.1 | 186 |
| 23436 | 58788 | 25.7 | 0.055 | 2 | 0.009 | 0.015 | 0.032 | - | 0.387 | 5212.9 | 5299.1 | 5694.7 | 5872.1 | 5426.4 | 5555.6 | 97 |
| 23436 | 58788 | 25.7 | 0.064 | 2 | 0.017 | 0.017 | 0.029 | - | 0.42 | 5649 | 5694.7 | 6252.5 | 6348.8 | 5943.2 | 6158.5 | 97 |
| 23436 | 58788 | 25.7 | 0.058 | 2 | 0.017 | 0.015 | 0.026 | - | 0.193 | 5641.6 | 6068.3 | 6071.9 | 6362.4 | 5986.2 | 6158.5 | 97 |
| 23436 | 58788 | 25.7 | 0.062 | 2 | 0.018 | 0.017 | 0.027 | - | 0.249 | 5336.8 | 5681.1 | 5598.6 | 6340.9 | 5383.3 | 5943.2 | 97 |
| 23436 | 58788 | 25.7 | 0.057 | 2 | 0.014 | 0.016 | 0.027 | - | 0.33 | 5561.6 | 5995.5 | 6106.6 | 6293.1 | 5814 | 6115.4 | 97 |

**Table S5**. Advertisement call parameters of *Allobates gasconi* sensu stricto (SS) from topotypic localities in the Jurua River, Amazonas, Brazil. Values within brackets in the first column of the table refer to the number of calls analyzed by each male. Values depict mean ± standard error and data in parentheses represent range (Pooled global). Abbreviations: FN, field number; NN, number of notes; AT, air temperature (ºC); SVL, snout–vent length; CD, call duration (ms); ND, note duration (ms), INI, inter-note interval (ms); ICI, inter-call interval (ms); LF, lower frequency (Hz); UF, upper frequency (Hz); DF, dominant frequency (Hz).

| FN | NN | AT | SVL | CD | ND | INI | ICI | LF | UF | DF |
|---|---|---|---|---|---|---|---|---|---|---|
| 23354 (5) | 2 | 26.5 | 15.9 | 0.060 ± 0.002 | 0.017 ± 0.002 | 0.025 ± 0.002 | 0.290 ± 0.090 | 5.209 ± 118 | 5.638 ± 108 | 5.467 ± 138 |
| 23354 (5) | 3 | 26.5 | 15.9 | 0.104 ± 0.002 | 0.016 ± 0.002 | 0.028 ± 0.002 | 0.028 ± 0.002 | 5.198 ± 121 | 5.595 ± 122 | 5.417 ± 129 |
| 23359 (5) | 3 | 25.8 | 17 | 0.101 ± 0.002 | 0.014 ± 0.002 | 0.029 ± 0.001 | 0.414 ± 0.089 | 5.284 ± 150 | 5.795 ± 71 | 5.529 ± 148 |
| 23359 (5) | 4 | 25.8 | 17 | 0.146 ± 0.001 | 0.015 ± 0.002 | 0.028 ± 0.003 | 0.512 ± 0.132 | 5.294 ± 163 | 5.821 ± 75 | 5.546 ± 124 |
| 23355 (5) | 2 | 25.8 | 16.4 | 0.057 ± 0.002 | 0.013 ± 0.002 | 0.030 ± 0.002 | 0.322 ± 0.120 | 5.151 ± 147 | 5.572 ± 110 | 5.404 ± 115 |
| 23355 (5) | 3 | 25.8 | 16.4 | 0.010 ± 0.002 | 0.012 ± 0.001 | 0.032 ± 0.002 | 0.458 ± 0.151 | 5.216 ± 75 | 5.622 ± 54 | 5.434 ± 75 |
| 23355 (1) | 4 | 25.8 | 16.4 | 0.146 ± 0.000 | 0.011 ± 0.001 | 0.033 ± 0.003 | 0.675 ± 0.000 | 5.272 ± 155 | 5.661 ± 52 | 5.469 ± 60 |
| 23360 (5) | 2 | 26.8 | 16.3 | 0.055 ± 0.003 | 0.012 ± 0.001 | 0.033 ± 0.003 | 0.274 ± 0.142 | 5.071 ± 93 | 5.539 ± 59 | 5.355 ± 112 |
| 23360 (5) | 3 | 26.8 | 16.3 | 0.104 ± 0.003 | 0.013 ± 0.003 | 0.033 ± 0.004 | 0.488 ± 0.142 | 4.990 ± 74 | 5.516 ± 67 | 5.284 ± 29 |
| 23356 (5) | 3 | 25.7 | 17.4 | 0.112 ± 0.005 | 0.015 ± 0.004 | 0.032 ± 0.002 | 0.618 ± 0.206 | 5.134 ± 110 | 5.568 ± 77 | 5.385 ± 130 |
| 23356 (5) | 4 | 25.7 | 17.4 | 0.164 ± 0.007 | 0.016 ± 0.003 | 0.033 ± 0.003 | 0.585 ± 0.121 | 5.153 ± 115 | 5.590 ± 74 | 5.391 ± 80 |
| 23357 (5) | 2 | 26.1 | 16.5 | 0.054 ± 0.005 | 0.012 ± 0.001 | 0.032 ± 0.001 | 0.352 ± 0.114 | 5.279 ± 19 | 5.668 ± 58 | 5.452 ± 41 |
| 23357 (5) | 3 | 26.1 | 16.5 | 0.105 ± 0.004 | 0.013 ± 0.003 | 0.033 ± 0.003 | 0.321 ± 0.156 | 5.235 ± 71 | 5.686 ± 76 | 5.465 ± 55 |
| 23341 (2) | 2 | 26.4 | 15.1 | 0.051 ± 0.001 | 0.01 ± 0.001 | 0.031 ± 0.001 | 0.237 ± 0.011 | 5.483 ± 37 | 5.965 ± 77 | 5.781 ± 88 |
| 23341 (5) | 3 | 26.4 | 15.1 | 0.094 ± 0.004 | 0.010 ± 0.002 | 0.032 ± 0.001 | 0.291 ± 0.042 | 5.401 ± 128 | 5.844 ± 109 | 5.659 ± 158 |
| 23341 (5) | 4 | 26.4 | 15.1 | 0.137 ± 0.002 | 0.009 ± 0.001 | 0.032 ± 0.002 | 0.354 ± 0.045 | 5.435 ± 90 | 5.869 ± 105 | 5.671 ± 95 |
| 23336 (5) | 2 | 24.4 | 16.5 | 0.058 ± 0.003 | 0.015 ± 0.003 | 0.027 ± 0.002 | 0.318 ± 0.090 | 5.340 ± 164 | 5.707 ± 131 | 5.508 ± 145 |
| 23336 (5) | 3 | 24.4 | 16.5 | 0.102 ± 0.004 | 0.014 ± 0.004 | 0.031 ± 0.004 | 0.399 ± 0.080 | 5.385 ± 130 | 5.804 ± 160 | 5.555 ± 121 |
| 23337 (5) | 2 | 24.4 | 16.7 | 0.065 ± 0.005 | 0.018 ± 0.003 | 0.030 ± 0.003 | 0.336 ± 0.091 | 5.517 ± 143 | 5.931 ± 136 | 5.753 ± 161 |
| 23337 (5) | 3 | 24.4 | 16.7 | 0.119 ± 0.005 | 0.021 ± 0.003 | 0.028 ± 0.003 | 0.354 ± 0.105 | 5.628 ± 85 | 6.068 ± 68 | 5.861 ± 89 |
| 23338 (4) | 2 | 25.1 | 17.1 | 0.064 ± 0.001 | 0.017 ± 0.001 | 0.030 ± 0.002 | 0.380 ± 0.026 | 5.405 ± 163 | 5.903 ± 63 | 5.684 ± 79 |

| | | | | | | | | | | |
|---|---|---|---|---|---|---|---|---|---|---|
| 23338 (5) | 3 | 25.1 | 17.1 | 0.108 ± 0.002 | 0.018 ± 0.002 | 0.028 ± 0.004 | 0.307 ± 0.073 | 5.328 ± 178 | 5.884 ± 117 | 5.646 ± 149 |
| 23338 (5) | 4 | 25.1 | 17.1 | 0.159 ± 0.004 | 0.017 ± 0.003 | 0.030 ± 0.005 | 0.440 ± 0.233 | 5.363 ± 172 | 5.904 ± 128 | 5.703 ± 155 |
| 23343 (5) | 2 | 26.1 | 16.4 | 0.060 ± 0.002 | 0.015 ± 0.003 | 0.030 ± 0.006 | 0.491 ± 0.325 | 5.321 ± 380 | 5.812 ± 435 | 5.555 ± 410 |
| 23343 (5) | 3 | 26.1 | 16.4 | 0.101 ± 0.003 | 0.011 ± 0.002 | 0.034 ± 0.003 | 0.485 ± 0.213 | 5.494 ± 305 | 6.050 ± 330 | 5.757 ± 299 |
| 23418 (5) | 2 | 26.6 | 15.4 | 0.060 ± 0.001 | 0.018 ± 0.001 | 0.024 ± 0.001 | 0.308 ± 0.038 | 5.565 ± 340 | 5.982 ± 344 | 5.831 ± 368 |
| 23418 (5) | 3 | 26.6 | 15.4 | 0.105 ± 0.004 | 0.017 ± 0.002 | 0.026 ± 0.002 | 0.473 ± 0.120 | 5.496 ± 106 | 5.942 ± 142 | 5.744 ± 178 |
| 23411 (5) | 2 | 25.7 | 16.7 | 0.062 ± 0.004 | 0.016 ± 0.003 | 0.031 ± 0.002 | 0.249 ± 0.019 | 5.587 ± 154 | 6.012 ± 85 | 5.831 ± 118 |
| 23411 (5) | 3 | 25.7 | 16.7 | 0.107 ± 0.003 | 0.015 ± 0.003 | 0.032 ± 0.003 | 0.280 ± 0.061 | 5.592 ± 151 | 6.029 ± 110 | 5.818 ± 115 |
| 23436 (5) | 2 | 25.7 | 17.1 | 0.059 ± 0.004 | 0.015 ± 0.002 | 0.028 ± 0.002 | 0.316 ± 0.094 | 5.614 ± 279 | 6.094 ± 282 | 5.848 ± 294 |
| Pooled global (142) | | (24.4-26.8) | (15.1-17.4) | 0.094 ± 0.033 (0.045-0.172) | 0.015 ± 0.004 (0.006-0.028) | 0.030 ± 0.012 (0.019-0.042) | 0.387 ± 0.154 (0.161-1.048) | 5.347 ± 234 (4.794-6.487) | 5.802 ± 233 (5.261-6.938) | 5.592 ± 235 (5.081-6.761) |

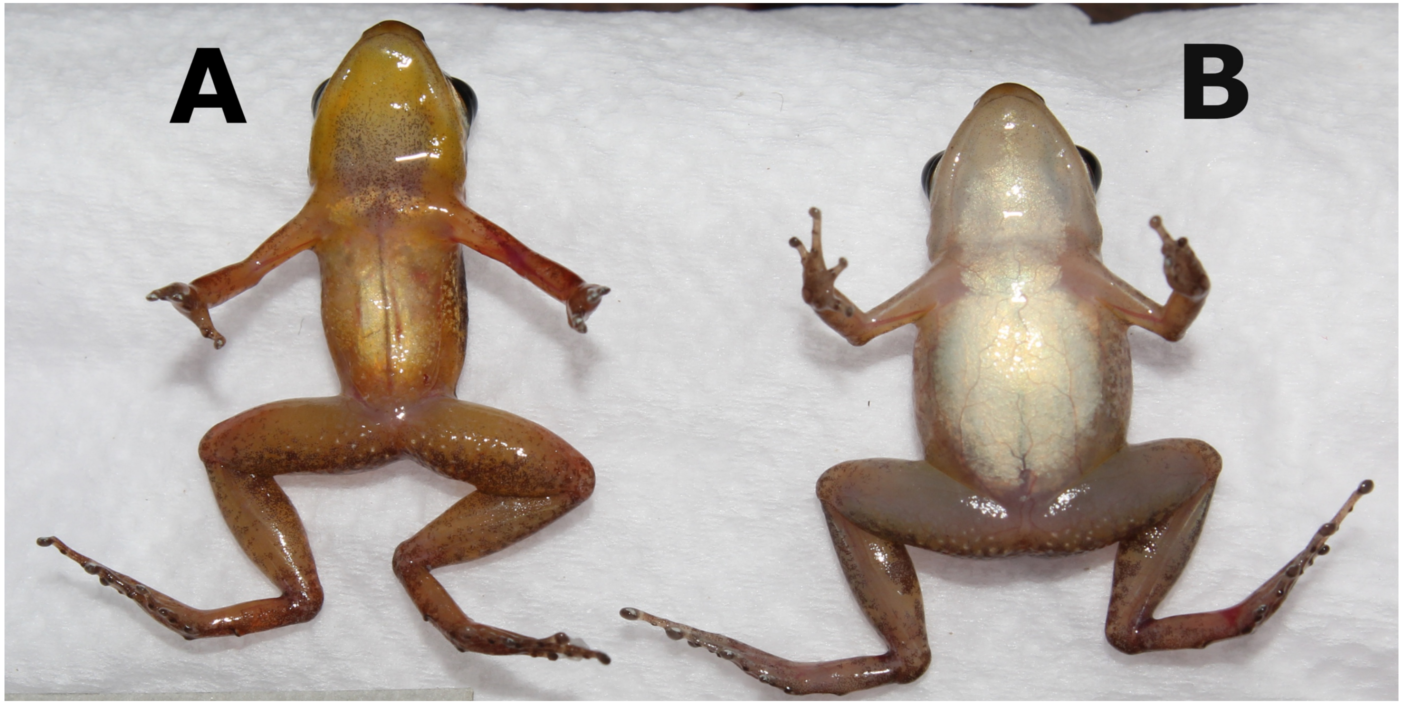


**Figure S1.** Specimens of *Allobates gasconi* SL2 from *Vai-Quem-Quer*. (**A**) ventral region of a male (APL 24069) showing a bright yellow throat and vocal sac with a few melanophores in the central and lower regions of the vocal sac; different of *A. gasconi* sensu stricto SS – which has the throat and vocal sac grey or dark grey. (**B**) ventral region of a female of *A. gasconi* SL2 (APL 24070), sowing the white throat, while the throat is light to bright yellow in female of *A. gasconi* sensu stricto SS.
